# Benchmarking Neural Decoders for Brain-Computer Interfaces and Neural Population Analysis

**DOI:** 10.64898/2026.07.21.739953

**Authors:** Joseph Soo, Kam Pang So, Xin Tang

## Abstract

The most accurate neural decoder on held-out trials is not necessarily the most useful for brain-computer interfaces or neural population analysis. In practical use, neural decoders may also need to remain robust to noisy neural inputs, satisfy calibration or deployment constraints, and produce comparable representations across recordings. We introduce BEND-BCI, an open-source benchmark of 23 neural decoding methods on motor, visual, speech and spatial decoding tasks across 16 real or synthetic neural recordings. BEND-BCI compares decoders across held-out prediction, robustness to input perturbation, computational cost and cross-recording latent consistency. These additional axes frequently changed decoder rankings: held-out accuracy did not reliably identify the most robust, efficient or cross-recording-consistent models. Simpler baselines were also competitive with, and in some cases outperformed, more heavily parameterized deep neural networks. Diagnostic analyses based on explainable machine learning further showed that decoder performance was associated with the use of expected neural features and could be improved by selecting high-quality training trials. BEND-BCI reframes neural-decoder selection from an accuracy leaderboard into a constrained decision over task, resource, representation and diagnostic goals.

---

Neural decoders are computational models that map recorded neural activity to variables of interest, including movements, sensory stimuli, task states^1^ or communication outputs^2–4^. They are central components of brain-computer interfaces (BCIs), where neural signals are translated into device commands^5,6^, and they are also widely used in systems neuroscience to relate population neural activity to behavioral and task variables^7,8^. Neural decoders are commonly compared by held-out predictive performance^1^. This convention is essential because it measures whether a trained model generalizes beyond the data used for training. However, held-out performance alone is an incomplete measure of decoder utility. In real-world BCI development, decoders may need to be robust despite unstable neural recordings^9^, recalibrated repeatedly^10^, deployed under computational constraints^11^, or adapted when current calibration data are limited^12^. In neural-population analysis, trained decoders and their learned representations may be used to compare task-related neural population structure across sessions^13–16^, subjects or conditions^17–19^. Neural decoders serve roles that extend beyond predicting target variables from neural activity.

Benchmarking decoders for these uses must compare heterogeneous decoder families without erasing their intended modes of operation. Neural decoders differ substantially in their input-output design. For example, some neural decoders map neural activity directly to the target variable^1^, some learn representations from neural activity alone and require a downstream readout for task prediction^20–23^, while others use the target variable during training to shape a representation^17,24–27^. These differences also determine which further analyses are possible. Thus, forcing every method through the same downstream readout risks obscuring what the method was designed to provide, whereas allowing every method to define its own endpoint risks losing comparability. A useful benchmark must therefore standardize final task outcomes while preserving method-faithful decoding paths.

Existing community benchmarks have already established important reference points for neural decoder evaluation^28^. The Neural Latents Benchmark provides standardized multi-dataset evaluation for latent-variable models of neural population activity^29^, and FALCON formalizes few-shot and cross-session BCI decoding^12^. The question we aim to address here is different: when multiple decoders are available, does the best held-out predictor remain the best choice under practical deployment and analytical constraints? If not, model rankings based solely on held-out predictive performance are an incomplete surrogate for neural-decoder selection.

Moreover, held-out performance does not explain what information the trained model used for prediction. A decoder can perform well by relying on neural features specific to one recording, even when those *features* are unlikely to carry task-relevant information. Conversely, a decoder can perform poorly because a small number of calibration *trials* are corrupted or mismatched to the target objective. Methods for model diagnostics from explainable learning provide useful starting points for asking these feature- and trial-level questions^30–33^. Yet these approaches have usually been applied to individual models or datasets. Their systematic application across heterogeneous neural decoders and tasks remains limited.

We introduce the <u>Be</u>nchmarking <u>N</u>eural <u>D</u>ecoders for <u>B</u>rain-<u>C</u>omputer <u>I</u>nterfaces (BEND-BCI), an open-source benchmark for selecting neural decoders beyond held-out performance (**Fig. 1**). BEND-BCI focuses on decoding from population spiking activity and brings heterogeneous decoders and datasets into a common evaluation setting. The benchmark evaluates four decoder-selection measurements: task prediction, robustness to noisy neural inputs, computational cost, and latent representation consistency. The results reveal that models leading in prediction accuracy often lose their advantage in robustness, resource efficiency or latent consistency. BEND-BCI also includes two model diagnostics, feature attribution and trial valuation, that ask whether a decoder relies on expected input features and which training trials help or hurt its performance. A freely available interactive online resource provides code, data manifests, benchmark outputs and visual summaries to support decoder selection and future benchmarking of new decoders and data (https://tang-lab-benchdash.hf.space/).

**Fig. 1.**
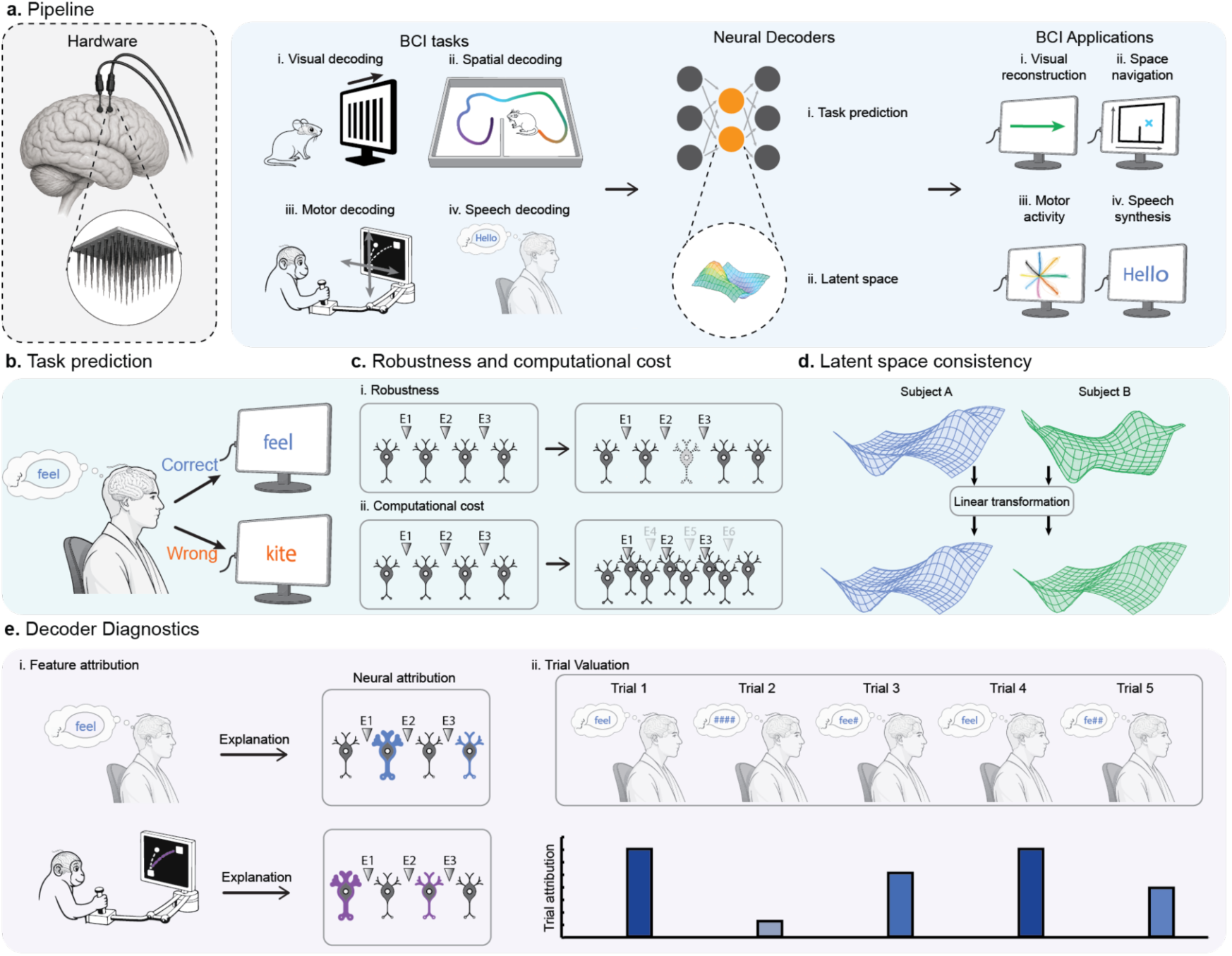
BEND-BCI evaluates neural decoders across selection measurements and diagnostic assays. **a,** Benchmark workflow. Biological recordings and synthetic simulations are converted to trial × time × feature arrays and paired with dataset-specific targets for visual-orientation classification, two-dimensional position or force regression, attempted-word classification and spatial-position regression. Evaluated decoder workflows yield task-specific predictions (i), either natively or through a downstream readout. Some methods also expose latent representations of neural population activity (ii); the orange nodes and enlarged view illustrate one such representation. Predictions are scored against held-out targets. **b,** Held-out task prediction is quantified by *R*^2^ for regression and accuracy for classification. **c,** Robustness and computational cost. Held-out neural inputs are perturbed with additive Poisson count noise to assess robustness (**i**); runtime and memory use during training and inference quantify computational cost (**ii**). **d,** Cross-recording latent alignability is assessed by matching task-defined landmarks across recordings, participants or independently generated simulations, fitting a linear alignment and scoring landmark agreement after alignment. **e,** Diagnostic assays estimate Shapley contributions of neural input features to the task score (**i**) and model-specific values of candidate training trials for a prespecified downstream decoding utility (**ii**).

## Results

### BEND-BCI defines a multi-axis benchmark for neural decoder selection and diagnostics

BEND-BCI organizes heterogeneous neural-decoding tasks into a common evaluation pipeline (**Fig. 1a**). The neural activity and the paired stimuli or behavior labels are organized into a standard data format (see **Methods**). The five primary benchmark tasks comprised two-dimensional hand-position decoding from macaque center-out reaching^34^, visual-orientation classification from the Allen Brain Observatory Visual Coding dataset^35^, attempted-word classification from intracortical speech recordings^36^, isometric-force decoding from MC PacMan^37^, and spatial-position decoding from synthetic RatInABox navigation data^38^. Different neural decoders then take the data to produce task-specific predictions directly or through a readout, and each prediction is scored against the target for that dataset (**Supplementary Table 1**).

We designed the benchmark to comprehensively evaluate neural decoders and perform standardized diagnostics that inform practical deployment or analysis decisions (**Fig. 1b-e**; see **Methods**). *Task prediction* asks how precisely a trained decoder produces the held-out output. *Robustness* asks whether the prediction score is preserved when neural inputs are perturbed with additive count noise, and *computational cost* measures training time, inference time, peak RAM and peak GPU memory. *Latent consistency* asks whether representations of the same task-defined states align across independent simulations, biological subjects^19^, or recording sessions. *Feature attribution* and *trial valuation* use Shapley-value-based procedures^39^ to estimate which neural features^30^ and training trials^32^ contribute to the task score. BEND-BCI reports these measurements and assays separately so that decoder choice can be conditioned on the intended goals; unsupported analyses are reported as missing coverage (**Supplementary Table 2**).

### Prediction, robustness and resource constraints favor different decoders

No decoder dominated held-out task prediction across five primary tasks (**Fig. 2a**). In the benchmark matrix, prediction, robustness to noisy neural inputs and resource cost are displayed as separate blocks to show whether the same trained decoder remained preferred across objectives. XGBoost^40^ led Allen visual coding (accuracy = 0.91), SVM^1^ led attempted speech (accuracy = 0.97), NEDS^41^ led MC PacMan force decoding (*R*² = 0.98), and NEDS-pretrained (NEDS-pt)^41^ led macaque reaching and RatInABox navigation (*R*² = 0.97 and 0.998, respectively). Thus, simpler decoders such as XGBoost and SVM were competitive on the trial-level classification tasks, whereas more complex models such as NEDS led several continuous trajectory tasks. Aggregating each method’s within-task ranks across the five tasks (see **Methods**) identified NEDS^41^ and NEDS-pt^41^, the from-scratch and cross-session-pretrained versions of the same model family, as the strongest task prediction choices, with MINT^42^, GRU^1^, LDNS^22^, DPAD^25^ and XGBoost^40^ forming the next group. Representative held-out predictions illustrate the scored outputs for continuous trajectories and discrete class labels (**Fig. 2b**).

**Fig. 2.**
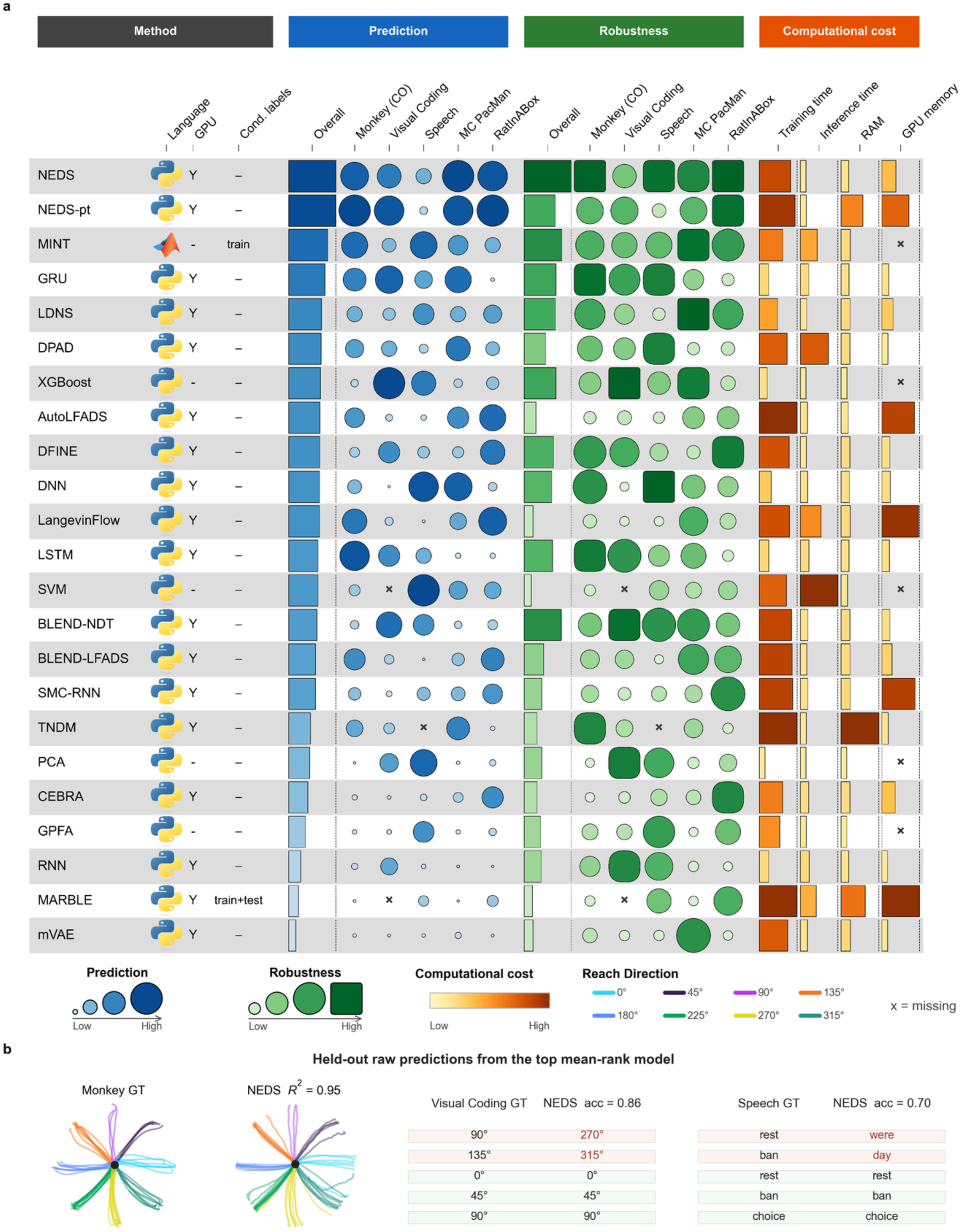
Prediction performance, robustness, and computational cost identify different decoder choices. **a,** Benchmark matrix for 23 decoder configurations across five primary tasks. Rows are ordered by mean within-task prediction rank; missing eligible entries receive the worst within-task rank for this aggregate. The Overall bars summarize each decoder’s average rank across the five tasks; longer bars indicate better average rank. The blue block reports held-out prediction, quantified by *R*^2^ for regression and accuracy for classification. The green block reports robustness as the raw area under the task-score-versus-noise curve after additive Poisson perturbations at *λ* = 0, 0.2, 0.4, 0.6, and 0.8. Because this summary uses the raw task score, it reflects both unperturbed performance and score retention under perturbation and is not normalized to the *λ* = 0 value. Within each displayed column, larger and darker symbols denote higher values. The orange block reports training time, inference time, peak RAM and peak GPU memory for the macaque-reaching comparison; greater orange extent and intensity denote greater resource cost. The left columns indicate implementation language, GPU use, and additional information used during fitting or representation construction. “Train” for MINT denotes training labels used to index neural and target trajectory libraries; RatInABox used no-condition construction. “Train+test” for MARBLE denotes condition labels from both splits used, where applicable, to group neural trajectories expected to share dynamics. Test targets were used only for scoring. Crosses denote unavailable entries; CPU-only methods have no GPU-memory value. **b,** Example held-out outputs from NEDS, the top mean-rank neural decoder across the five tasks. Panels show macaque hand trajectories and five held-out trials each from Allen visual coding and attempted speech. For speech, rest denotes the “Do Nothing” cue. Trajectory colors denote reach direction; green and red rows denote correct and incorrect class predictions, respectively.

Decoder rankings remained task dependent under alternative scoring metrics, including balanced accuracy, macro and weighted F1, macro precision and macro recall for classification, and RMSE, MAE, Pearson’s *r* and Spearman’s *r* for regression (**Supplementary Figs. 1–3**). Adding spike-band-power-derived features reduced speech accuracy on average (mean Δ accuracy = −0.033, see **Methods**), while preserving a broadly similar method ordering (Spearman’s *r* = 0.88, *n* = 22; **Supplementary Fig. 4**).

Decoders with the highest unperturbed scores were not always the most robust to noisy neural inputs (**Fig. 2a**). We evaluated each trained model on test arrays perturbed with additive Poisson count noise while leaving the training data, task targets and trained model unchanged (see **Methods**). Robustness was summarized as the raw area under the score-versus-noise curve, which captures both baseline score and score retention under added count noise. In MC PacMan force decoding, NEDS^41^ had the highest unperturbed *R*² but LDNS^22^ had the largest raw area: NEDS fell from *R*² = 0.98 to 0.35 at the largest perturbation, whereas LDNS fell from 0.87 to 0.72. In speech, SVM^1^ started with the highest accuracy (0.97), but DNN^1^ had the larger raw area under the perturbation curve (0.3 versus 0.188 for SVM), retaining 0.61 accuracy at the first noise level (0.2) compared with 0.3 for SVM. These reversals show that held-out prediction was informative but did not fully determine robustness to noisy neural inputs (**Supplementary Fig. 5**). Dataset-specific robustness examples are shown in **Supplementary Figs. 6–10**.

Resource constraints changed the top-ranked decoder set again (**Fig. 2a** and **Supplementary Figs. 11 and 12**). On the shared macaque-reaching resource comparison, training time ranged from 30 ms to about 39 min, inference time from 1.6 ms to 13 s and peak RAM from 0.27 GB to 14.8 GB. PCA^1,43^ and XGBoost^40^ defined the low-cost end of the comparison. NEDS combined strong mean prediction rank with millisecond-scale inference time but required substantially longer training time than these lightweight baselines. Pairing prediction rank with measured cost produced budget-dependent choices: sub-second training budgets favored PCA^1,43^ and XGBoost^40^, budgets from about 2 s to 1 min favored GRU^1^, budgets of roughly 1–8 min favored MINT^42^, and NEDS^41^ became the top mean-rank option beyond approximately 8 min (see **Methods**). The decoder favored for frequent retraining was therefore not necessarily the decoder favored when training could be amortized and inference latency was the limiting constraint.

### Latent-consistency leaders differ across tasks and from prediction leaders

A model’s predictive accuracy within one recording does not establish whether its learned representation can be compared across recordings. Cross-session, cross-subject and cross-condition analyses require a stronger property: latent consistency. The neural representation of the same task state should occupy a comparable region of the latent space after a post hoc linear alignment. This latent consistency is not guaranteed, because independently trained latent models can decode well within one recording while differing by distortions or recording-specific dimensions introduced by neuron sampling and recording instability^9,44^. Following recent methods such as CEBRA and MARBLE, which explicitly target consistent embeddings across recordings^17,18^, we defined matched task landmarks as points or trajectories in each model’s representation space that corresponded to the same task state. These landmarks were reach trajectories in macaque reaching^34^, stimulus- or word-class centroids in Allen visual coding^35^ and speech^36^, and spatial-bin centroids in RatInABox^38^. We measured latent consistency by fitting a simple linear alignment between matched task landmarks and scoring how well the aligned landmarks agreed across recordings (**Fig. 3a** and **Supplementary Figs. 13–16**; see **Methods**).

**Fig. 3.**
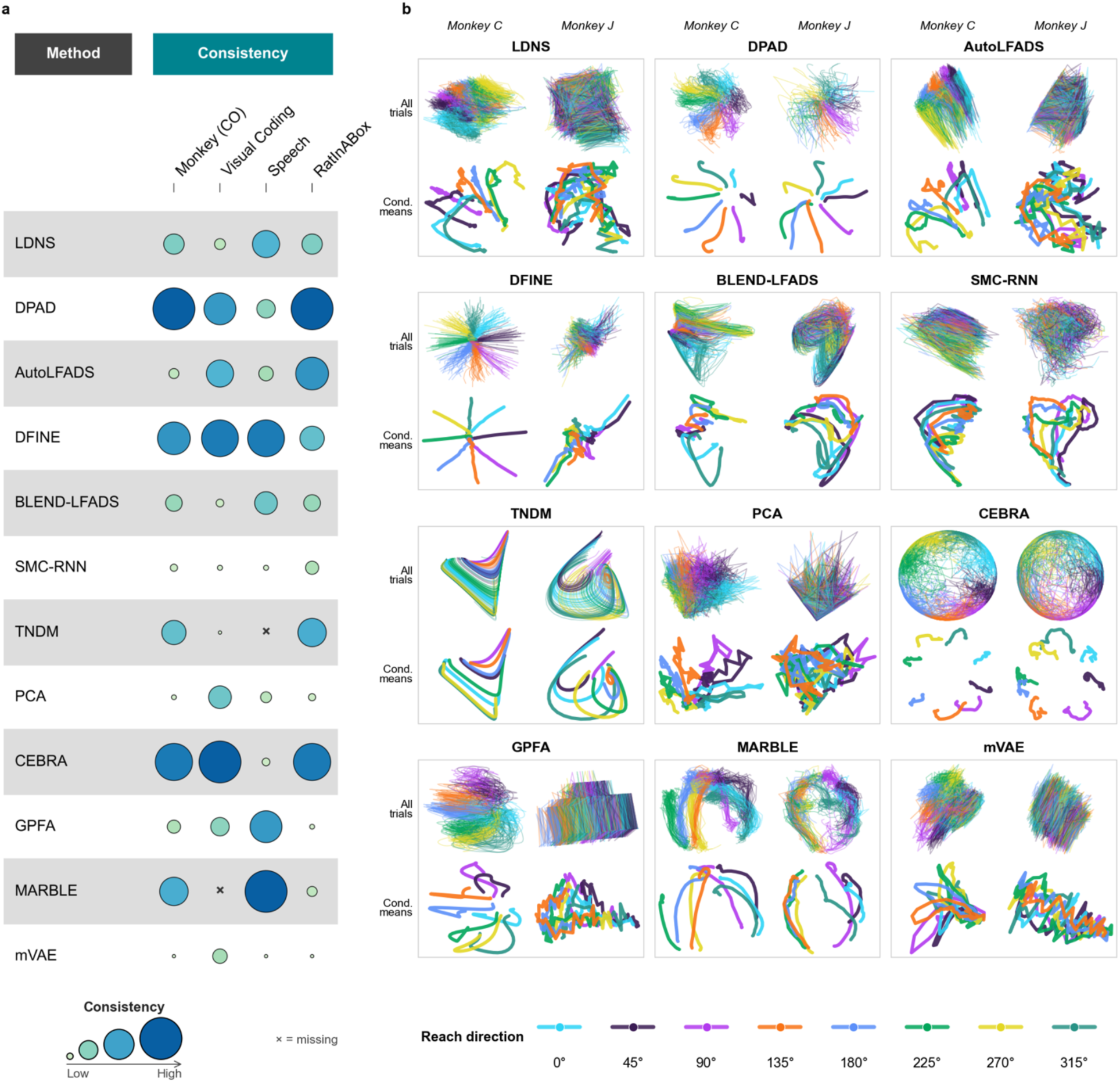
Cross-recording latent consistency of matched task landmarks varies across methods and tasks. **a,** Alignment summary for 12 methods configured to produce three-dimensional representations on macaque reaching, Allen visual coding, attempted speech and RatInABox navigation. Representations were centered and whitened within each recording, and matched task landmarks were aligned using an intercept-free linear map in both directions. The reported symmetric alignment score is the mean of the two directional *R*² values. Bubble size and shade encode within-task rank; crosses denote unavailable entries. The score quantifies linear alignability of the latent space through landmarks and does not measure cross-recording decoder transfer or identification of a unique latent coordinate system. **b,** Macaque-reaching representations for monkeys C and J. The upper row for each method shows individual trial trajectories, and the lower row shows reach-direction means. Colors denote reach direction. Three-dimensional representations are displayed in a common two-dimensional view. For visualization only, monkey J was Procrustes-aligned to monkey C within each method; this display alignment was not used to compute the scores in **a**.

The most consistent method depended on the task (**Fig. 3a**), and the task-specific leaders matched different representation-use cases. CEBRA^17^ was most consistent for the Allen visual-coding data (*R*² = 0.998), whereas DPAD^25^ led the macaque center-out reaching (*R*² = 0.856) and RatInABox spatial decoding data (*R*² = 0.977). Attempted speech had a compressed top group led by MARBLE^18^, DFINE^27^, GPFA^20^, LDNS^22^, BLEND-LFADS^24^ and DPAD^25^ (*R*² = 0.51–0.6). These task-specific leaders differed from the prediction leaders, underscoring that latent consistency was not a restatement of held-out prediction. Within-recording prediction accuracy and cross-recording latent consistency were only weakly related in the cross-analysis summary: a strong within-recording predictor did not guarantee a representation that would align well across sessions.

Latent-space visualizations revealed whether decoders preserved comparable structure across recordings. In macaque center-out reaching, high-consistency methods kept the eight reach-direction trajectories separated and in a similar relative arrangement, whereas lower-consistency methods collapsed reach directions or changed the landmark arrangement under the same alignment (**Fig. 3b**). Similar examples in Allen visual coding, attempted speech and RatInABox used class or spatial centroids to show that neural representations retained their relative arrangement (**Supplementary Figs. 13–16**).

We asked how far the held-out prediction rank carries to these other axes, examining its agreement with robustness to noisy neural inputs, cross-recording latent consistency and feature-attribution validation (**Supplementary Fig. 17**). To compare decoders across datasets without conflating task difficulty or metric scale, we ranked decoders within each dataset before pooling model-dataset entries (see **Methods**). Prediction rank correlated with robustness to noisy neural inputs (Spearman’s *r* = 0.64, *P* < 0.001, *n* = 112) and showed a smaller positive association with latent consistency (Spearman’s *r* = 0.3, *P* = 0.041, *n* = 46). The results showed that held-out prediction rank incompletely captures the other benchmark axes.

### Feature attribution as a diagnostic for whether decoder performance is grounded in expected inputs

An ideal neural decoder’s performance should rest on neural signals that encode the true target variables of interest. If a decoder instead relies on idiosyncratic or irrelevant input features, a high prediction score can be misleading, and the source of low or unstable performance can be hard to diagnose from the score alone. We therefore used feature attribution to probe decoder behavior: when a decoder succeeded or failed, did benchmark utility concentrate on inputs expected to carry task information or on features not expected to support the task? BEND-BCI estimated these contributions with global Kernel SHAP^30^ (**Fig. 4**; see **Methods**). We then compared estimated feature contributions with expected feature importance to ask whether decoders learned to rely on the expected signal or on an unreliable shortcut (**Fig. 4a**).

**Fig. 4.**
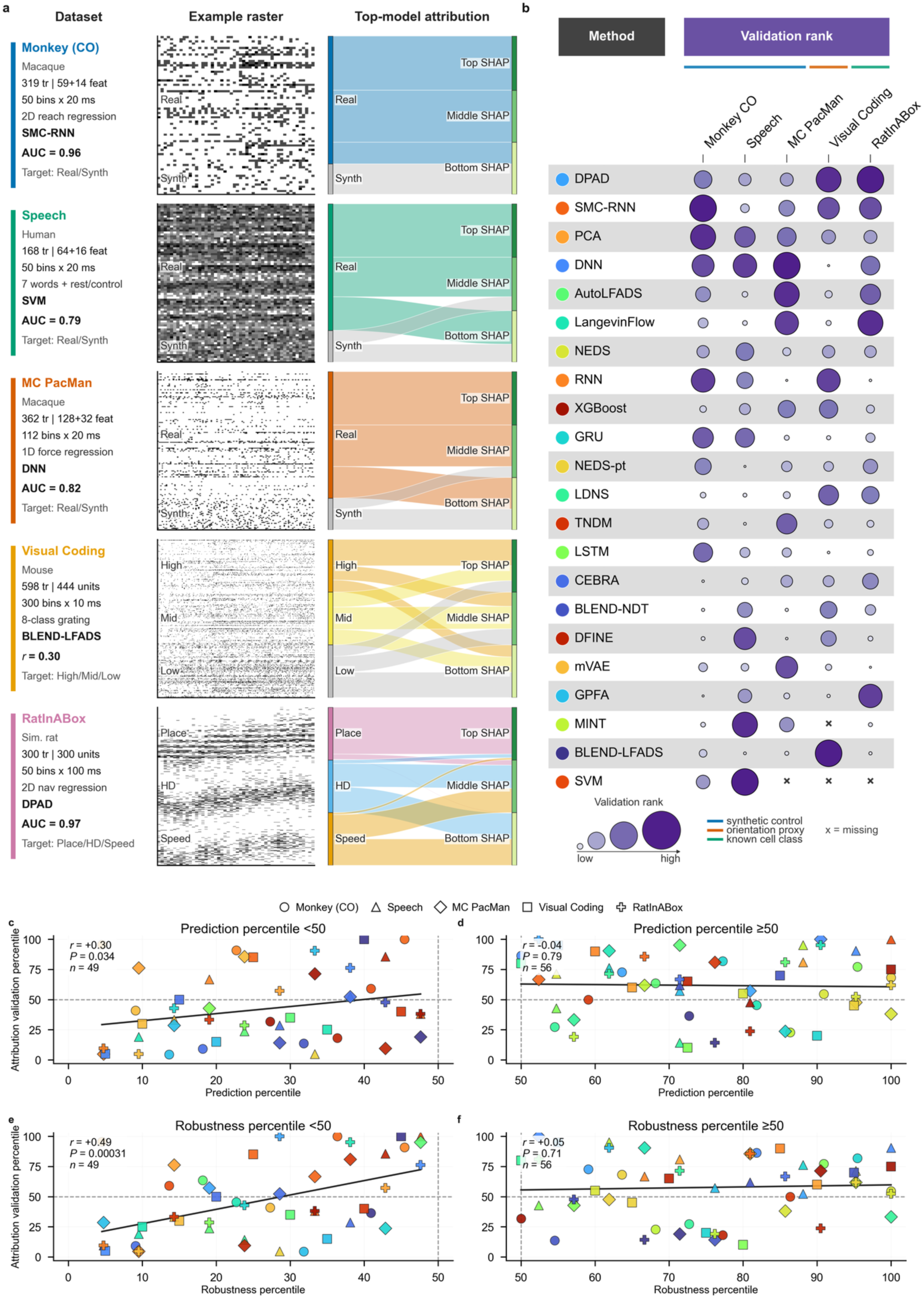
Feature-attribution analysis shows that lower-performing decoders rely less on the expected signal. **a,** Dataset-level examples. Each row summarizes the neural input and validation target, shows an example feature raster and maps feature groups to the upper, middle and lower thirds of the signed global Kernel SHAP ranking for the model with the highest validation score in that dataset. Ribbon width is proportional to the number of features assigned to each rank bin. These selected models are shown for illustration. **b,** Feature-attribution validation across evaluated models. For macaque reaching, attempted speech and MC PacMan, ROC-AUC measures whether recorded neural features rank above appended synthetic controls. For Allen visual coding, Spearman’s *r* quantifies alignment with measured neural tuning. For RatInABox, ROC-AUC measures whether place cells rank above head-direction and speed cells. Chance performance is 0.5 for ROC-AUC; larger positive correlations indicate stronger agreement with the Allen proxy. Bubble size and shade encode within-dataset validation rank; crosses denote unavailable entries. **c,d,** Prediction percentile versus attribution-validation percentile below (**c**) and at or above (**d**) the median prediction percentile. **e,f,** Corresponding relationships for robustness percentile. Each point is a model-dataset entry after within-dataset percentile normalization; marker shape denotes dataset and color denotes method. Dashed lines mark the 50th percentile. Solid lines are linear fits for visualization; annotations report Spearman’s *r*, the two-sided *P* value and the number of model-dataset entries. Associations of prediction and robustness with agreement between feature attributions and expected task-relevant inputs were concentrated among lower-performing decoders.

Feature-attribution validation recovered expected feature structure across constructed controls, simulated neuron classes and measured neural tuning (**Fig. 4b** and **Supplementary Figs. 18–20**). In the synthetic-control analysis, we appended features with no task information and asked whether SHAP values ranked recorded inputs as more important than those controls. Recorded inputs ranked above synthetic controls most clearly in macaque reaching (mean ROC-AUC, 0.83; chance, 0.5), with weaker separation in MC PacMan force decoding (0.66) and speech (0.63). RatInABox provided a positive-control setting with known simulated neuron classes^38^: decoders predicted two-dimensional position from place, head-direction and speed cells (**Supplementary Fig. 21**), and SHAP values ranked place cells above the other neuron classes (mean ROC-AUC, 0.79), with DPAD^25^ and LangevinFlow^23^ leading. Allen visual coding provided a complementary biological validation: we compared feature attribution with each unit’s independently measured drifting-gratings orientation selectivity^35^, an external tuning property relevant to orientation classification. Agreement with this external tuning measure was generally positive (mean Spearman’s *r* = 0.14), with BLEND-LFADS^24^ and DPAD^25^ showing the strongest agreement.

More accurate decoders tended to agree better with the feature-validation targets (Spearman’s *r* = 0.350, *P* = 0.00025, *n* = 105; **Fig. 4c, d**). Attribution validation increased with prediction below the median prediction percentile (Spearman’s *r* = 0.304, *P* = 0.034, *n* = 49), while the corresponding association was near zero at or above the median (Spearman’s *r* = −0.037, *P* = 0.787, *n* = 56). Robustness percentile showed the same pattern (**Fig. 4e, f**): attribution validation increased below the median robustness percentile (Spearman’s *r* = 0.494, *P* = 0.00031, *n* = 49), while the corresponding association was near zero at or above the median (Spearman’s *r* = 0.052, *P* = 0.706, *n* = 56). Together, these results establish feature attribution as a diagnostic of decoder failure modes. The relationship was concentrated in the lower-to-moderate performance range, where weak or fragile performance more often coincided with poorer agreement between attributed utility and expected task-relevant feature classes (**Fig. 4c–f**).

### Trial valuation as a diagnostic for which training trials support or degrade decoder performance

Neural recordings are noisy and unstable, with signal quality varying across channels, units, time periods and behavioral trials^9,44^. As a result, not every training trial is equally useful for calibrating a decoder: some trials may reflect corrupted signals, atypical behavior or recording-specific noise. A complementary diagnostic asks whether and why a data-selection intervention helps or fails for a fitted decoder: which candidate trials support the decoding utility, and which degrade it. We used Data Shapley^32^, a training-data importance attribution method, to assign each candidate training trial a marginal contribution to held-out decoding performance. Because the value is defined relative to a specific model output, readout, candidate pool, valuation split and performance metric, we used trial valuation as a decoder-specific diagnostic of two defined interventions: removing synthetically corrupted trials and selecting older same-subject trials for target-session decoding (see **Methods**).

We first tested this diagnostic in a controlled assay with known corrupted examples (**Fig. 5a**). Specifically, to create this controlled corruption, within each dataset we rotated approximately one third of the training trials by 75 degrees in population-activity space while leaving the benchmark target unchanged (see **Methods**), so that the rotated trials were known to be corrupted. If trial values capture trial utility, rotated trials should receive lower values than unrotated trials. Lower trial values separated rotated from unrotated trials most clearly for MC PacMan force decoding (mean ROC-AUC = 0.794), RatInABox navigation (0.790) and macaque center-out reaching (0.748), with weaker separation for Allen visual coding (0.608) and speech (0.562) (**Supplementary Figs. 22–26**).

**Fig. 5.**
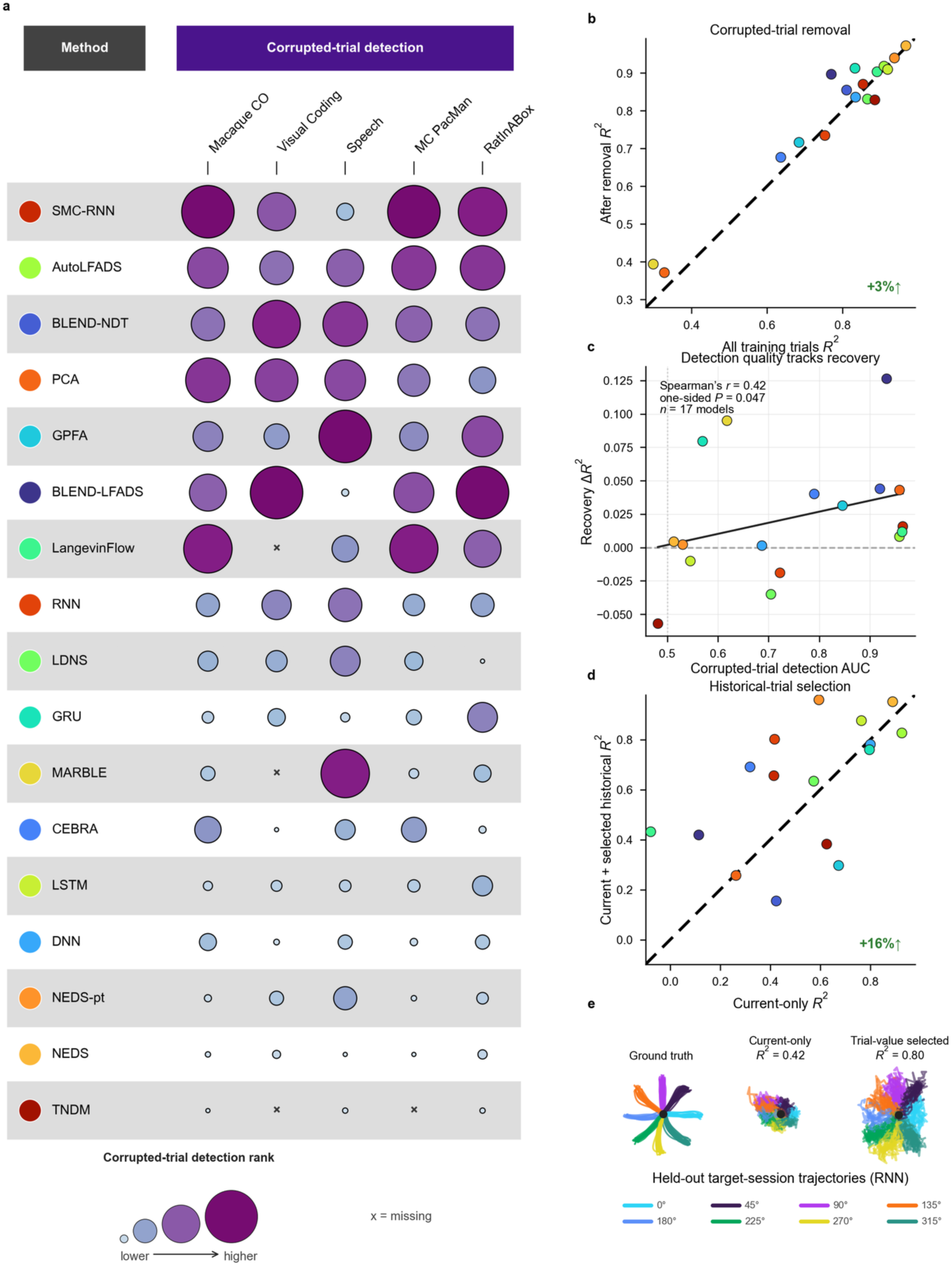
Trial valuation identifies training trials that support or degrade decoder performance. **a,** Controlled corrupted-trial detection. One third of the training trials were perturbed by a 75° rotation in population-activity space while the corresponding targets were left unchanged. ROC-AUC was calculated using negative trial value as the detection score, with larger values indicating better separation of perturbed and unperturbed trials. Bubble size and color encode within-dataset detection rank; crosses indicate unavailable results. **b,** Trial-value-guided removal of corrupted macaque-reaching trials. Each point represents one decoder trained on a mixture of unperturbed and perturbed trials. The *x* axis shows test-set *R*² before filtering, and the *y* axis shows test-set *R*² after removing trials with negative values and retraining the decoder. The dashed line indicates equal performance. **c,** Relationship between corrupted-trial detection ROC-AUC and recovery in decoding performance after trial removal. Recovery was calculated as post-removal *R*² minus pre-removal *R*². The annotation reports Spearman’s *r*, one-sided permutation *P* value and the number of evaluated models; the solid line shows a linear fit for visualization. **d,** Historical-trial selection for same-subject decoding. Current-session training trials were retained, and trial values were assigned to candidate trials from earlier sessions represented in a shared M1 channel space. The *x* axis shows held-out target-session *R*² using current-session training data alone, and the *y* axis shows *R*² after adding historical trials with non-negative trial values. The dashed line indicates equal performance. In b and d, green annotations report the percentage change in mean *R*² relative to the *x*-axis condition. **e,** Representative held-out target-session trajectories for the RNN trained with current-session trials alone or with trial-value-selected historical trials. Colors denote reach direction.

We next asked whether corrupted-trial detection quality tracked recovery after trial removal. In the macaque-reaching data, we trained the models on a mixture of unperturbed trials and trials corrupted by the same neural-feature rotation described above, and then retrained them after removing trials with negative trial value (**Fig. 5b,c**). Trial-value-guided removal improved test-set *R*² in 13 of 17 evaluated models and increased mean *R*² by 3%. Across models, corrupted-trial detection ROC-AUC was positively associated with recovery after removal (Spearman’s *r* = 0.42, *P* = 0.047, *n* = 17; **Fig. 5c**), consistent with lower trial values diagnosing examples whose inclusion degraded the defined benchmark objective.

Neural recordings may drift across sessions^9^, and continually recalibrating a decoder on freshly collected data is burdensome and not always feasible; this has motivated approaches that reduce dependence on fresh calibration each session by reusing neural representations across prior recordings^9,10,14,15,45^. We therefore asked whether trial valuation could identify useful historical trials for target-session decoding when current-session calibration data were limited. To keep this test focused on within-subject session drift, we performed a same-subject macaque-reaching case study (**Fig. 5d, e** and **Supplementary Fig. 27**). Trial-value selection raised mean held-out *R*² to 0.617, above both current-only training (*R*² *=* 0.532) and all-session pooling (*R*² *=* 0.546), and the median showed the same ordering (*R*² *=* 0.673 versus 0.584 and 0.615).

## Discussion

BEND-BCI shows that neural-decoder benchmarking should treat prediction accuracy as the starting point for model selection, not the endpoint. A high held-out score establishes that a decoder solves a particular benchmark task under a specified split. It does not establish that the same model is robust to input perturbation, feasible under calibration and deployment budgets, or suitable for representation comparison. Held-out performance alone cannot diagnose the model’s failure modes. In our benchmark, held-out prediction rank was associated with robustness to noisy neural inputs and feature-attribution validation, but was a weaker guide to latent consistency. The practical question is not which decoder is best in the abstract, but which decoder satisfies the constraints imposed by a particular BCI or neural-population analysis.

This framing turns decoder benchmarking into a use-case-specific decision problem. For different neural decoding tasks, users should inspect task-specific scores because the benchmark did not support a universal winner. For frequent recalibration, measured training time and memory become part of model choice. For amortized use, fast inference can make a higher-training-cost decoder feasible. For cross-session or cross-subject representation analysis, latent consistency can select methods different from the most accurate within-recording predictor. For model inspection, feature attribution and trial-level data valuation provide diagnostics for asking whether a decoder relies on expected input features and which training trials help or hurt performance.

These use cases imply a practical design principle for neural-decoder builders: architectural complexity should be justified by improved performance on an evaluation dimension. In BEND-BCI, more complex models did not uniformly dominate simpler alternatives. Lightweight baselines such as PCA^1,43^ and XGBoost^40^ were competitive on several tasks and occupied the low-cost end of the benchmark. More sophisticated neural decoders were most compelling when their added complexity served a specific goal. Decoder development should therefore begin with strong simple baselines and then add architectural complexity. This principle is particularly relevant for plug-and-play decoders^46^ and large-scale pretrained neural models^41,47^. Their value should be assessed by measurable gains in calibration efficiency or cross-recording representation consistency, not by model scale alone.

The diagnostics in BEND-BCI should be interpreted with care^48^. Feature attribution and trial-level data valuation are validation-checked diagnostics of trained decoder behavior, not causal explanations of neural computation or universal labels of feature or trial quality. Their values depend on the trained decoder, masking or valuation design, data split, candidate pool and performance metric^30,32^. Future work should test whether these diagnostic patterns are stable across alternative attribution^49^ and valuation methods^50^ and whether diagnostic-guided interventions improve performance on independent future sessions or online BCI use.

Several extensions can broaden BEND-BCI. The controlled count-noise assay quantifies sensitivity to neural-count perturbations and can be expanded to biological drift, electrode instability and longitudinal nonstationarity. Runtime and memory measurements provide a reference under the benchmark execution environment and can be repeated on additional hardware and embedded platforms. The current benchmark spans four major neural-decoding task families; additional species, recording modalities, behavioral outputs, closed-loop settings and longitudinal timescales remain opportunities for expansion. Analysis availability follows the outputs exposed by each decoder. The open-source benchmark and interactive resource support these extensions with new datasets, perturbations, tasks and decoder families.

## Methods

### Benchmark design

We designed the benchmark to compare neural decoding models under a shared evaluation framework while preserving the target variable and temporal structure of each dataset. We represented neural activity as trials by time bins by input features. Depending on the dataset, we used either a trial-level class label or a time-varying continuous trajectory as the target. We used the same train/test split for the main prediction, robustness, feature-attribution and trial-valuation analyses, except in the trial-valuation retraining case studies.

We evaluated 23 neural decoding methods and model variants on five datasets and organized the primary analyses around four decoder-selection measurements and two diagnostic assays. The selection measurements were task prediction, robustness to added count noise, computational cost and cross-recording latent consistency; the diagnostic assays were feature attribution and trial-level data valuation.

We evaluated primary prediction, robustness and computational cost broadly across method-dataset combinations that supported the relevant task. We evaluated feature attribution across eligible method-dataset combinations using dataset-specific validation targets and evaluated trial-level data valuation when fitted outputs or representations could be rescored without retraining the full model. **Supplementary Notes 1–7** provide additional details on model integration, task adaptations, preprocessing, label use, feature attribution and trial valuation.

### Datasets

We selected five datasets to cover four BCI-relevant task families: motor decoding, visual decoding, speech decoding and spatial decoding (**Supplementary Table 1**). We chose the datasets to include regression and classification targets, single-recording and multi-recording analyses, biological and synthetic validation signals, and both spike-sorted and channel-level feature definitions.

The macaque reaching prediction dataset used DANDI 000688 version 0.250122.1735 session sub-C_ses-CO-20151104_behavior+ecephys^34^. The main task was regression from spiking activity to two-dimensional hand position. After benchmark preprocessing, the prediction array contained 319 successful trials, each represented by 50 bins of spike counts from 59 spike-sorted units. We aligned trials to the go cue, binned spikes at 20 ms resolution and included 300 ms of pre-go activity followed by 700 ms of post-go activity. To account for the physiological delay between cortical activity and movement, we sampled hand-position targets 150 ms after the corresponding neural bin. This fixed lag is consistent with the range reported by Wu et al., who found that 140–150 ms lags produced the most accurate Kalman decoding of hand kinematics from motor-cortical firing activity in their experiments^51^. The historical-selection case study used a separate channel-level M1 representation from the same subject across different sessions. The representation is defined below for trial-value-guided retraining.

The Allen Neuropixels dataset used drifting-gratings session 721123822 from the Allen visual coding Neuropixels release^35^. Consistency analysis additionally used sessions 715093703 and 732592105. Sessions 721123822, 715093703 and 732592105 were recorded from a Pvalb-IRES-Cre × Ai32 mouse, an Sst-IRES-Cre × Ai32 mouse and a wild-type mouse, respectively. The task was eight-way orientation classification. The main prediction dataset contained 598 drifting-grating presentations, 300 bins per trial, and 444 units. We binned neural activity at 10 ms resolution, with 1 s before stimulus onset for history and 2 s after stimulus onset for scoring. We used the drifting-gratings global orientation selectivity measure provided with the Allen release as the validation measure for feature-attribution.

The speech dataset used the attempted-speech condition from the intracortical isolated-word recordings of Kunz et al.^36^ The main prediction task used participant t12, and the consistency analysis used participants t12, t15, t16 and t17. We defined an eight-class benchmark task comprising all released cue conditions: the seven attempted words analyzed by Kunz et al.—ban, choice, day, feel, kite, though and were—and the accompanying “Do Nothing” cue. For each participant, we stored threshold-crossing counts from the source-study motor array in 50 bins at 20 ms resolution: 500 ms immediately before the participant-specific attempted-speech decoding window followed by the 500 ms decoding window itself. The scoring window was therefore bins 25–49, corresponding to the 500 ms windows used by Kunz et al. We aligned this window to the participant- and array-specific start times as reported by Kunz et al.: 0 ms for t12 on array i6v, 1,000 ms for t15 on i6v, 2,100 ms for t16 on ventral6v, and 600 ms for t17 on s6v-v^36^.

The MC PacMan force-decoding dataset was released with the study introducing the MINT decoder^42^. It recorded motor-cortex activity while a monkey controlled a PacMan cursor by applying isometric force against an immovable handle to match scrolling force targets. We treated it as a continuous force-prediction task. We rebinned spikes to 20-ms counts and force to 20-ms means. We extracted fixed 2.24-s windows of 112 complete 20-ms bins beginning nominally 1.0 s before force-profile onset. The first 0.5 s was retained as input history but excluded from evaluation; scores used bins 25–111, beginning approximately 0.5 s before force-profile onset. When the nominal start fell before the available trial trace, we filled missing early samples by repeating the first rebinned value. The resulting benchmark array contained 362 trials, each with 112 time bins and 128 neural features, and the target was the corresponding one-dimensional force trajectory.

We generated the RatInABox dataset synthetically as a two-dimensional navigation task using RatInABox^38^ v1.15.3. The primary prediction analysis used one generated session, and latent-consistency analyses compared independently generated synthetic sessions. Across simulations, the virtual trajectory, place-cell field centers and Poisson spike sampling were reinitialized while the cell-class composition and simulation settings were held fixed. A virtual agent foraged for 1,500 s in a 1 m by 1 m open-field arena at 100 ms time steps, and we divided the continuous simulation into 300 non-overlapping 5 s trials with 50 bins per trial. The population comprised 100 Gaussian place cells with 0.2 m widths and 20 Hz maximum firing rate, 100 head-direction cells with 10 Hz maximum firing rate and 100 speed cells with 10 Hz maximum firing rate. We sampled spike counts from Poisson firing rates. For feature attribution, we used the known simulated cell-type labels. For latent consistency, we assigned each time bin to one of 100 fixed spatial bins in a 10 by 10 arena grid and used the mean latent representation for each shared spatial bin as the matched landmark across independently generated sessions.

### Data preprocessing and train/test splits

We converted each dataset to a common trial-by-time-by-feature array before model fitting. This step fixed the neural array, trial identifiers, task targets, condition identifiers when available, feature set and scored window. These choices were made before benchmarking and held fixed across methods.

Dataset-specific inclusion and feature rules were predefined. For the primary macaque-reaching session, we began with 320 rewarded trial records, excluded one record that lacked a go cue and retained 319. Hand position was interpolated 150 ms after each neural-bin center. Retention required the source-trial interval to span from at least 150 ms before to 850 ms after the go cue and at least two within-trial position samples to bracket all requested target times. Across the four-session latent-consistency cohort, 846 of 849 rewarded records were retained: two records, including the primary-session record above, lacked go cues, and one trial with a defined go cue failed the predefined trial-edge criterion because its source-trial interval ended approximately 10 ms before the required endpoint. For the same-subject historical-trial selection case study, we constructed a separate unsorted M1 electrode-channel representation from center-out reaching sessions of the same animal^34^, providing a fixed physical-channel feature space across old and current recordings. For Allen Neuropixels, we used units passing Allen default quality filters and applied no additional manual firing-rate, brain-region or performance-dependent filter. For the primary Speech benchmark, we used 64 threshold-crossing features from the selected motor array; the feature-stream sensitivity analysis compared these features with a combined 128-feature stream that added 64 spike-band-power-derived pseudo-count^36^ features. For participant T12, both streams used the same trial labels and train/test split, and each decoder used the same training and scoring procedure under both feature definitions. Across the 22 decoders with results for both streams, we defined Δaccuracy as combined-stream accuracy minus threshold-crossing accuracy, reported the unweighted mean of these decoder-level differences, and calculated the Spearman rank correlation between decoder accuracies under the two feature streams. For MC PacMan, we retained all 128 neural features in the released spike arrays. RatInABox used all simulated cells described above.

We assigned 80% of trials to training and 20% to testing with random seed 42. For the two classification tasks, Allen Neuropixels and attempted speech, trial splits were label-stratified. Except for MARBLE’s transductive representation-construction path described below, fitted normalization, dimensionality-reduction and scaling steps used in model preprocessing or benchmark scoring were estimated on training data only and then applied unchanged to held-out data. These training-only steps included PCA fitting and scaling, spike-history standardization for feedforward decoders, neuron z-scoring for recurrent decoders, target scaling for support-vector regression, and the method-specific preprocessing summarized in **Supplementary Note 3**. Primary shared ridge or logistic scoring models used the scale produced by each method unless the method integration specified a different training-only preprocessing step; trial-valuation scoring used the separate training-only scaling convention described in **Supplementary Note 5**.

### Scored time windows

We defined a scoring window for each dataset to match the task-relevant period. Models received the stored trial window as input, including pre-event history when present, unless a method-specific recipe required a different input construction described in **Supplementary Note 3**. We fit any shared models used to predict targets from fixed model outputs on the task window and evaluated all reported scores on the same window.

For macaque reaching, we scored the 700 ms after the go cue. For Allen Neuropixels, we scored the 2 s after stimulus onset. For speech, we scored the 500 ms attempted-speech window used by Kunz et al.^36^ For MC PacMan, we scored bins 25–111 of the 112-bin force sequence. For RatInABox, we scored the full 5 s navigation segment. For classification, class labels were unchanged when pre-event bins were excluded; predictions within the scoring window were aggregated to one trial label as described in **Supplementary Note 2**.

### Benchmark methods and prediction workflows

The evaluated methods and variants were BLEND-LFADS^21,24^, BLEND-NDT^24,52^, CEBRA^17^, DNN^1^, DPAD^25^, GPFA^20^, GRU^1^, LangevinFlow^23^, LDNS^22^, AutoLFADS^21,53,54^, LSTM^1^, MARBLE^18^, MINT^42^, NEDS^41^, NEDS-pt^41^, mVAE^55^, PCA^1,43^, RNN^1^, SMC-RNN^56^, SVM^1^, TNDM^26^, DFINE^27^ and XGBoost^40^. They span the cited method lineages and released implementations for GPFA^20^, LFADS/AutoLFADS^21,53,54^, CEBRA^17^, MARBLE^18^, BLEND^24^, Neural Data Transformers^52^, DPAD^25^, LDNS^22^, LangevinFlow^23^, NEDS^41^, mVAE^55^, SMC-RNN^56^, TNDM^26^, DFINE^27^ and standard machine-learning baselines^1,40,43^. NEDS used the released five-layer single-session configuration and was trained from scratch on each benchmark task. To probe whether large-scale cross-session pretraining transfers to these tasks, we additionally evaluated NEDS-pt, a 22-layer variant that followed Zhang et al.’s multi-session pretraining recipe^41^ by initializing from a NEDS checkpoint pretrained with multimodal masked modeling on 74 International Brain Laboratory repeated-site Neuropixels sessions^41,57^ and then fine-tuned end-to-end on each benchmark train split; NEDS is the method in our set with a published large-scale multi-session pretraining recipe, making it the natural case for this comparison. The pretraining corpus shares no recordings, subjects or tasks with any benchmark dataset, so NEDS-pt’s transfer is consistent with the reuse of learned neural-population structure and cannot be explained by overlap with evaluation data.

Before evaluating benchmark performance, we assigned method configurations from published or released method settings and prespecified dataset-specific adaptations described in **Supplementary Note 3**. Where an upstream task-family recipe or package default was available, we used the closest match by task family, data domain and input/output structure; we did not tune hyperparameters on benchmark results. Resource conventions, prediction representations and scoring models, consistency representations and model settings are summarized in **Supplementary Tables 3–6**.

Primary prediction scoring followed each method’s intended prediction target whenever the released workflow exposed one. Methods with native behavior heads or supervised prediction APIs were scored from their behavioral trajectory, class-probability or class-label outputs after applying the common scoring window.

For methods that exported factors, embeddings, rates or latent states, we used the method’s documented prediction step when available, such as CEBRA k-nearest-neighbor prediction^17^, MARBLE ordinary least-squares regression or k-nearest-neighbor regression^18^.

When a method exposed representations but no documented way to predict benchmark target values or class labels from them, we used a shared linear scoring model fitted on training representations and applied unchanged to test representations: ridge regression for continuous targets and logistic regression for class labels^43^. The estimator settings and feature-shaping convention are reported in **Supplementary Note 1**. This design evaluated task decoders through their intended prediction path and made representation-only methods comparable through the same ridge-or-logistic scoring rule. Method-specific departures are described in **Supplementary Notes 1 and 2**.

We separately identified methods whose fitting or representation-construction path used task-condition labels or test neural activity before final scoring. We recorded this distinction because these inputs change the information available before scoring and can shape the fitted trajectories or representations being evaluated. MINT used training condition labels for macaque reaching, Allen, Speech and MC PacMan; RatInABox followed MINT’s no-condition recipe. We interpreted MARBLE results as transductive representation results because MARBLE representation construction used train and test neural trials before downstream scoring. **Supplementary Note 4** reports dataset-level label-use and graph-construction details.

Some methods were originally designed for motor regression or neural reconstruction and did not cover every task type in the benchmark. We used a task adaptation only when the released implementation exposed a corresponding behavior head, loss, representation interface or documented way to predict benchmark target values or class labels from the model output; otherwise, the adaptation was reported as an added benchmark scoring model fitted on fixed model outputs, not as a native task decoder.

We evaluated primary prediction on test trials without artificial perturbation. We scored regression tasks by coefficient of determination, *R*², averaged uniformly across target dimensions when a dataset had more than one output dimension, and scored classification tasks by accuracy. We computed sequence-regression scores after slicing predictions and targets to the dataset-specific scoring window. We computed sequence-classification scores by aggregating framewise predictions within the scoring window to a trial-level prediction. Unless otherwise specified, we used the same score definitions for primary prediction, robustness, feature-attribution utilities and trial-valuation utilities.

For regression datasets, let *y_i_*_,*d*_ and *ŷ_i,d_* denote the target and predicted value for scored sample *i* and output dimension *d*, after applying the dataset-specific scoring window. The reported score was the uniform mean of per-dimension coefficients of determination,

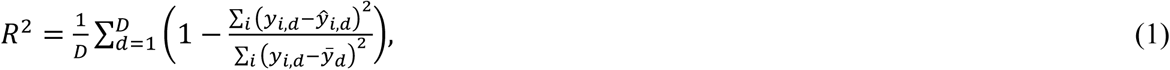

where *ȳ_d_* is the test-set mean target value for dimension *d*. For classification datasets, let *c_i_* be the trial label and *ĉ̂_i_* the majority-vote trial prediction from the scored time window. Accuracy was

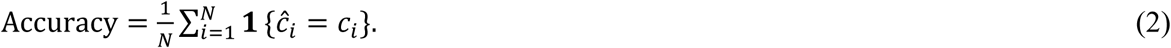

If framewise class predictions tied during majority voting, we used the tied class that appeared first in the scored window.

Alternative metrics were computed from the same zero-noise held-out predictions used for the primary analyses. Classification metrics were calculated after majority voting across the dataset-specific scoring window; macro or support-weighted averaging was used as indicated, with undefined precision, recall and F1 values set to zero. For regression, trials and scored time bins were pooled; RMSE was the square root of the mean squared error averaged uniformly across output dimensions, MAE was averaged uniformly across dimensions, and Pearson’s *r* and Spearman’s *r* were computed per output dimension and then averaged. Rankings were calculated within each dataset and metric using average ranks for ties.

### Robustness to added count noise

To assess whether performance was preserved under perturbation, we evaluated each trained model on noisy versions of the test neural data. We left the training data unchanged. For each dataset, we generated additive Poisson noise at five levels, 0, 0.2, 0.4, 0.6, and 0.8. At noise level *λ*, we replaced the test neural array with *X* + Poisson(*λX̄*), where *X* is the unperturbed test neural array and *X̄* is the scalar mean count over all entries of that array. We applied noise after the dataset loader and preprocessing steps had produced the unperturbed test array supplied to the robustness utility and before later inference-time transformations in the model wrapper. We reused identical noisy arrays across runs with the same unperturbed test array, noise level and seed. Conditional on *λX̄*, entries of the noise array were independent Poisson draws generated with a fixed seed. The zero-noise level used the original test array. For direct prediction models, we applied the trained model directly to the noisy test neural data. For methods evaluated through learned features, we transformed noisy test data with the trained representation procedure and scored the transformed features with the readout trained on unperturbed training features.

We summarized robustness as the trapezoidal area under the raw score-versus-noise curve. For noise levels *λ*_1_, …, *λ_K_* and corresponding test scores *S*(*λ_k_*), the robustness summary was

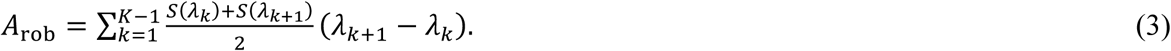

Because we used the raw task score, not a score normalized to the unperturbed value, this summary reflects both the unperturbed task score and the retention of performance as noise increases. It has task-score-by-noise-level units over the interval from 0 to 0.8, is not constrained to the interval from 0 to 1 and can be negative for *R*² tasks. The assay quantifies sensitivity to a controlled count-level perturbation of neural inputs.

### Runtime and memory assessment

We evaluated computational cost from the training and inference runs used in the benchmark. For each model-dataset evaluation, we measured wall-clock training time, wall-clock inference time, peak RAM and peak GPU memory when the evaluated workflow used a GPU. The macaque reaching task provided a common data shape for the compact main-figure comparison across all methods; per-dataset resource measurements are reported in **Supplementary Fig. 11**. Evaluations without GPU use were marked not applicable for GPU memory. Runtime and memory were measured under fixed GPU-enabled and CPU-only allocations; full tooling, hardware and measurement conventions are provided in **Supplementary Table 3** and the accompanying text. These measurements report practical runtime behavior under the benchmark hardware. For the budget examples in the Results, we paired each method’s mean prediction rank across the five primary tasks with its macaque-reaching training time and identified the best mean-rank method available at each time threshold.

### Cross-recording latent alignment

We evaluated cross-recording latent consistency for methods selected before evaluation because they produced method-defined, time-resolved representations suitable for landmark-based comparison across recordings. We use “representation” to include latent trajectories, learned embeddings, inferred factors, reconstructed-rate representations and low-rank states. The included methods were BLEND-LFADS, CEBRA, DPAD, GPFA, LDNS, AutoLFADS, MARBLE, mVAE, PCA, SMC-RNN, TNDM and DFINE (**Supplementary Table 5**). We chose this set because each method exposes a representation as part of its intended modeling workflow. We excluded direct supervised decoders whose only candidate representations were internal activations. We performed the analysis only for datasets with matched task structure across recordings, subjects or synthetic seeds.

We applied whitening after model fitting as a coordinate normalization step. This placed independently learned representations on a common scale so that the score emphasized alignment of matched task landmarks over recording-specific differences in latent variance or covariance.

For each recording, we first restricted latent trajectories to the dataset-specific scoring window and then constructed task landmarks in a dataset-specific manner. For macaque center-out reaching, the mean hand trajectory for each reach condition defined a reach direction; we projected trial-time samples onto that direction, normalized them by reach length and divided them into ten uniform progress bins. Each macaque landmark was the mean latent vector for one reach-condition/progress-bin combination. For RatInABox, each landmark was the mean representation over trial-time samples assigned to one of 100 fixed spatial bins. For classification datasets, landmarks were class centroids. We excluded landmarks that were not shared across the recordings being compared.

Before alignment, we centered and whitened the landmark matrix within each recording. For a landmark matrix *Z*, with landmarks as rows, we centered each column to obtain *Z_c_*, computed the sample covariance across landmark rows using the *n* − 1 normalization, and whitened the centered matrix as

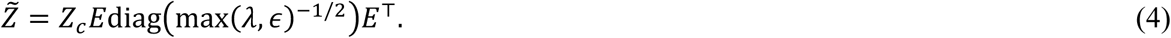

where *E* and *λ* are the covariance eigenvectors and eigenvalues, respectively, and the eigenvalue floor *ε* = 10^−8^ stabilizes the inverse square-root. For recordings *a* and *b*, let *Z̃_a_* and *Z̃_b_* be matrices whose rows are the matched whitened landmarks. The directional alignment from *a* to *b* fit an intercept-free linear map,

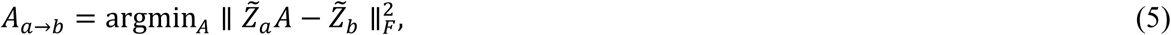

and scored the aligned source landmarks by uniform-average *R*^2^ against the target landmarks,

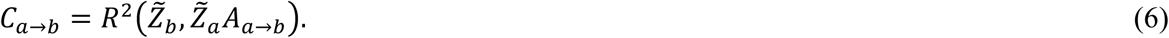

We reported the consistency score as the mean of *C_a_*_→*b*_ and *C_b_*_→*a*_ for each recording pair, averaged across pairs. We computed Orthogonal Procrustes alignment^58^ as a secondary diagnostic. The score quantifies empirical alignment of matched task landmarks across recordings after coordinate normalization.

### Feature-attribution estimation with Kernel SHAP

We estimated feature attribution with Kernel SHAP to test whether a trained model assigned task utility to inputs that agreed with validation targets defined before feature scoring. Kernel SHAP re-evaluated each trained decoder on masked neural inputs and scored each masked evaluation with the same task metric, giving a common feature-attribution procedure across model families. These analyses provided global feature-attribution summaries of benchmark utility over an evaluation split. We used the term feature because the input dimensions differed across datasets: some were spike-sorted units, some were channel-level or spike-band-power features, and some were synthetic control features.

SHAP values are based on the Shapley value from cooperative game theory^30,39^. Let *F* be the set of input features, *j* one feature and *u*(*S*) the scalar task score on the evaluation split after retaining feature subset *S* and replacing excluded features with their training-set means. The target feature value was

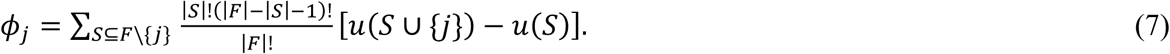

Kernel SHAP estimated this target quantity from sampled feature coalitions without exhaustive enumeration of all feature subsets. The model state being explained was the unmasked evaluation, with all input features retained, and the reference state was the fully masked evaluation, with all input features replaced by training-set means. Intermediate coalitions specified which features were retained for each masked evaluation. In this benchmark, the utility function was a single task score for a fixed trained model under a specified feature mask. For classification tasks, that scalar score was aggregate accuracy after scoring-window aggregation, so a feature value estimates contribution to global accuracy. For regression tasks, the scalar score was *R*². A positive value therefore means that including the feature increased the benchmark utility under the masking convention, whereas a negative value means that including the feature decreased the utility. All masked-evaluation scores were computed on the dataset-specific scoring window. Implementation details for mask imputation, coalition sampling and coefficient retention are reported in **Supplementary Note 6**.

We defined validation targets separately for each dataset. For macaque reaching, speech and MC PacMan, we appended synthetic noise features with no target information and measured whether original recorded features received larger signed feature-attribution values than synthetic features. The number of synthetic controls was approximately 25% of the number of original features, with values generated from a train-derived global rate (**Supplementary Note 6**). We summarized performance by ROC-AUC with original features as positives and signed feature-attribution values as ranking scores. For Allen Neuropixels, we compared feature values with each unit’s drifting-gratings global orientation selectivity using Spearman correlation; constant attribution vectors, which provide no feature ordering, were assigned zero agreement in benchmark summaries. For RatInABox, AUC used place cells as positives, head-direction and speed cells as negatives and signed feature-attribution values as ranking scores; mean values were also summarized by cell type. We retained the sign of each value for ranking and summaries because the analysis tested whether a feature increased or decreased the global utility under the mask. Feature grounding quantified agreement between signed feature-attribution values and each dataset’s prespecified validation target, using ROC-AUC or Spearman correlation as described above.

The validation targets differed in the degree of ground-truth control they provided: appended synthetic controls and RatInABox cell classes provided known labels, whereas Allen orientation selectivity provided a biologically motivated proxy.

### Trial-level data valuation

We used trial-level data valuation to estimate which training examples helped or harmed a concrete benchmark-task utility. We used Data Shapley^32^, which applies the Shapley value^39^ to training examples. Let *T* be the set of candidate training trials, *i* one trial and *v*(*S*) the utility assigned to a sampled subset *S*. In our implementation, for subsets that supplied enough data to train the scoring model from fixed learned features to benchmark targets, *v*(*S*) was the valuation score obtained after training that model on trial subset S and scoring it on the valuation split.

Empty subsets, and classification subsets containing only one class, were assigned baseline utilities because no classifier could be trained from those subsets; **Supplementary Note 7** defines those baselines.

The trial value was

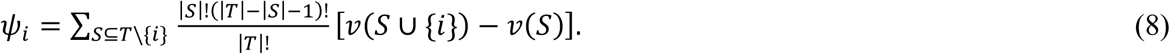

The estimator repeatedly samples subsets of training trials; in this benchmark, we held the full trained model fixed and retrained only a lightweight scoring model on each sampled subset when trial valuation required subset-specific scoring. Depending on the method, this scoring model matched the model used for primary prediction from fixed features, or it was a shared linear model trained from fixed features to target values or class labels. A trial value is therefore specific to the fixed model features, scoring model, valuation split, candidate pool and performance metric.

For the main corrupted-trial analysis and within-session corrupted-trial removal case study, we computed fixed model features for training trials and used the test split as the valuation split. We trained the subset-specific scoring model only on sampled subsets of training trials. The same-subject historical-trial case used the split convention described below.

For classification tasks, we computed utilities for nonempty subsets as accuracy after slicing to the scored window and aggregating framewise class outputs or predictions from the scoring model trained on sampled training trials with the same trial-level classification rule used for prediction scoring. For regression tasks, we computed utilities for nonempty subsets as *R*². We estimated trial values with truncated Monte Carlo sampling over random permutations of the candidate training trials, using up to 3,000 permutations for corrupted-trial valuation summaries, including the within-session corrupted-trial removal case study, and up to 2,000 permutations for same-subject historical selection. Baseline utilities and sampling details are reported in **Supplementary Note 7**.

Applying Data Shapley required recomputing utility for many sampled subsets of training trials. We therefore limited BEND-BCI trial valuation to method-dataset combinations whose subset utilities could be recalculated by training only the valuation scorer, without rerunning the full primary decoder for every subset. This rule included 17 methods and excluded DPAD, DFINE, MINT, mVAE, SVM and XGBoost from the corrupted-trial valuation grid (**Supplementary Note 5**).

### Corrupted-trial detection with trial values

To generate corrupted training trials, we applied a subspace-rotation perturbation to selected training examples. For each dataset, we centered available training neural samples by their mean, projected them into the full SVD basis of the centered training activity, rotated paired basis coordinates and transformed the rotated samples back to neural-feature space. The basis provided a stable data-adaptive coordinate system for perturbation.

The corrupted-trial assay used a 75-degree rotation and requested perturbation of 0.33 of the training trials, approximately one third. We selected trials randomly without replacement from available training trials. We converted the requested fraction to a count by rounding 0.33*n*_train_, with at least one selected trial when the fraction was nonzero and with the count capped at the number of available trials. We applied perturbations to the full training window before scoring-window slicing. We quantified corrupted-trial detection by ROC-AUC using the negative trial value as the detection score, because perturbed trials were expected to have lower utility than unperturbed trials.

### Trial-value-guided retraining case studies

We evaluated two macaque reaching case studies to test whether trial valuation could guide data selection. In the within-session corrupted-trial removal case study, we trained each model on a mixture of unperturbed trials and trials corrupted by the same subspace-rotation perturbation used in the corrupted-trial assay. We computed Data Shapley values over candidate training trials using performance on the benchmark test split as the utility function. We then removed training trials with negative signed value, retrained each model only on the retained training trials and compared test reaching performance with retraining on the mixed corrupted set. Thus, benchmark-test examples were not used for model fitting, but the final comparison was post-selection with respect to the valuation split. We interpret this analysis as a controlled test of data selection for the benchmark objective under synthetic training-trial corruption.

In the same-subject historical-trial selection case study, we used an unsorted channel-level M1 representation, not spike-sorted units because unit identities can vary across sessions. We pooled all spike trains assigned to the same physical M1 electrode before binning, producing a fixed channel-level feature space across old and current sessions for every decoder. This choice follows prior evidence that threshold-crossing and multiunit channel features can preserve low-dimensional motor population structure for population-level analyses and BCI decoding^59,60^. The target session was 20150716. We split its 191 trials into 28 current-session training trials, 28 validation trials from the current session and 135 held-out test trials from the current session, with the current-session training set intentionally capped to create a low-calibration setting.

First, we trained models using the current-session training set plus 992 old candidate trials from seven earlier same-subject center-out sessions to produce fixed learned features in a common channel-level feature space.

Second, we assigned trial values only to historical candidate trials. We held the learned features from the first training run fixed; Shapley sampling varied only which old trials were used to retrain the scoring model. For historical-trial scoring, each sampled coalition retrained that model on old candidate trials and scored transfer to the current-session training trials. An empty old-trial coalition used a null *R*² value of 0. We did not assign values to current-training trials.

Third, we retrained each model for the final comparison using the current-session training set plus historical trials after excluding those with negative signed Data Shapley values, corresponding to older trials whose average marginal contribution to transfer utility on the current-session training trials was detrimental. We evaluated performance on held-out test trials from the current session. The separate 28-trial validation subset from the current session was reserved by the split definition but was not used as the utility split for this historical-selection case study.

Baselines included current-only training and all-session pooling. These comparisons defined the scope of the case study: we tested trial valuation as a selector of useful historical examples when current and old recordings were represented in a common physical-channel space.

### Inclusion rules and statistical analysis

We generated benchmark matrices from method-dataset evaluations after applying eligibility requirements defined before each analysis. Missing entries and their causes are reported in **Supplementary Table 2**. Display matrices mark these entries explicitly. Aggregate mean-rank summaries assign missing eligible entries the worst within-dataset rank.

For summary heat maps, metrics were compared within dataset to avoid conflating model performance with dataset difficulty or metric scale. When overall rankings were reported, methods were ranked within each dataset and then aggregated across datasets or analysis categories. For paired retraining comparisons, percentage changes in mean *R*² were calculated as 100 × [mean(*y*) − mean(*x*)]/mean(*x*), where *x* and *y* denote the axis conditions. Associations between benchmark measurements were assessed using Spearman rank correlations. For each pair of measurements, we filtered to model–dataset entries with both measurements, converted each measurement to a within-dataset percentile rank over that shared set and pooled entries across datasets. Unless otherwise specified, Spearman-correlation *P* values were two-sided values from scipy.stats.spearmanr^61^. For the association between corrupted-trial detection ROC-AUC and recovery after trial removal, we used a one-sided permutation test of Spearman’s *r* for a positive association, based on 1,000,000 permutations of the recovery values with seed 42; the Monte Carlo *P* value used a +1 correction. ROC-AUC values followed the positive-label conventions described above for feature-attribution validation and corrupted-trial detection, and can be read as the probability that a randomly selected positive example is ranked above a randomly selected negative example, with ties assigned half credit^62^. We used random seed 42 for dataset splitting, perturbation generation and trial-valuation sampling.

## Data availability

This study analyzed previously published neural datasets and generated synthetic RatInABox data. Macaque reaching data are available from DANDI Archive 000688, version 0.250122.1735 (https://doi.org/10.48324/dandi.000688/0.250122.1735). Allen Visual Coding Neuropixels data are available through the AllenSDK (https://allensdk.readthedocs.io/en/stable/visual_coding_neuropixels.html); the session identifiers used here are listed in **Supplementary Table 1**. Attempted-speech data are available from Dryad (https://doi.org/10.5061/dryad.gf1vhhn1j), and MC PacMan data are available from Zenodo (https://doi.org/10.5281/zenodo.14769889). RatInABox data were generated using the procedure described in **Methods**; generation seeds for the primary and consistency sessions are provided in **Supplementary Table 1**. Upon publication, machine-readable benchmark outputs underlying the figures and reported results will be released in the associated Code Ocean capsule.

## Code availability

The custom code and computational environments used to prepare the datasets, run the benchmark analyses and generate the reported results are available to editors and reviewers upon request. A frozen version will be released publicly through Code Ocean with a permanent DOI upon publication. The development repository (https://github.com/TangLab-UBC/behavior_benchmarking) and interactive dashboard (https://tang-lab-benchdash.hf.space/) will also be made public upon publication.

## Acknowledgments

Computations were performed on the Trillium supercomputer at the SciNet HPC Consortium, with support from the Digital Research Alliance of Canada. This work was supported by start-up funding from the University of British Columbia, the Canada Research Chairs Program [CRC-2024-00164], and the Natural Sciences and Engineering Research Council of Canada Discovery Program [RGPIN-2025-04460], to X.T. The research infrastructure used in this work was supported by the Canada Foundation for Innovation John R. Evans Leaders Fund and the British Columbia Knowledge Development Fund [45547].

## Author contributions

J.S. and X.T. conceived the study and designed the benchmarking strategy. J.S. designed and implemented the benchmarking framework, developed the software, performed the experiments, analyzed the results, prepared the figures, and wrote the initial draft. K.P.S. reviewed code, assisted with dataset identification and preliminary dataset preprocessing, contributed to drafting the manuscript, and prepared the Figure 1 schematic. All authors discussed and interpreted the results. X.T. supervised the project and acquired funding. All authors reviewed and edited the manuscript.

## Competing interests

The authors declare no competing interests.

## Additional information

Correspondence and requests for materials should be addressed to Xin Tang.

## Notes

### Competing Interest Statement

The authors have declared no competing interest.

