## Supplementary Information for "Benchmarking Neural Decoders for Brain-Computer Interfaces and Neural Population Analysis"

### Supplementary Notes, Tables and Figures


**Supplementary Note 1**: Benchmark method integration and prediction scoring
**Supplementary Note 2**: Classification scoring and task adaptations
**Supplementary Note 3**: Preprocessing, model recipes and run-defining settings
**Supplementary Note 4**: MINT and MARBLE condition annotations
**Supplementary Note 5**: Model outputs and scoring models for trial valuation
**Supplementary Note 6**: Feature-attribution estimator conventions
**Supplementary Note 7**: Data Shapley sampling, perturbation and split rules
Supplementary Table 1: Benchmark datasets
Supplementary Table 2: Analysis availability and missing entries
Supplementary Table 3: Resource and runtime conventions
Supplementary Table 4: Prediction representations, outputs and scoring models
Supplementary Table 5: Consistency representations
Supplementary Table 6: Model configurations, preprocessing and training settings
Supplementary Fig. 1 | Classification rankings across scoring metrics.
Supplementary Fig. 2 | Regression rankings across scoring metrics.
Supplementary Fig. 3 | Agreement among prediction metrics.
Supplementary Fig. 4 | Attempted-speech feature-stream sensitivity.
Supplementary Fig. 5 | Robustness curves across benchmark datasets.
Supplementary Fig. 6 | Macaque reaching robustness examples.
Supplementary Fig. 7 | MC PacMan robustness examples.
Supplementary Fig. 8 | Allen Neuropixels robustness examples.
Supplementary Fig. 9 | Attempted-speech robustness examples.
Supplementary Fig. 10 | RatInABox robustness examples.
Supplementary Fig. 11 | Computational cost measurements across datasets.
Supplementary Fig. 12 | Prediction–resource frontiers.
Supplementary Fig. 13 | Macaque reaching latent-consistency visualizations.
Supplementary Fig. 14 | Allen Neuropixels latent-consistency visualizations.
Supplementary Fig. 15 | Attempted-speech latent-consistency visualizations.
Supplementary Fig. 16 | RatInABox latent-consistency visualizations.
Supplementary Fig. 17 | Cross-analysis metric correlations.
Supplementary Fig. 18 | Feature-attribution validation with synthetic controls and simulated cell classes.
Supplementary Fig. 19 | Allen orientation-selectivity validation.
Supplementary Fig. 20 | Highest- and lowest-ranked feature examples.
Supplementary Fig. 21 | RatInABox cell-class atlas.
Supplementary Fig. 22 | Macaque reaching trial-valuation summaries.

Supplementary Fig. 23 | MC PacMan trial-valuation summaries.
Supplementary Fig. 24 | Allen Neuropixels trial-valuation summaries.
Supplementary Fig. 25 | Attempted-speech trial-valuation summaries.
Supplementary Fig. 26 | RatInABox trial-valuation summaries.
Supplementary Fig. 27 | Same-subject historical-trial selection controls.

### Supplementary Note 1: Benchmark method integration and prediction scoring

We preserved released or documented model training and prediction procedures when they could operate on trial-by-time neural activity and benchmark targets. Here, benchmark targets are the variables scored by BEND-BCI: continuous hand position, force or two-dimensional position for regression tasks, and trial labels for classification tasks. We set the scoring rule before inspecting benchmark performance. Each reported prediction score used the same dataset split, scoring window and task metric within a task; the scoring window is the set of time bins used for evaluation. The scoring rules fell into three explicit cases: predictions generated directly by the trained model; method-specific mappings from learned features or reconstructed firing rates to benchmark targets; or a standard ridge/logistic readout, defined here as a lightweight model trained after the primary model to map fixed model outputs to benchmark targets.

Methods trained to predict benchmark targets generated target trajectories or class scores during the primary model run. Continuous tasks used decoded target trajectories. Classification tasks used class labels, class logits or one-hot class trajectories, with predictions from individual time bins aggregated to one trial label over the scoring window. Method-specific representations or outputs used for prediction and their scoring models are listed in **Supplementary Table 4**.

We distinguished prediction procedures specified by a method recipe from prespecified benchmark adaptations. A method recipe was defined as a mapping described in the source paper, a released example, or the documented package defaults. CEBRA[1] embeddings were trained using benchmark targets or class labels as supervised contrastive labels and evaluated with k-nearest-neighbor prediction, following the released CEBRA workflow. For MARBLE[2], macaque center-out reaching used ordinary least-squares regression, and the same mapping was applied to MC PacMan as a motor-task adaptation. RatInABox followed the released k-nearest-neighbor position-prediction recipe. Because MARBLE provides no released recipe for visual or speech classification, Allen and Speech used a standard logistic readout on fixed MARBLE embeddings, as described in **Supplementary Note 3**. LDNS[3] decoded continuous targets from autoencoder-reconstructed firing rates using ridge regression with a regularization parameter of $10^{-6}$. Because LDNS provides no upstream classification recipe, its classification analyses used the standard logistic readout on the same fixed reconstructed-rate features.

For methods that produced learned features or rates but no model component for predicting a BEND-BCI target, we trained the standard ridge/logistic readout from fixed outputs to benchmark targets. Fixed outputs are representations or firing-rate estimates computed once by the trained primary model and held constant while the readout is trained; the specific outputs are listed in **Supplementary Table 4**. For BLEND[4], the neural-only student received only neural input at test. Continuous targets used ridge regression and classification targets used logistic regression. For LangevinFlow[5] continuous prediction, this matched the Neural Latents Benchmark behavioral-decoding convention: fit ridge regression from training rates to target trajectories, then apply the fitted mapping to test rates[6].

BLEND-LFADS and BLEND-NDT[4,7] followed this scoring rule for fixed model outputs. During BLEND training, we supplied continuous benchmark targets, or repeated one-hot class labels for classification, as privileged training variables, omitted held-out-neuron and future-prediction streams, and used the neural-only student at test time.

For standard ridge or logistic scoring in the primary prediction analysis, we restricted model outputs to the dataset scoring window and flattened them over trial and time before training the readout. We did not apply additional feature z-scoring at this step; the readout used the scale produced by the model. Continuous targets used ridge regression with regularization 1.0. Class labels used logistic regression with default L2 regularization and a maximum of 1,000 iterations. We repeated trial-level class labels across the scoring window for training and aggregated time-bin predictions by majority vote for scoring.

**Supplementary Note 2: Classification scoring and task adaptations**

The supervised baseline classifiers were trained directly to predict benchmark targets without adding classifiers on learned features after training. DNN[8] used a 400-unit feedforward classifier trained with categorical cross-entropy on one-hot trial labels. RNN, GRU and LSTM[8] used 400-unit sequence classifiers with trial labels repeated across time and time-bin predictions aggregated by majority vote. SVM/SVR[8] and XGBoost[9] used spike-history features and direct supervised classifier or regressor outputs, with time-bin classification predictions aggregated to trial labels.

Several sequence models were trained directly against classification targets without adding a separate classifier after model training. NEDS and NEDS-pt[10] trained their transformer prediction heads against repeated class labels with categorical cross-entropy. DPAD[11] received integer class labels as categorical target values, learned its behavior-readout mapping (Cz) from latent states to class-score outputs and converted those scores to labels by argmax before majority-vote scoring over the window. TNDM[12] includes a model step that predicts targets from task-relevant latent factors. For classification, BEND-BCI used this step with repeated one-hot label sequences and trained the class scores with categorical cross-entropy while preserving TNDM's relevant and irrelevant factor structure.

DFINE[13] predicted target trajectories from smoothed manifold latent factors. For classification targets, we used the same manifold-factor-to-target prediction step with one output dimension per class and trained the class scores with cross-entropy on class labels repeated across time.

MINT[14] builds condition-indexed neural and target trajectory libraries, so classification used the class label as the condition identity and represented each class as a repeated one-hot target trajectory. At each time bin, the decoded trajectory was converted to a class label by argmax, and time-bin labels were aggregated by majority vote over the scoring window. The trajectory-library smoothing and causal observation-window settings are reported in **Supplementary Note 3**.

For mVAE[15] classification, we retained the released neural-behavioral masking architecture, represented each trial label as a repeated one-hot behavioral stream and trained the behavioral decoder with categorical cross-entropy in place of the continuous-target Gaussian likelihood. At test time, the entire behavioral stream was masked, so class scores were generated from neural input alone.

CEBRA[1] used k-nearest-neighbor prediction from CEBRA embeddings. For the other classification entries without a model trained to produce class scores, we trained logistic regression from the listed fixed outputs to class labels: MARBLE[2] embeddings, LDNS[3] reconstructed firing rates, LangevinFlow[5] predicted firing rates, GPFA[16] latent trajectories, principal-component coordinates[8,17], AutoLFADS[18–20] factors, SMC-RNN[21] posterior-mean latent trajectories, and representations from the BLEND[4,7] neural-only student.

### Supplementary Note 3: Preprocessing, model recipes and run-defining settings

Preprocessing followed a released task recipe when available; otherwise, we prespecified the closest task-family adaptation before examining benchmark results. CEBRA[1] applied the 40 ms Gaussian smoothing used in its macaque motor recipe only for macaque reaching and MC PacMan; Allen, Speech and RatInABox used unsmoothed counts.

For MARBLE[2], macaque reaching followed the released macaque preprocessing pipeline: Gaussian smoothing with a 100 ms standard deviation, Savitzky–Golay filtering[22] with a window length of 9 and polynomial order of 2, and dimensionality reduction to five principal components. MC PacMan used the same preprocessing as a motor-task adaptation, with ordinary least-squares regression for target prediction. RatInABox followed the released navigation recipe, using Gaussian smoothing with a 10 ms standard deviation, no Savitzky–Golay filtering, dimensionality reduction to 20 principal components, and k-nearest-neighbor regression for position prediction. Because MARBLE does not provide released recipes for visual or speech classification, Allen and Speech used the macaque preprocessing pipeline as a prespecified benchmark adaptation, with a three-dimensional MARBLE representation and logistic regression for class prediction.

DPAD received each benchmark dataset's canonical binned neural inputs. For continuous-target analyses, we additionally applied Gaussian smoothing with a 50-ms s.d., matching the smoothing width used in the published DPAD movement analyses[11]. Because DPAD has no released visual or speech classification preprocessing recipe, these analyses retained the benchmark's binned counts without additional Gaussian smoothing.

MINT used released task-family settings where available. Macaque reaching transferred the released MC Maze motor configuration with a 300-ms causal window, 30-ms Gaussian trajectory smoothing, Type II trial averaging and two cross-trajectory candidates. We retained the benchmark's canonical absolute-position target and full condition dimensionality because the benchmark has eight reach conditions, fewer than the 21 condition dimensions in the released configuration. MC PacMan used its task-specific 35-ms smoothing setting with the same 300-ms causal window and Type II averaging. Because MINT does not provide a released visual or speech classification recipe, Allen and Speech used the 30-ms trajectory-library smoothing width from the released MC Maze configuration and a 12-observation causal window matching the observation count of the released Area2_Bump configuration. We held this observation count fixed across both classification tasks on their native sampling grids; the window comprised the current and 11 preceding bins. RatInABox followed the published MINT MC RTT no-condition procedure. In the original MINT study, Perkins et al. ran AutoLFADS[18–20] twice on the MC RTT training data and averaged the two rate estimates to reduce the influence of run-specific rate variability. We transferred that procedure to RatInABox using only training data and the released LFADS-Torch MC RTT architecture, with 40 factors and 100 generator units; held-out MINT inference used raw spike counts. At RatInABox's native 100-ms grid, we represented the MC RTT 480-ms causal window with the closest available five-bin window, 500 ms. Following the MC RTT interpolation procedure, we used six within-trajectory candidates separated by at least 1 s and applied no additional Gaussian smoothing[14].

LDNS used raw binned counts and LDNS's autoencoder preprocessing[3]: continuous-target analyses and non-speech classification used the released monkey autoencoder recipe, whereas attempted-speech classification used the released human/phoneme autoencoder recipe adapted to the benchmark's fixed-length isolated-word trials. NEDS, NEDS-pt[10], mVAE[15], LangevinFlow[5], AutoLFADS[18–20] and SMC-RNN[21] received raw binned counts and otherwise followed their model-specific preprocessing. DFINE[13] z-scored neural observations with training-split statistics before model input. GPFA used square-root transformed counts[16] and removed neurons with zero activity across the GPFA training split; PCA[8,17] used raw counts, scaling fit on training data and PCA fit on training trial-time samples.

**Supplementary Note 4: MINT and MARBLE condition annotations**

MINT used training condition labels to index neural and target trajectory libraries for macaque reaching, Allen, Speech, and MC PacMan; RatInABox used the no-condition construction described in **Supplementary Note 3**. Held-out condition labels were not used for prediction. MINT predicted target trajectories from held-out spike sequences, which BEND-BCI compared with held-out targets to compute $R^2$ or accuracy.

MARBLE representation construction was transductive for all datasets because it fit PCA and the MARBLE transformation on train and test neural trials before final scoring. For macaque reaching and MC PacMan, MARBLE also used condition-stratified graph construction with reach-direction conditions for macaque reaching and eight force-profile conditions for MC PacMan. Allen, Speech and RatInABox MARBLE runs used a single graph: class labels and RatInABox spatial-bin labels were not used to construct MARBLE graphs. These annotations describe representation construction, while final scoring models were fit without test labels.

For MARBLE robustness, the PCA transform, trained MARBLE network, graph construction rule and target-prediction model from the unperturbed benchmark run were reused at every noise level. The zero-noise score used the clean test embeddings from the original transductive fit. For nonzero noise levels, noisy test arrays were passed through the same MARBLE preprocessing, projected with the reused PCA transform and transformed by the reused MARBLE network. The condition-stratified graph construction was reused only for macaque reaching and MC PacMan, matching the unperturbed MARBLE run. Allen, Speech and RatInABox used a single graph for noisy test embeddings, so class labels and RatInABox spatial-bin labels were not introduced during robustness scoring.

**Supplementary Note 5: Model outputs and scoring models for trial valuation**

Trial-level valuation did not retrain the full primary model for every sampled subset of training trials. It held each method's learned features, factors, rates or latents fixed, and retrained only a lightweight scoring model on sampled training-trial subsets. These valuation scoring models mapped fixed model outputs to benchmark targets.

For CEBRA, MARBLE, LDNS and LangevinFlow, trial valuation used the same fixed model outputs and prediction models as the primary score (**Supplementary Table 4**), applied to each sampled training subset. TNDM valuation used least-squares or logistic models on task-relevant latent trajectories while preserving the causal time restriction used for TNDM target prediction; the full TNDM model remained fixed across sampled subsets.

For the remaining included methods, valuation trained ridge or logistic models on fixed outputs available after the primary model run. This group used BLEND-LFADS factors, representations from the BLEND-NDT neural-only student, learned features from DNN, RNN, GRU, LSTM, NEDS and NEDS-pt, GPFA latent trajectories, AutoLFADS factors, principal-component coordinates and SMC-RNN normalized posterior-mean latent trajectories. For BLEND-LFADS, BLEND-NDT, GPFA, AutoLFADS, PCA and SMC-RNN, this ridge/logistic scoring rule matched the primary prediction rule. For DNN, RNN, GRU, LSTM, NEDS and NEDS-pt, primary prediction used the full model trained to predict benchmark targets, whereas trial valuation used fixed learned features only to estimate trial utility without retraining the full model.

Before training trial-valuation scoring models, we sliced model outputs to the scoring window. Trial-valuation scoring models z-scored each feature using training-trial samples only before training, except for CEBRA k-nearest-neighbor prediction, MARBLE k-nearest-neighbor prediction for RatInABox, the TNDM causal-linear valuation model and the LDNS reconstructed-rate model; these models used the feature scale from the primary model output.

The corrupted-trial valuation analysis therefore included BLEND-LFADS, BLEND-NDT, CEBRA, DNN, GPFA, GRU, LangevinFlow, LDNS, AutoLFADS, LSTM, MARBLE, NEDS, NEDS-pt, PCA, RNN, SMC-RNN and TNDM. We excluded DPAD, DFINE, MINT and mVAE because their prediction scores came from full models trained to predict benchmark targets, and those scores could not be reproduced by retraining only a lightweight scoring model on fixed outputs. We excluded SVM and XGBoost because valuing arbitrary training-trial subsets would have required retraining the flattened-feature classifier or regressor for each sampled subset, unlike the included methods for which valuation retrained only lightweight scoring models.

**Supplementary Note 6: Feature-attribution estimator conventions**

Kernel SHAP[23] used mean-imputation masks. Masked evaluations replaced excluded input features with their training-set mean activity, and scores were computed only on the dataset-specific evaluation window. Feature values remained signed throughout the feature-ranking, correlation and AUC summaries. The feature-attribution runs passed nsamples="auto" to shap_values, using SHAP's automatic coalition-sampling budget. The weighted local-linear fit used l1_reg="num_features(N)", where N was the number of neural input features being valued.

For synthetic-control feature attribution, the controls used one global mean count estimated from the original training split across all original features, trials and time bins. The same train-derived rate generated synthetic train and test controls. This construction fixed a reproducible target-free control distribution; the controls were not treated as independent biological features.

For the consensus feature ranking, we averaged each feature's within-model rank based on signed SHAP values across models meeting the dataset-specific validation criterion. We included models with ROC-AUC ≥ 0.55 for the synthetic-control and RatInABox comparisons and Spearman's $r \geq 0$ for Allen; constant attribution vectors were excluded.

Feature-attribution results were unavailable when repeated masked-input evaluation was incompatible with a method or exceeded computational limits. MARBLE results were unavailable for all five datasets: the Allen run exceeded memory limits before a SHAP vector could be generated, whereas masked-feature evaluations for macaque reaching, Speech, MC PacMan, and RatInABox invalidated MARBLE graph construction. The MINT evaluation for Allen did not complete the MATLAB-backed repeated masked-input procedure within the allocated execution time. For SVM, the Allen run exceeded the 24 h limit during initial exact kernel-SVC fitting, whereas the MC PacMan and RatInABox runs exceeded the 24 h limit during repeated masked-input SVC or SVR evaluations. Accordingly, the feature-attribution matrix includes 22 methods and four additional unavailable cells, with MARBLE omitted (**Fig. 4**).

**Supplementary Note 7: Data Shapley sampling, perturbation and split rules**

Trial-level valuation used truncated Monte Carlo Data Shapley sampling over random permutations of candidate training trials, following the released TMC-Shapley implementation conventions of Ghorbani and Zou[24]. A coalition is the subset of candidate training trials accumulated at a given step of a permutation, and coalition utility is the score obtained after training the valuation scoring model on that subset. Valuation summaries based on the controlled rotation, including the within-session corrupted-trial removal case study, used a 3,000-permutation cap per summary. Same-subject historical selection used 2,000 permutations per old-only transfer valuation summary.

Within a sampled permutation, valuation stopped after more than five consecutive trial additions whose coalition utility was within $0.01 \times \max(|v(T)|, 10^{-12})$ of the full-candidate utility; remaining trials in that permutation were assigned zero marginal contribution.

Empty subsets used baseline utilities because no scoring model could be trained with zero trials. For classification, the baseline was the majority-class fraction from candidate training labels, and one-class coalitions used the same baseline because a classifier cannot be trained from a single class. For regression, the baseline was the mean score from target-permutation scoring models. In the same-subject historical-selection analysis, the old-only transfer utility assigned null $R^2 = 0$ to an empty old-trial coalition because no transferred scoring model could be trained.

The controlled rotation applied the same 75-degree subspace rotation used in the trial-valuation analyses. It centered training neural samples, projected them into the full SVD basis of the centered training activity, rotated consecutive coordinate pairs and transformed the rotated coefficients back into neural-feature space. Nonnegative count arrays were clipped at zero and rounded back to integer counts after rotation. Perturbed trials were sampled from training trials without replacement using the predefined perturbation seed. The requested fraction was 0.33; the number of selected trials was the nearest integer to 0.33 times the number of available training trials, with a minimum of one selected trial when the fraction was nonzero and a maximum equal to the number of available training trials.

The controlled-rotation analysis trained valuation scoring models on sampled subsets of training trials and scored them on the benchmark test split. The within-session corrupted-trial removal case study used the same benchmark test split for valuation utility and final retraining comparison, providing a controlled test of benchmark-objective data selection under synthetic training-trial corruption. The same-subject historical-selection analysis used target session 20150716: historical-trial coalitions trained the scoring model on sampled old trials only and scored transfer to the current-session training trials; selected historical trials were then added to the current-session training set and final comparisons used held-out test trials from the current session.

We used negative trial value as the corrupted-trial detection score.

For the corrupted trial detection, we used one-sided Mann–Whitney $U$ tests to test whether unperturbed trials had higher signed trial values than rotated trials.

**Supplementary Table 1: Benchmark datasets**

| Dataset | Source/session identifiers | Main task | Preprocessed trials x time bins x features | Target and score |
|---|---|---|---|---|
| Macaque center-out reaching | DANDI 000688[25,26]; main C-20151104; consistency sessions from monkeys C, J, M and T; historical target C-20150716 plus seven earlier C sessions. | Center-out reaching | 319 x 50 x 59 | 2D hand position, $R^2$ |
| Allen Neuropixels | Main session 721123822; consistency sessions 715093703, 721123822 and 732592105[27]. | Drifting-grating visual coding | 598 x 300 x 444 | Eight-way orientation class, accuracy |
| Attempted speech | Main participant t12; consistency participants t12, t15, t16 and t17[28]. | Isolated-word attempted speech | 168 x 50 x 64 | Eight-class attempted-speech label, accuracy |
| MC PacMan | Released MC PacMan force dataset from the MINT study[14,29]. | Force decoding | 362 x 112 x 128 | One-dimensional force, $R^2$ |
| RatInABox | Primary generated session (generation seed 42); latent-consistency cohort (generation seeds 42, 123, 456 and 789)[30]. | Synthetic 2D position decoding | 300 x 50 x 300 | 2D position, $R^2$ |

**Supplementary Table 2: Analysis availability and missing entries**

| Analysis | Coverage (available/eligible) | Missing entries, exclusions or eligibility notes |
|---|---|---|
| Prediction, robustness and computational cost | 112/115 | MARBLE x Allen; TNDM x Speech; SVM x Allen. |
| Latent consistency | 46/48 | MARBLE x Allen; TNDM x Speech. |
| Feature-attribution analysis | 106/115 | MARBLE for all datasets; MINT × Allen; SVM × Allen, MC PacMan and RatInABox. Reasons for |

| Analysis | Coverage (available/eligible) | Missing entries, exclusions or eligibility notes |
|---|---|---|
| | | unavailable entries are detailed in **Supplementary Note 6**. |
| Trial-valuation corrupted-trial detection analysis | 81/85 | LangevinFlow x Allen; MARBLE x Allen; TNDM x Allen; TNDM x MC PacMan. |
| Within-session corrupted-trial removal case study | 17/17 | No missing entries; eligibility was restricted to methods satisfying the fixed-output/lightweight-rescoring rule described in **Supplementary Note 5**. |
| Same-subject historical-trial selection | 16/17 | MARBLE failed during pooled cross-session graph construction before scoring. Eligibility was restricted to trial-valued decoder entries in the shared M1 channel feature space. |

Each analysis stage used an independent model fit, so a numerical failure in one stage did not determine inclusion in another. The three missing primary prediction, robustness and resource entries arose at different pipeline stages. MARBLE × Allen exceeded the 192-GB host-memory allocation during graph and embedding construction, before prediction or resource metrics were produced. TNDM × Speech returned non-finite relevant-prior/posterior KL values during variational training. SVM × Allen exceeded the 24-h allocation during exact kernel-SVC fitting on the flattened sequence-classification design. The corresponding MARBLE × Allen and TNDM × Speech latent-consistency entries were unavailable for the same reasons.

Four configured trial-valuation runs did not produce final summaries. MARBLE × Allen lacked a Stage 1 representation because graph and embedding construction exceeded the host-memory allocation. LangevinFlow × Allen exceeded the available execution time during repeated full-rank logistic-regression coalition fits. TNDM × Allen and TNDM × MC PacMan likewise exceeded their execution allocations during repeated causal-logistic and causal-linear coalition fits, respectively. These failures were confined to the subset-rescoring stage of Data Shapley.

**Supplementary Table 3: Resource and runtime conventions**

| **Resource measurement** | **Definition** |
|---|---|
| Training time | Timed model fitting, including required training-side representation extraction and downstream readout fitting where applicable. |
| Inference time | One complete pass over the held-out test split, including downstream readout prediction but excluding metric calculation. |
| Peak RAM | Maximum sampled combined resident set size (RSS) of the Python process and registered external workers. |
| Peak GPU memory | Maximum sampled device-wide GPU memory; marked not applicable when the complete measured workflow used no GPU. This applied to all GPFA, PCA, SVM and XGBoost evaluations and all MINT evaluations except RatInABox. |

Across decoders for macaque center-out reaching (**Main Fig. 2**), training time, inference time and peak RAM span 0.03–2,360 s, 0.002–13 s and 0.27–14.8 GB, respectively.

For MINT on RatInABox, the decoder remained CPU-only. Following the published MINT MC RTT construction[14], its training trajectory library used rates averaged across two GPU-trained AutoLFADS[18–20] fits. Reported training time combined MINT fitting with the longer concurrent AutoLFADS fit and rate averaging; reported peak RAM and GPU memory were the maxima across these components.

Except for the composite RatInABox MINT workflow described above, each reported model-dataset value was obtained from a single benchmark execution. Timing was measured using time.perf_counter. Inference was evaluated over the complete held-out test split: 63 macaque, 119 Allen, 33 speech, 72 MC PacMan, and 60 RatInABox trials, using each implementation's native batching configuration. Accordingly, inference times are totals for the complete test set.

Dataset loading and metric calculation were performed outside the timed regions. Timed calls included any compilation overhead triggered during execution.

Host and GPU memory usage were sampled every 0.1 s during both training and inference. Host memory was defined as the combined RSS of the Python process and any explicitly registered external workers and their descendants, including MATLAB workers used by MINT. GPU memory reflected device-wide usage on the assigned GPU.

GPU-enabled runs used a single NVIDIA H100 80 GB GPU, 24 AMD EPYC 9654 CPU cores, and approximately 188 GiB of host memory. CPU-only runs used nodes with 192 AMD EPYC 9655 CPU cores and approximately 755 GiB of host memory.

**Supplementary Table 4: Prediction representations, outputs and scoring models**

| Method or group | Representation or output used for prediction | Scoring model |
|---|---|---|
| BLEND-LFADS | LFADS factors from the neural-only student | Ridge/logistic |
| BLEND-NDT | NDT representations from the neural-only student | Ridge/logistic |
| CEBRA | CEBRA embedding | k-nearest-neighbor |
| DNN | Predicted target values or class scores | Feedforward classifier/regressor trained with benchmark targets |
| RNN, GRU and LSTM | Predicted target sequences or class scores | Recurrent classifier/regressor trained with benchmark targets |
| SVM/SVR | Spike-history features | Support-vector classifier/regressor |
| XGBoost | Spike-history features | Tree-ensemble classifier/regressor |
| DPAD | Target trajectories or class scores decoded from DPAD latent states | DPAD behavior readout (Cz) |
| GPFA | GPFA latent trajectories | Ridge/logistic |

| Method or group | Representation or output used for prediction | Scoring model |
|---|---|---|
| LangevinFlow | Predicted firing rates | Ridge/logistic |
| LDNS | Reconstructed firing rates | Ridge/logistic |
| AutoLFADS | LFADS factors | Ridge/logistic |
| MARBLE | MARBLE embeddings | Ordinary least-squares, k-nearest-neighbor or logistic |
| MINT | Predicted target trajectories | MINT trajectory-library prediction |
| NEDS | Target trajectories or class scores predicted by the transformer | Transformer model trained with benchmark targets |
| NEDS-pt | Target trajectories or class scores predicted by the transformer | Transformer model trained with benchmark targets |
| mVAE | Target trajectories or class scores predicted from neural input | mVAE prediction model |
| PCA | Principal-component coordinates | Ridge/logistic |
| SMC-RNN | Posterior-mean latent trajectories | Ridge/logistic |
| TNDM | Target predictions from task-relevant latent factors | TNDM target-prediction model |
| DFINE | Target predictions from smoothed manifold latent factors | DFINE manifold-to-target model |

**Supplementary Table 5: Consistency representations**

For the latent-consistency analysis, each included method was configured to produce a three-dimensional representation; no post hoc dimensionality reduction was applied before landmark alignment (**Fig. 3**). This common dimensionality ensured that alignment scores were compared across representations with the same number of coordinates.

| Method | Representation aligned across recordings |
|---|---|
| BLEND-LFADS | LFADS factors from the neural-only student. |

| Method | Representation aligned across recordings |
|---|---|
| CEBRA | CEBRA embedding. |
| DPAD | Behaviorally relevant DPAD latent states. |
| GPFA | GPFA latent trajectories. |
| LDNS | Autoencoder-inferred LDNS latents. |
| AutoLFADS | LFADS factors. |
| MARBLE | MARBLE embeddings. |
| mVAE | mVAE latent variables. |
| PCA | Principal-component coordinates. |
| SMC-RNN | Posterior-mean latent trajectories from the filtering posterior. |
| TNDM | Task-relevant latent factors. |
| DFINE | Smoothed manifold latent factors. |

The remaining methods were outside this grid for predefined reasons. DNN, RNN, GRU, LSTM, MINT, SVM/SVR and XGBoost were direct prediction or readout baselines without an intended latent representation for cross-recording comparison. BLEND-NDT, NEDS and NEDS-pt exposed high-dimensional neural-only or transformer representations, but not a three-dimensional latent or factor representation for matched-landmark alignment; for BLEND-NDT, the neural-only student's NDT representation width was tied to the input feature count. LangevinFlow contributed prediction scores through predicted firing rates: ridge regression decoded continuous targets from those rates, matching the Neural Latents Benchmark behavioral-decoding convention, and logistic regression decoded classification labels. Its available internal state concatenated high-dimensional Langevin position variables, Langevin velocity variables and recurrent hidden states tied to the model hidden size, so including it in a latent consistency analysis fairly would require an additional post hoc reduction choice. These method-level exclusions are distinct from missing configured **Fig. 3** cells, where MARBLE x Allen and TNDM x Speech were configured but failed.

**Supplementary Table 6: Model configurations, preprocessing and training settings**

Representation dimensions in this table refer to the primary prediction configurations; the three-dimensional configurations used for latent consistency are reported in **Supplementary Table 5**.

| **Method or group** | **Configuration, preprocessing and task adaptation** | **Optimization, training budget or stopping rule** |
|---|---|---|
| BLEND-LFADS | Raw counts; 32 factors; neural-only student used at test. | Batch size 64; teacher and student were each trained for up to 50,501 updates, with validation every 20 updates and early-stopping patience of 3,000 updates. |
| BLEND-NDT | Raw counts; representation width matched the input-feature count; neural-only student used at test. | Batch size 64; teacher and student were each trained for up to 50,501 updates, with validation every 20 updates and early-stopping patience of 3,000 updates. |
| CEBRA | 32-dimensional embedding; 32 hidden units; offset10-model, cosine distance and InfoNCE. | Batch size 512, learning rate 3e-4, Adam and 10,000 CEBRA optimization iterations. |
| DNN, RNN, GRU and LSTM | DNN: 14-bin spike history, 400 hidden units, train-set standardization and no dropout. RNN, GRU and LSTM: sequence inputs with per-neuron z-scoring from training statistics, 400 recurrent units, MSE for regression and cross-entropy for classification. | 10 epochs; DNN used Adam and recurrent baselines used RMSprop. |
| SVM/SVR | 14-bin spike history; train-set standardization. | SVC/SVR implementation from the Neural_Decoding package; C = 3.0; unrestricted solver iterations. |
| XGBoost | 14-bin spike history; tree depth 3; train-set standardization. | Learning rate 0.3 and 300 boosting rounds. |
| DPAD | 16 behaviorally relevant latent states; 50 ms smoothing for | 2,500 epochs, 20% validation split. |

| Method or group | Configuration, preprocessing and task adaptation | Optimization, training budget or stopping rule |
|---|---|---|
| | sequence regression; raw counts for classification. | |
| GPFA | Square-root-transformed counts; 10 latent dimensions; 100 ms initial Gaussian-process timescale. | 500 EM iterations. |
| LangevinFlow | One rate trace per input feature; 280 hidden units; 5% dropout; 30% coordinated dropout. | Batch size 256, learning rate 0.003, weight decay 2e-5, 2,000 epochs and 50 stochastic forward passes for predicted firing rates. |
| LDNS | One rate trace per input feature; 16 inferred latents, except 32 for attempted speech; 256 hidden channels; monkey recipe for continuous and non-speech classification; human/phoneme recipe for attempted speech. | Monkey recipe: batch size 64, 260 primary epochs and 10 warmup epochs. Human/phoneme speech recipe: batch size 256, 400 primary epochs and 20 warmup epochs. Both used learning rate 0.001, weight decay 0.01 and cosine scheduling; consistency used 200 epochs. |
| AutoLFADS | 100 factors; 200 generator units; LFADS-Torch NLB MC Maze profile. | Batch size 256; AutoLFADS PBT with 20 trials, 1,000 epochs, perturbation interval = 25, burn-in = 105 and PBT patience = 4. |
| MARBLE | Three-dimensional embeddings except 32-dimensional RatInABox position-decoding embeddings. | Batch size 64, learning rate 0.01, 120 epochs for macaque/MC PacMan motor regression; 100 epochs for RatInABox spatial regression and classification adaptations. |
| MINT | Dataset-specific causal history windows and trajectory-library smoothing; repeated one-hot target trajectories for classification. | For macaque reaching, Allen, Speech and MC PacMan, MINT trajectory-library construction was non-iterative. RatInABox followed the published MINT MC RTT construction and averaged rates from two independent training-only AutoLFADS PBT fits, with seeds 42 and 43, using the batch size, PBT-trial count, perturbation interval, burn-in and patience listed in the AutoLFADS row and an up-to-1,000-epoch budget. |

| **Method or group** | **Configuration, preprocessing and task adaptation** | **Optimization, training budget or stopping rule** |
|---|---|---|
| NEDS and NEDS-pt | Shared: 256 hidden units and 8 heads. NEDS: 5 layers, 10% temporal masking and full zero masking. NEDS-pt: 22 layers and initialization from a pretrained checkpoint. | Batch size 16, learning rate 2e-4, weight decay 0.01, 2,000 epochs and 15% learning-rate warmup. |
| mVAE | 40 latent variables; benchmark-target stream masked at test. | Batch size 32, learning rate 0.001, 25,000 primary optimization iterations and 20,000 consistency iterations; structured masking began after 5,000 iterations. |
| PCA | Full PCA basis for prediction; three dimensions for consistency; train-set scaling; no whitening. | One train-set PCA fit; no iterative stopping rule. |
| SMC-RNN | 32 latent variables; 512 recurrent units; Poisson likelihood; 64 particles. | Batch size 16, learning rate 0.001 and 2,000-epoch budget with likelihood-history early stopping (patience = 50) and alpha-floor stopping at 0.005. |
| TNDM | 2 task-relevant and 2 task-irrelevant factors; repeated one-hot labels for classification. | 1,000 epochs with validation-driven ReduceLROnPlateau; factor 0.95, patience 10, minimum learning rate $10^{-5}$ |
| DFINE | 16 manifold and 16 dynamic latent factors; training-set z-scoring of neural observations. | Batch size 32, initial learning rate 0.02 with decay factor 0.9 and 200 training epochs for all benchmark tasks. |

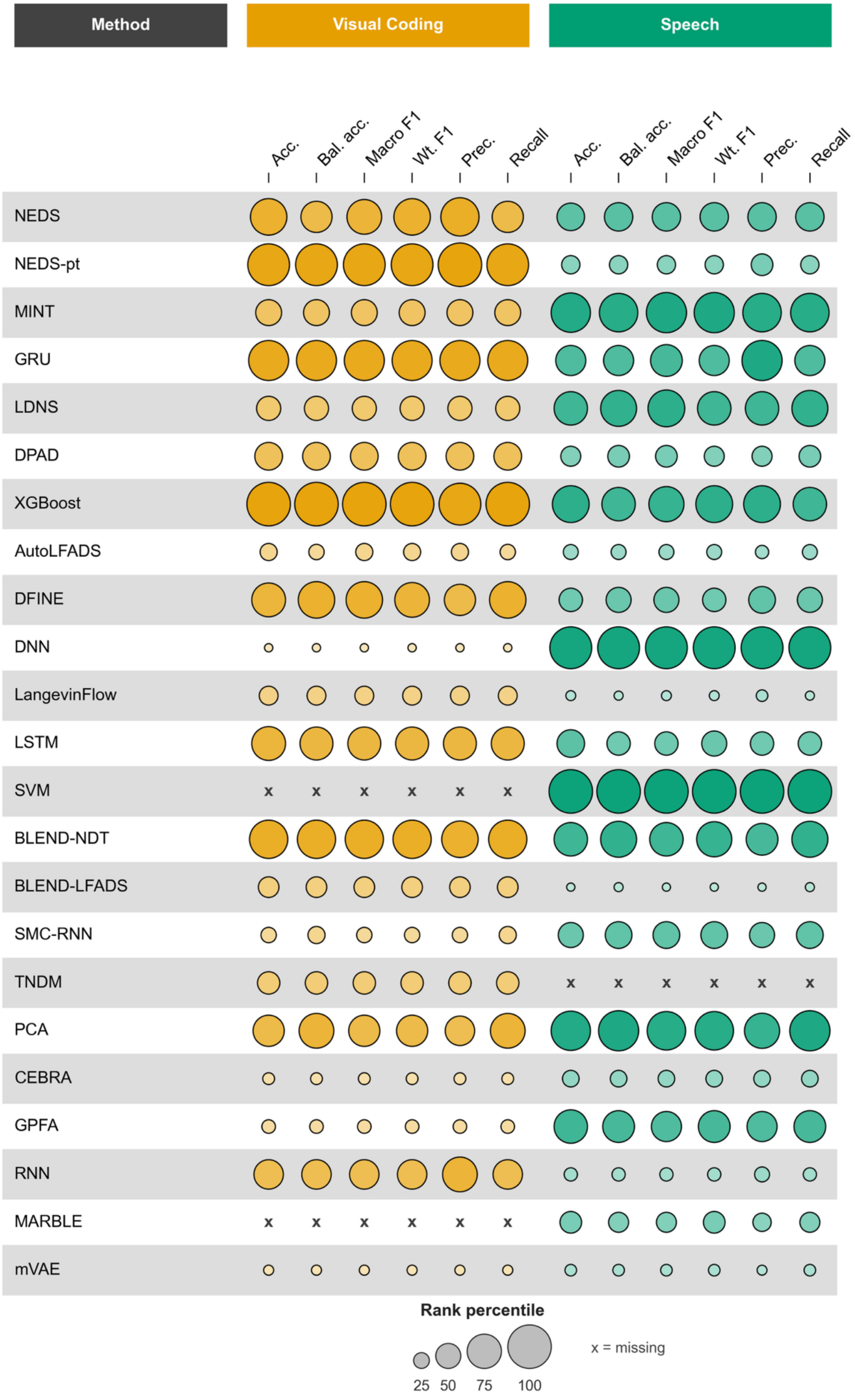
Method
Visual Coding
Speech
Acc.
Bal. acc.
Macro F1
Wt. F1
Prec.
Recall
NEDS
NEDS-pt
MINT
GRU
LDNS
DPAD
XGBoost
AutoLFADS
DFINE
DNN
LangevinFlow
LSTM
SVM
BLEND-NDT
BLEND-LFADS
SMC-RNN
TNDM
PCA
CEBRA
GPFA
RNN
MARBLE
mVAE
Rank percentile
25 50 75 100
x = missing

**Supplementary Fig. 1 | Classification rankings across scoring metrics.**

Within-dataset rank percentiles are shown for accuracy, balanced accuracy, macro F1, weighted F1, macro precision and macro recall in the Allen Neuropixels visual-coding task and attempted speech. Rows follow the mean prediction-rank order across the five primary tasks in **Fig. 2a**. Larger, darker bubbles indicate better ranks; the size legend uses the same rank-percentile mapping as the plotted bubbles. Crosses denote unavailable decoder–dataset entries.

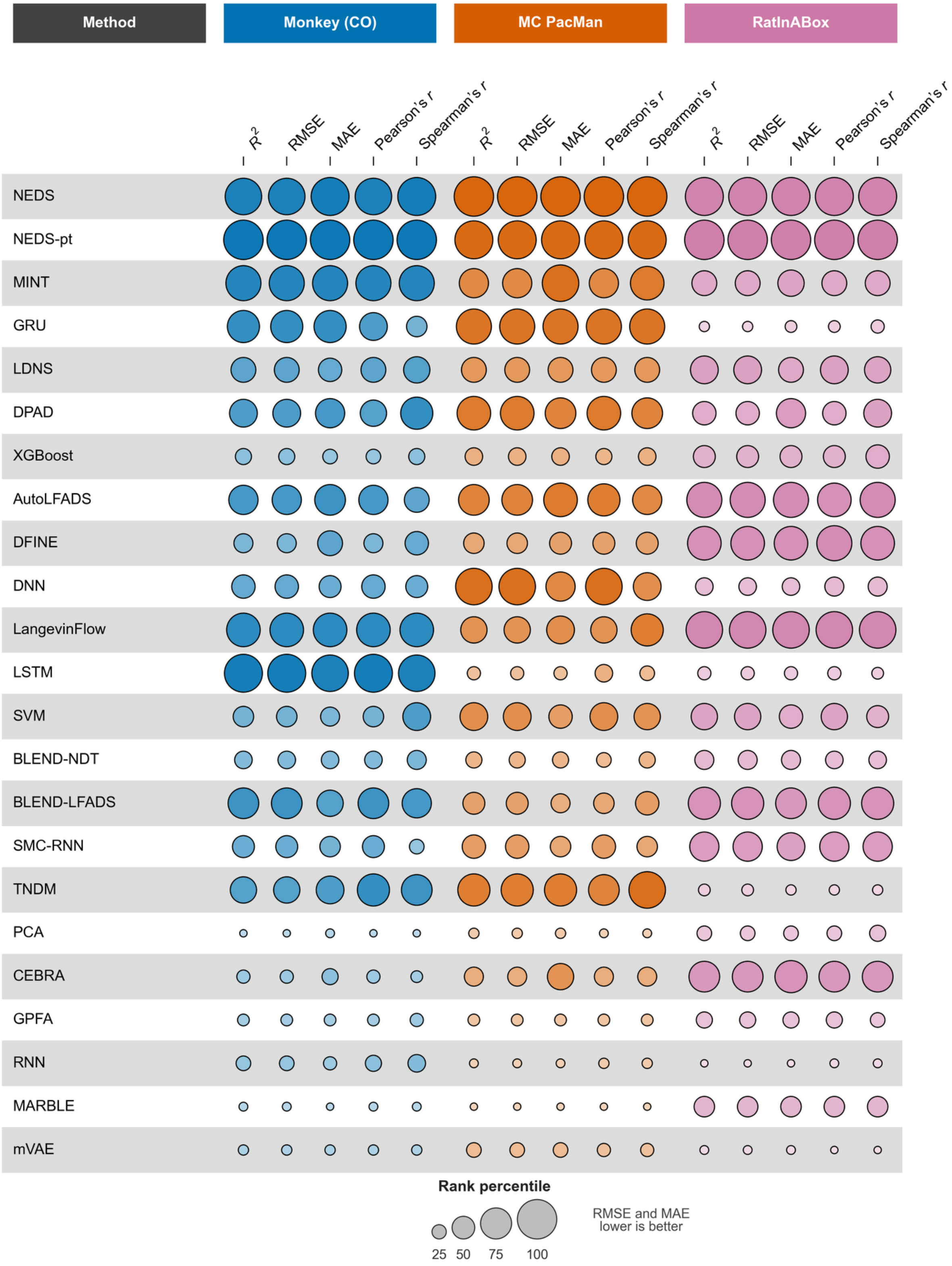

Method
Monkey (CO)
MC PacMan
RatInABox
R2
RMSE
MAE
Pearson's r
Spearman's r
NEDS
NEDS-pt
MINT
GRU
LDNS
DPAD
XGBoost
AutoLFADS
DFINE
DNN
LangevinFlow
LSTM
SVM
BLEND-NDT
BLEND-LFADS
SMC-RNN
TNDM
PCA
CEBRA
GPFA
RNN
MARBLE
mVAE
Rank percentile
25 50 75 100
RMSE and MAE lower is better

**Supplementary Fig. 2 | Regression rankings across scoring metrics.**

Within-dataset rank percentiles are shown for $R^2$, RMSE, MAE, Pearson's $r$ and Spearman's $r$ in macaque reaching, MC PacMan and RatInABox. Rows follow the mean prediction-rank order across the five primary tasks in **Fig. 2a**. Larger, darker bubbles consistently indicate better ranks, corresponding to lower RMSE and MAE and higher $R^2$ and correlation values; the size legend uses the same rank-percentile mapping as the plotted bubbles.

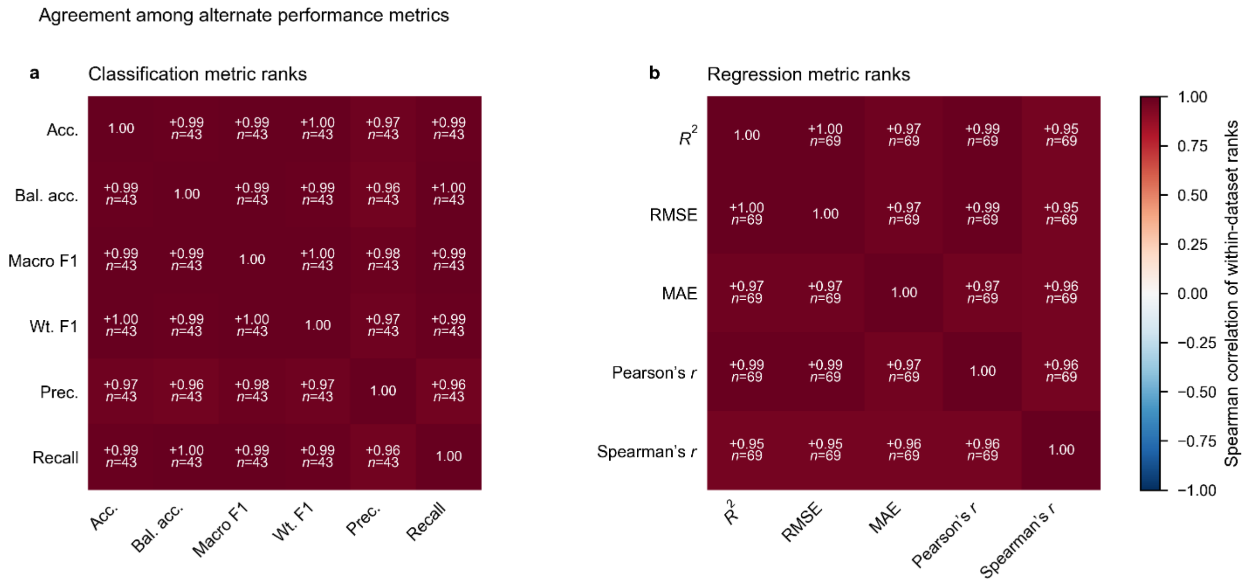


**Supplementary Fig. 3 | Agreement among prediction metrics.**

**a,** Pairwise Spearman correlations among within-dataset decoder ranks for accuracy, balanced accuracy, macro F1, weighted F1, macro precision and macro recall across the Allen Neuropixels visual-coding and attempted-speech tasks. **b,** Pairwise Spearman correlations among within-dataset decoder ranks for $R^2$, RMSE, MAE, Pearson's $r$ and Spearman's $r$ across macaque reaching, MC PacMan and RatInABox. Lower RMSE and MAE values were assigned better ranks before calculating the correlations. Off-diagonal matrix cells report Spearman's $r$ and the number of shared decoder–dataset entries, $n$; color denotes correlation strength and direction.

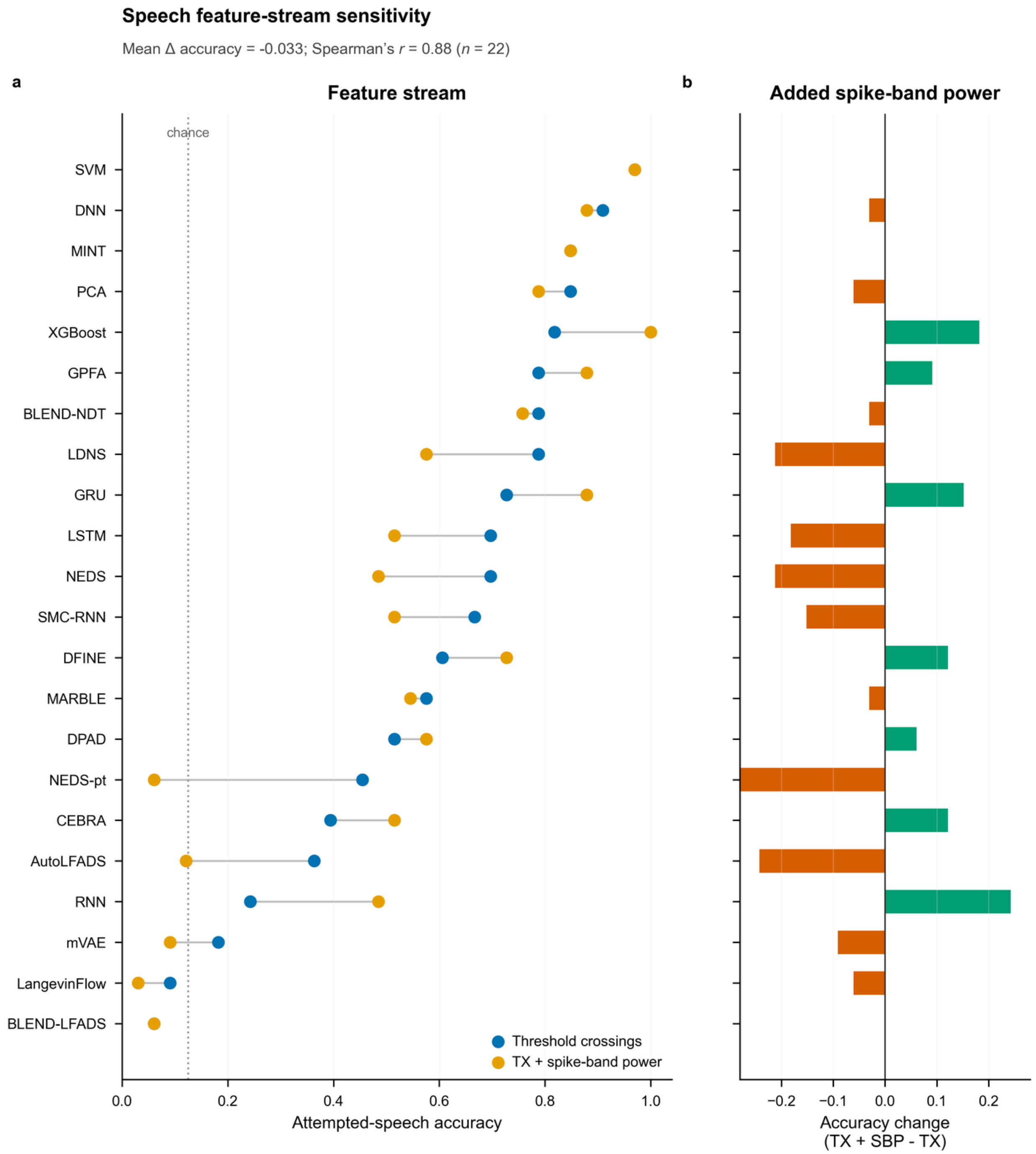


**Supplementary Fig. 4 | Attempted-speech feature-stream sensitivity.**

**a,** Classification accuracy for participant T12 using the primary threshold-crossing feature stream and a separate stream combining threshold crossings with spike-band-power-derived features. Each row represents one of the 22 decoders with results for both streams, ordered by primary-stream accuracy; lines connect paired accuracies. Blue and gold points denote the primary and combined feature streams, respectively. Both streams used the same class labels and train/test split. The dotted vertical line indicates chance accuracy for eight classes. **b,** Within-decoder accuracy change after adding spike-band-power-derived features, calculated as combined-stream accuracy minus primary-stream accuracy. Green and orange bars denote increases and decreases, respectively; zero indicates no change. The mean paired accuracy change and the Spearman correlation between decoder accuracies for the two streams are reported in the header.

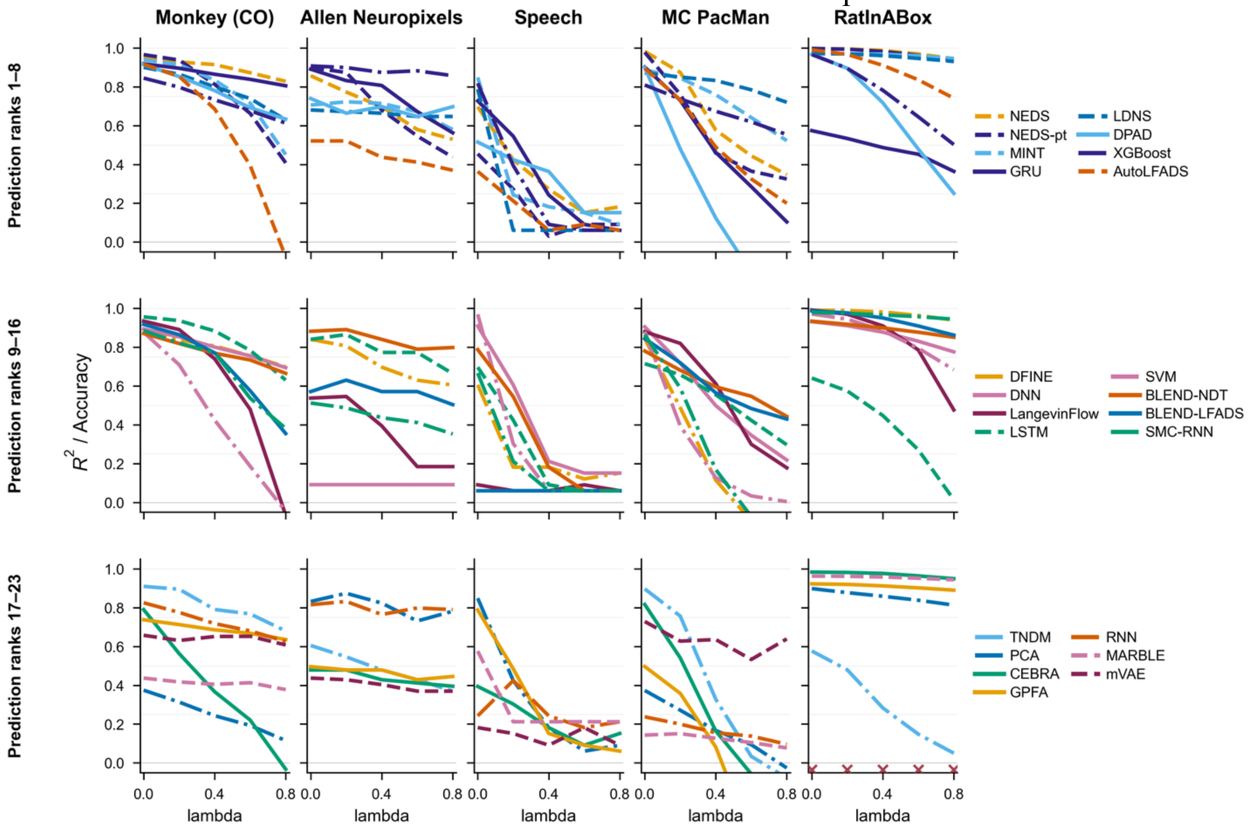


**Supplementary Fig. 5 | Robustness curves across benchmark datasets.**

Columns show held-out task performance under increasing additive Poisson count noise for macaque reaching, Allen Neuropixels, attempted speech, MC PacMan and RatInABox. Decoders follow the mean prediction-rank order in **Fig. 2a** and are divided across three rows for legibility; the rows are layout groups and do not denote decoder categories. Curves show $R^2$ for regression datasets and accuracy for classification datasets at noise levels $\lambda = 0, 0.2, 0.4, 0.6$ and $0.8$; $\lambda = 0$ is the unperturbed test set. Each decoder retains the same color and line style throughout. The raw trapezoidal area under each score-versus-$\lambda$ curve is the robustness summary reported in **Fig. 2a**. All plots use the fixed vertical range −0.05 to 1.05. Twenty-three of 560 scores from ten curves fall below the lower bound and are clipped; the minimum is $R^2 = -3.50$. Crosses at the lower boundary identify curves with negative unperturbed performance. Three unavailable decoder–dataset curves are omitted; their causes are reported in **Supplementary Table 2**.

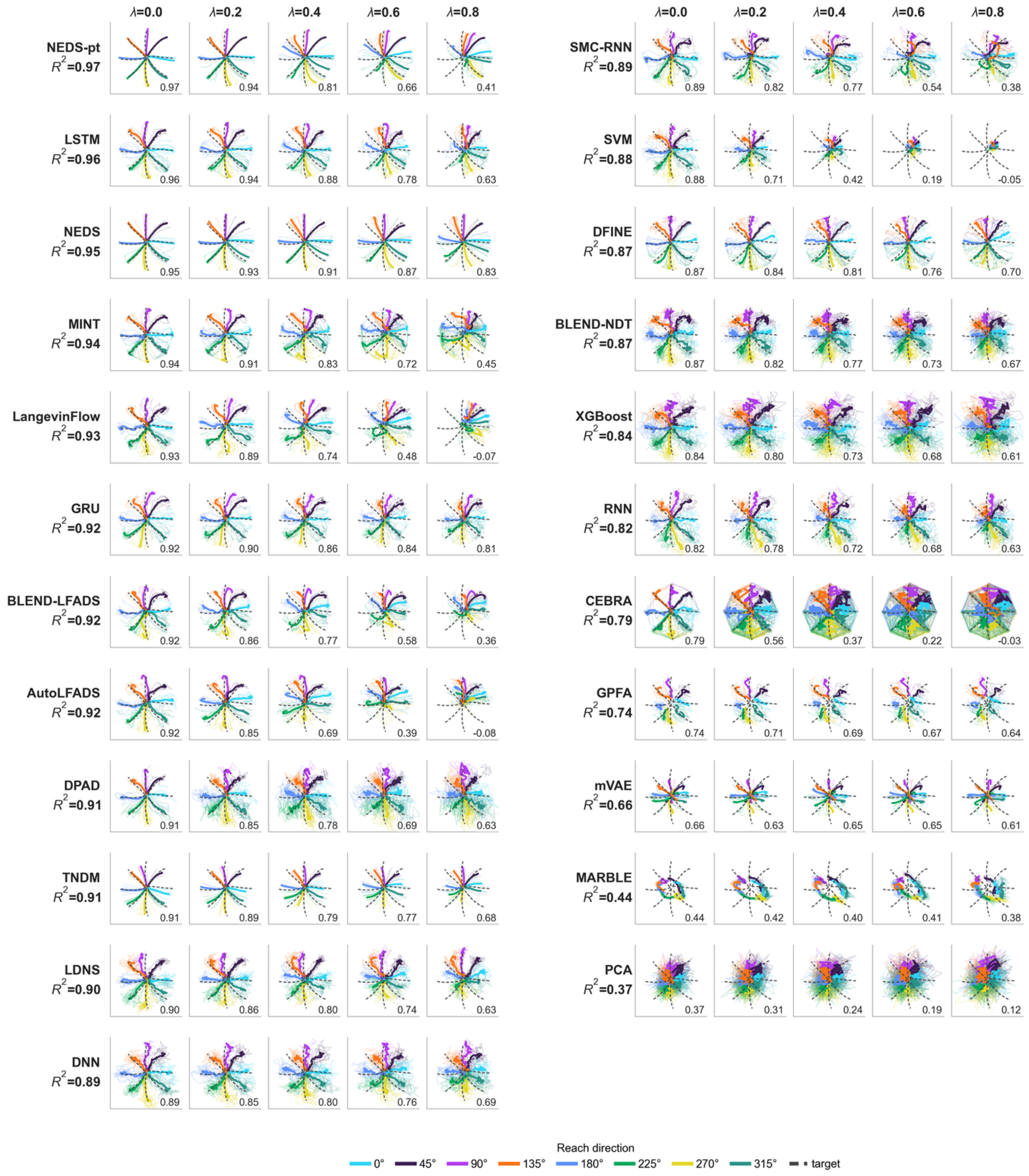


**Supplementary Fig. 6 | Macaque reaching robustness examples.**

Held-out hand-position trajectories and decoder predictions are shown at count-noise levels $\lambda$ = 0, 0.2, 0.4, 0.6 and 0.8. Decoders are ordered by unperturbed $R^2$, beginning in the left block and continuing in the right block; each block repeats the five noise-level columns. Faint colored trajectories show individual-trial predictions, solid colored trajectories show mean predictions for each reach direction and dashed gray trajectories show the corresponding mean targets. Numbers within the plots report $R^2$ at each noise level, and the value beside each decoder label is its unperturbed $R^2$.

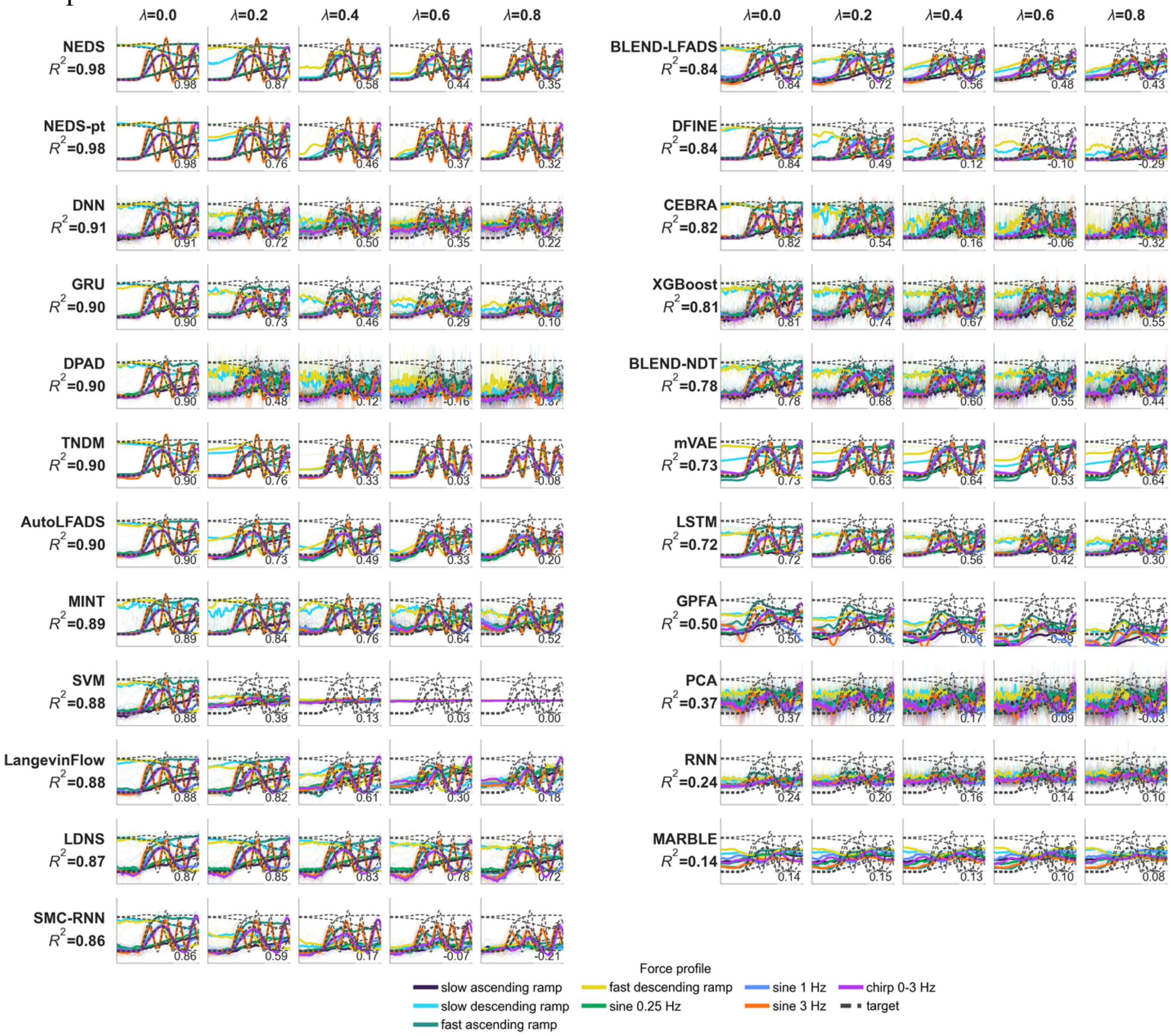


**Supplementary Fig. 7 | MC PacMan robustness examples.**

Held-out force trajectories and decoder predictions are shown at count-noise levels $\lambda = 0, 0.2, 0.4, 0.6$ and $0.8$. Decoders are ordered by unperturbed $R^2$, beginning in the left block and continuing in the right block; each block repeats the five noise-level columns. Faint colored traces show individual-trial predictions, solid colored traces show mean predictions for each force-profile condition and dashed gray traces show the corresponding mean targets. Numbers within the plots report $R^2$ at each noise level, and the value beside each decoder label is its unperturbed $R^2$.

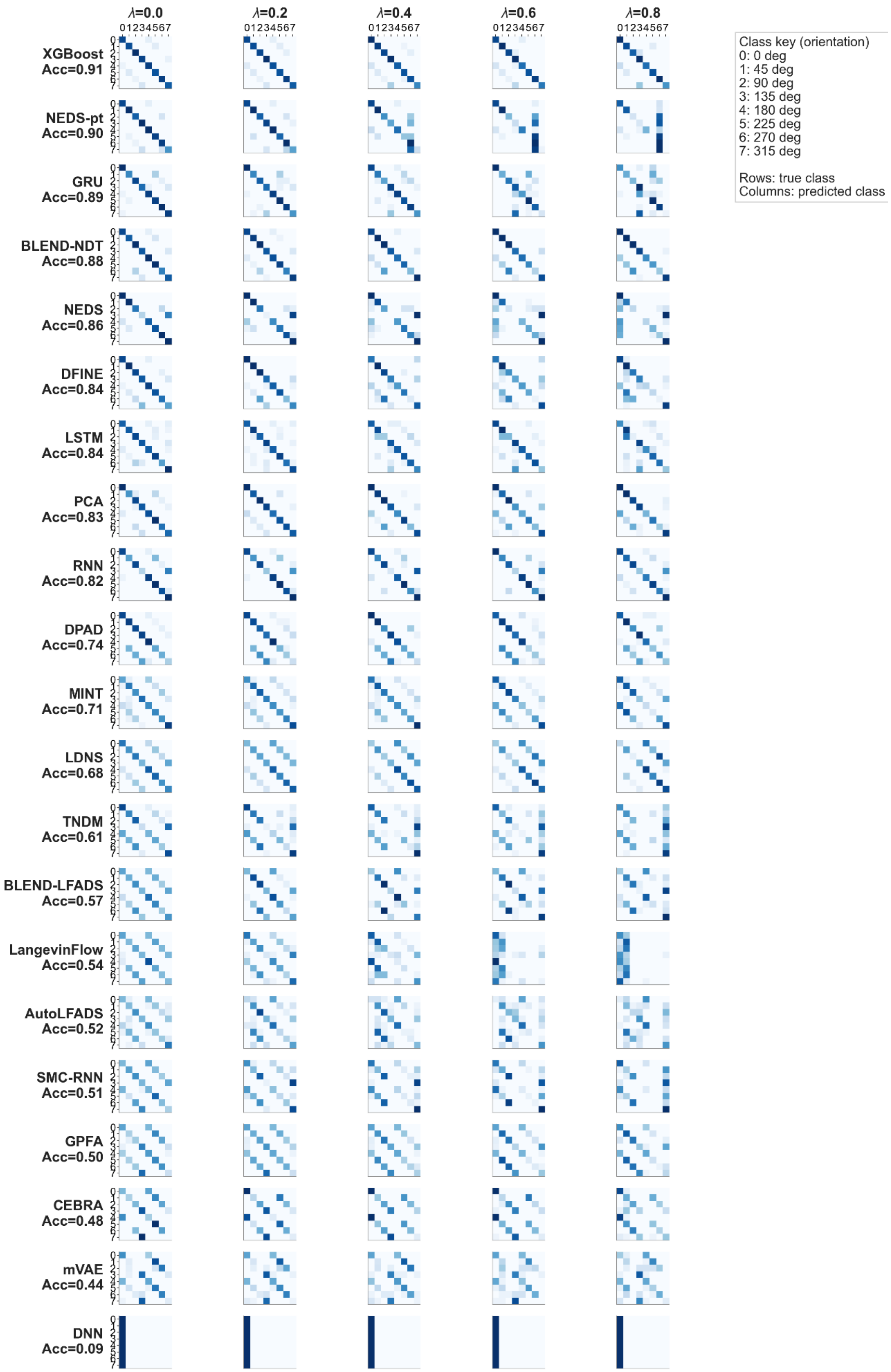
λ=0.0
λ=0.2
λ=0.4
λ=0.6
λ=0.8
XGBoost Acc=0.91
NEDS-pt Acc=0.90
GRU Acc=0.89
BLEND-NDT Acc=0.88
NEDS Acc=0.86
DFINE Acc=0.84
LSTM Acc=0.84
PCA Acc=0.83
RNN Acc=0.82
DPAD Acc=0.74
MINT Acc=0.71
LDNS Acc=0.68
TNDM Acc=0.61
BLEND-LFADS Acc=0.57
LangevinFlow Acc=0.54
AutoLFADS Acc=0.52
SMC-RNN Acc=0.51
GPFA Acc=0.50
CEBRA Acc=0.48
mVAE Acc=0.44
DNN Acc=0.09
Class key (orientation)
0: 0 deg
1: 45 deg
2: 90 deg
3: 135 deg
4: 180 deg
5: 225 deg
6: 270 deg
7: 315 deg
Rows: true class
Columns: predicted class

**Supplementary Fig. 8 | Allen Neuropixels robustness examples.**

Trial-level confusion matrices are shown at count-noise levels $\lambda$ = 0, 0.2, 0.4, 0.6 and 0.8. Available decoders are ordered by unperturbed accuracy, and noise increases across columns. Within each matrix, rows are true drifting-grating orientations and columns are predicted orientations. Each row shows the fraction of trials from that true class assigned to each predicted class and therefore sums to 1. The class key gives the orientation assigned to each integer label. Predictions within the scoring window were aggregated to one trial label by majority vote. Darker blue indicates a larger fraction of trials, and the value beside each decoder label is its unperturbed accuracy.

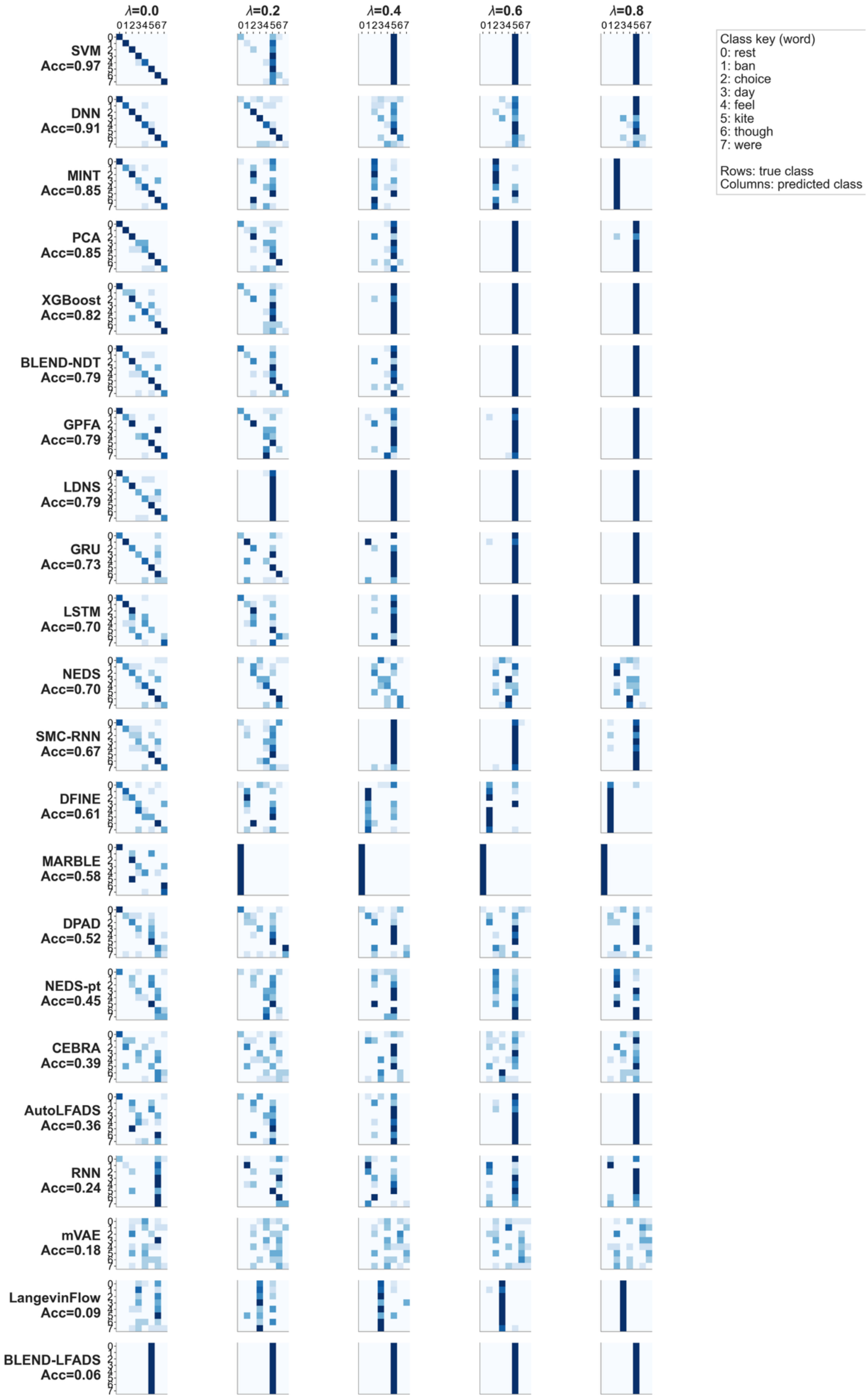
λ=0.0
λ=0.2
λ=0.4
λ=0.6
λ=0.8
SVM Acc=0.97
DNN Acc=0.91
MINT Acc=0.85
PCA Acc=0.85
XGBoost Acc=0.82
BLEND-NDT Acc=0.79
GPFA Acc=0.79
LDNS Acc=0.79
GRU Acc=0.73
LSTM Acc=0.70
NEDS Acc=0.70
SMC-RNN Acc=0.67
DFINE Acc=0.61
MARBLE Acc=0.58
DPAD Acc=0.52
NEDS-pt Acc=0.45
CEBRA Acc=0.39
AutoLFADS Acc=0.36
RNN Acc=0.24
mVAE Acc=0.18
LangevinFlow Acc=0.09
BLEND-LFADS Acc=0.06
Class key (word)
0: rest
1: ban
2: choice
3: day
4: feel
5: kite
6: though
7: were
Rows: true class
Columns: predicted class

**Supplementary Fig. 9 | Attempted-speech robustness examples.**

Trial-level confusion matrices are shown at count-noise levels $\lambda = 0, 0.2, 0.4, 0.6$ and $0.8$. Available decoders are ordered by unperturbed accuracy, and noise increases across columns. Within each matrix, rows are true classes, comprising rest and seven attempted words, and columns are predicted classes. Each row shows the fraction of trials from that true class assigned to each predicted class and therefore sums to 1. The class key maps each integer label to rest or an attempted word; rest denotes the released dataset's "Do Nothing" cue. Predictions within the scoring window were aggregated to one trial label by majority vote. Darker blue indicates a larger fraction of trials, and the value beside each decoder label is its unperturbed accuracy.

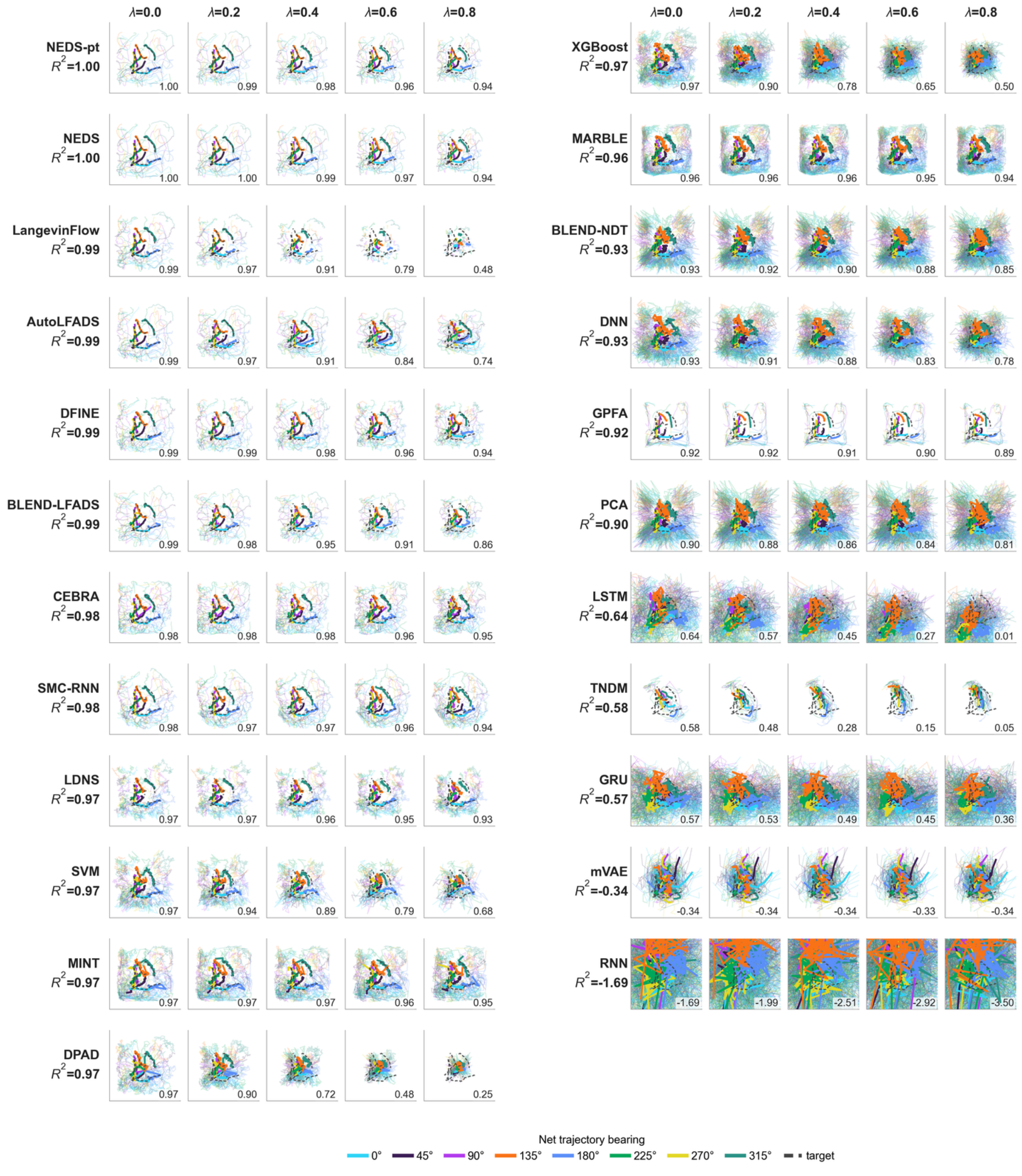


**Supplementary Fig. 10 | RatInABox robustness examples.**

Held-out two-dimensional position trajectories and decoder predictions are shown at count-noise levels $\lambda = 0, 0.2, 0.4, 0.6$ and $0.8$. Decoders are ordered by unperturbed $R^2$, beginning in the left block and continuing in the right block; each block repeats the five noise-level columns. Faint colored trajectories show individual-trial predictions, solid colored trajectories show mean predictions grouped by net target-trajectory bearing and dashed gray trajectories show the corresponding mean targets. Numbers within the plots report $R^2$ at each noise level, and the value beside each decoder label is its unperturbed $R^2$.

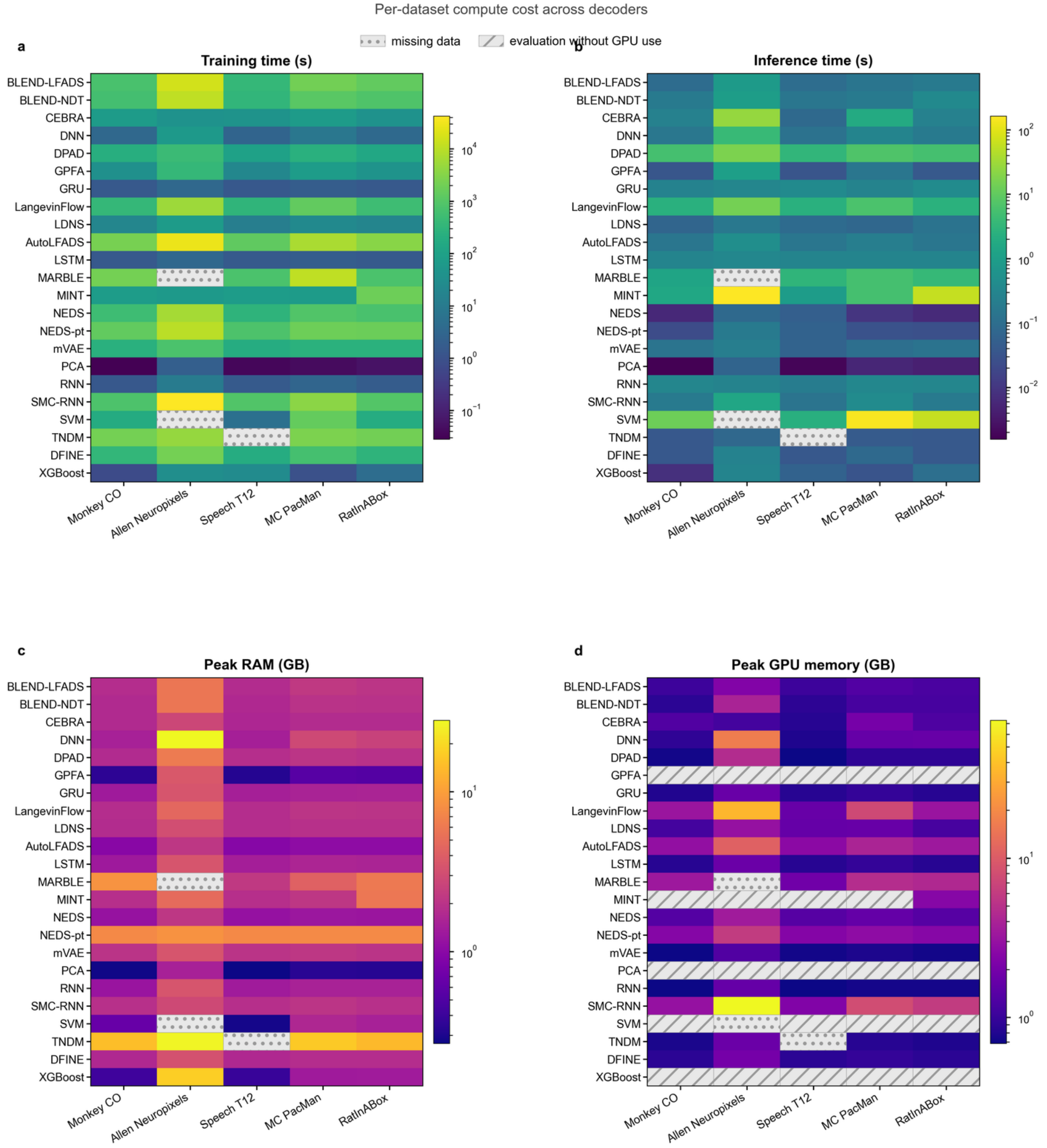


**Supplementary Fig. 11 | Computational cost measurements across datasets.**

**a,** Wall-clock training time. **b,** Wall-clock inference time. **c,** Peak host RAM. **d,** Peak GPU memory. Rows denote decoders and columns denote datasets; all color scales are logarithmic, with lower values indicating lower measured cost. Dotted cells mark unavailable decoder–dataset results. Diagonally hatched cells in d mark evaluations whose complete measured workflow used no GPU. MINT decoding itself was CPU-only; for RatInABox, the reported training time, peak RAM and peak GPU memory also include the GPU-trained AutoLFADS rate-construction step described in **Supplementary Table 3**.

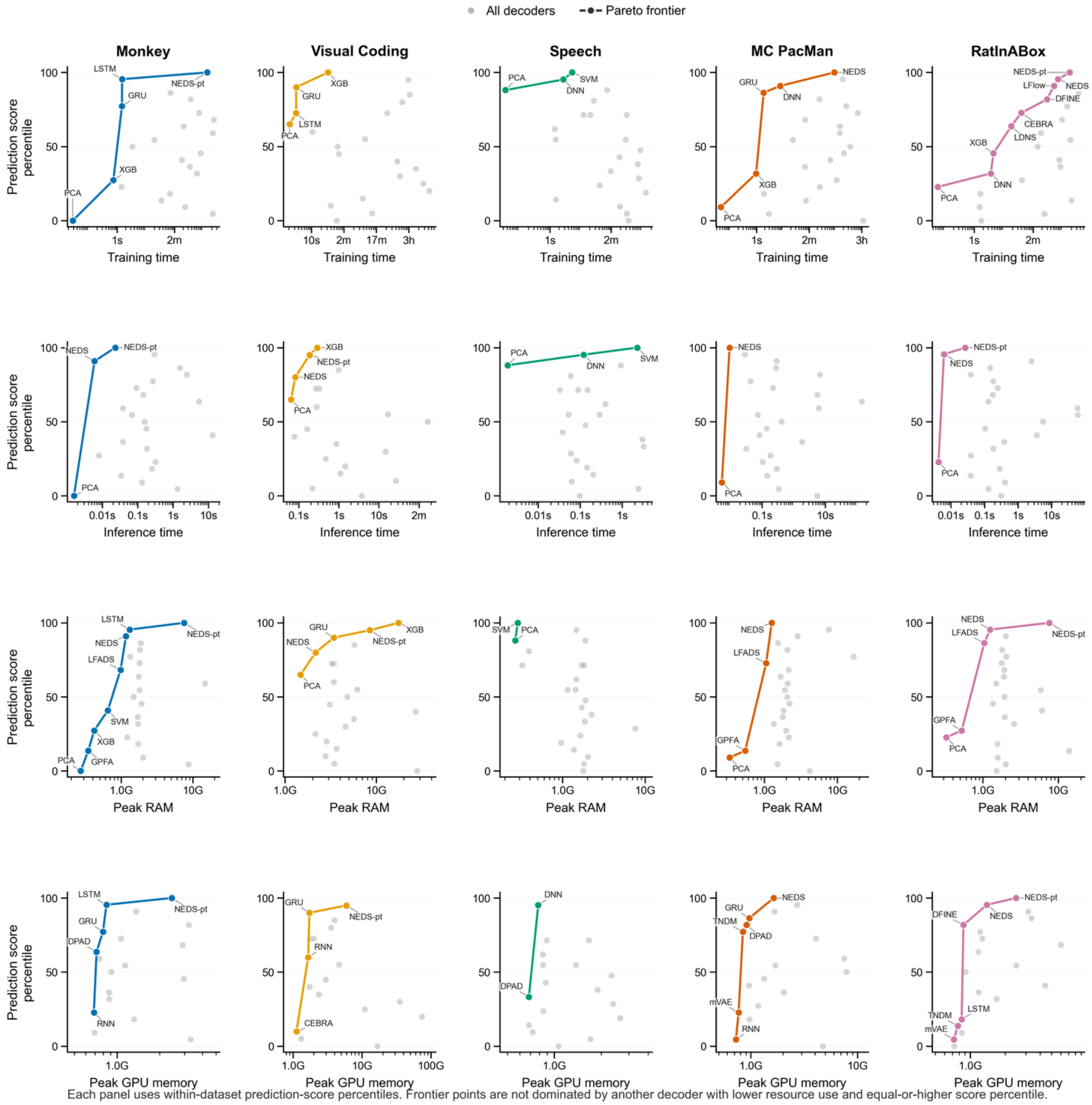


**Supplementary Fig. 12 | Prediction–resource frontiers.**

Columns denote datasets and rows show training time, inference time, peak RAM and peak GPU memory. Each point is one decoder–dataset evaluation. Gray points show all available decoders; colored points and connecting lines show the non-dominated frontier within each dataset and resource measurement. A frontier decoder has no alternative with equal or lower resource use and equal or higher prediction percentile, with at least one strict improvement. Prediction percentiles rank task performance within each dataset, and all resource axes are logarithmic. The GPU-memory row includes only evaluations whose measured workflow used a GPU. For MINT on RatInABox, the training-time, peak-RAM and peak-GPU-memory points include the AutoLFADS rate-construction step described in **Supplementary Table 3**; the inference-time point reflects the fitted MINT decoder.

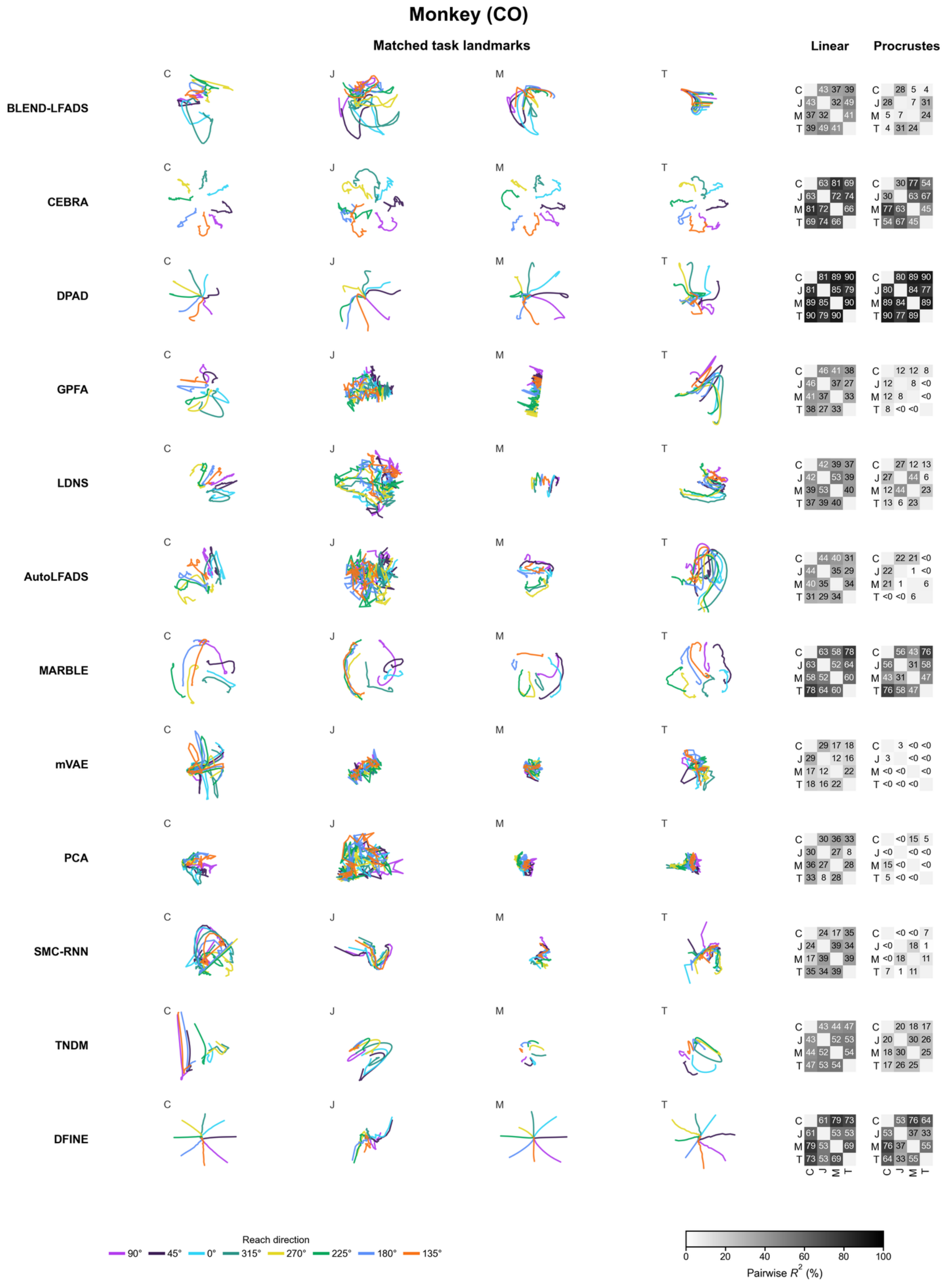

Monkey (CO)
Matched task landmarks
Linear
Procrustes
C
J
M
T
BLEND-LFADS
C 43 37 39 / J 43 32 49 / M 37 32 41 / T 39 49 41
C 28 5 4 / J 28 7 31 / M 5 7 24 / T 4 31 24
CEBRA
C 63 81 69 / J 63 72 74 / M 81 72 66 / T 69 74 66
C 30 77 54 / J 30 63 67 / M 77 63 45 / T 54 67 45
DPAD
C 81 89 90 / J 81 85 79 / M 89 85 90 / T 90 79 90
C 80 89 90 / J 80 84 77 / M 89 84 89 / T 90 77 89
GPFA
C 46 41 38 / J 46 37 27 / M 41 37 33 / T 38 27 33
C 12 12 8 / J 12 8 <0 / M 12 8 <0 / T 8 <0 <0
LDNS
C 42 39 37 / J 42 53 39 / M 39 53 40 / T 37 39 40
C 27 12 13 / J 27 44 6 / M 12 44 23 / T 13 6 23
AutoLFADS
C 44 40 31 / J 44 35 29 / M 40 35 34 / T 31 29 34
C 22 21 <0 / J 22 1 <0 / M 21 1 6 / T <0 <0 6
MARBLE
C 63 58 78 / J 63 52 64 / M 58 52 60 / T 78 64 60
C 56 43 76 / J 56 31 58 / M 43 31 47 / T 76 58 47
mVAE
C 29 17 18 / J 29 12 16 / M 17 12 22 / T 18 16 22
C 3 <0 <0 / J 3 <0 <0 / M <0 <0 <0 / T <0 <0 <0
PCA
C 30 36 33 / J 30 27 8 / M 36 27 28 / T 33 8 28
C <0 15 5 / J <0 <0 <0 / M 15 <0 <0 / T 5 <0 <0
SMC-RNN
C 24 17 35 / J 24 39 34 / M 17 39 39 / T 35 34 39
C <0 <0 7 / J <0 18 1 / M <0 18 11 / T 7 1 11
TNDM
C 43 44 47 / J 43 52 53 / M 44 52 54 / T 47 53 54
C 20 18 17 / J 20 30 26 / M 18 30 25 / T 17 26 25
DFINE
C 61 79 73 / J 61 53 53 / M 79 53 69 / T 73 53 69
C 53 76 64 / J 53 37 33 / M 76 37 55 / T 64 33 55
Reach direction
90° 45° 0° 315° 270° 225° 180° 135°
0 20 40 60 80 100
Pairwise R2 (%)

**Supplementary Fig. 13 | Macaque reaching latent-consistency visualizations.**

Rows denote methods. The four visualization columns show two-dimensional views of whitened three-dimensional, condition-averaged trajectories from macaques C, J, M and T after Procrustes alignment[31] to macaque C for display. Colors denote reach direction and each trajectory follows movement progress. The two rightmost columns show pairwise $R^2$ after intercept-free linear alignment and Procrustes alignment, respectively. Matrix rows identify the source recording and columns identify the target recording; values are percentages. Values below zero are displayed as <0, and N/A denotes an unavailable result.

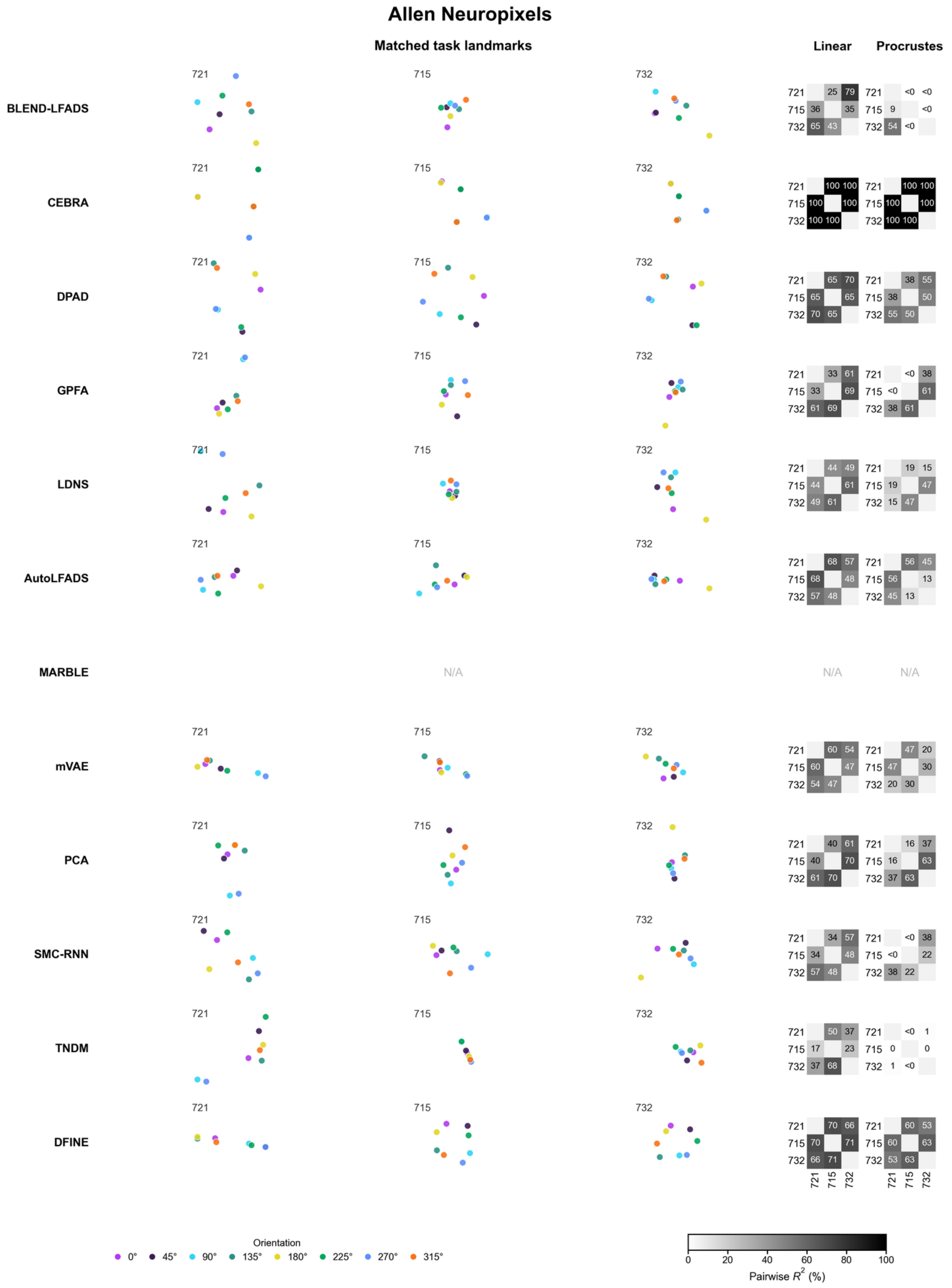

Allen Neuropixels
Matched task landmarks
Linear
Procrustes
BLEND-LFADS
CEBRA
DPAD
GPFA
LDNS
AutoLFADS
MARBLE
N/A
mVAE
PCA
SMC-RNN
TNDM
DFINE
721
715
732
Orientation
0° 45° 90° 135° 180° 225° 270° 315°
Pairwise $R^2$ (%)
0 20 40 60 80 100

**Supplementary Fig. 14 | Allen Neuropixels latent-consistency visualizations.**

Rows denote methods. The three visualization columns show two-dimensional views of whitened three-dimensional representations from sessions 721123822, 715093703 and 732592105 after Procrustes alignment to session 721123822 for display. Points are orientation-specific centroids and colors denote drifting-grating orientation. The two rightmost columns show pairwise $R^2$ after intercept-free linear alignment and Procrustes alignment, respectively. Matrix rows identify the source session and columns identify the target session; values are percentages. Values below zero are displayed as <0, and N/A denotes an unavailable result.

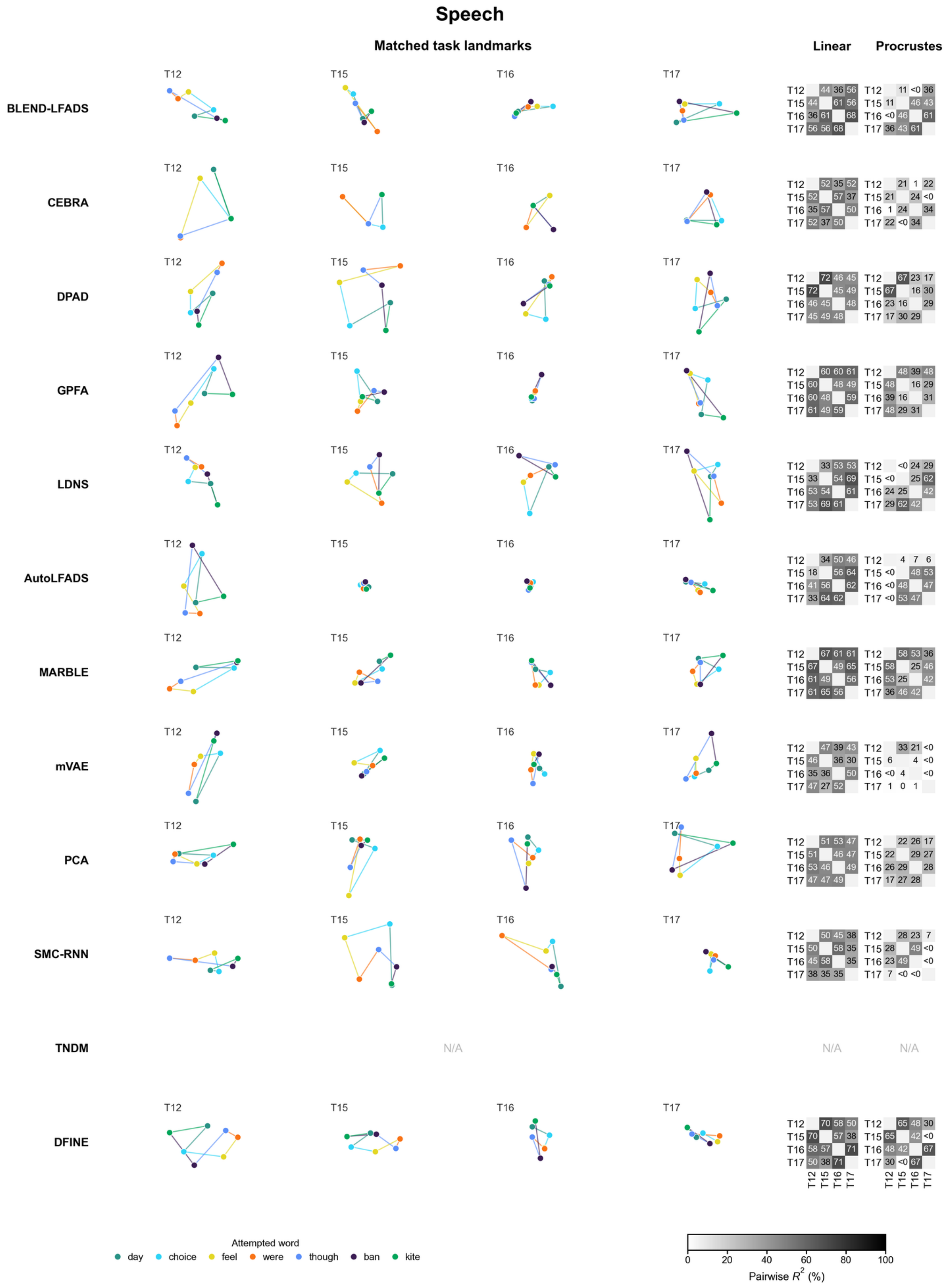
Speech
Matched task landmarks
Linear
Procrustes
BLEND-LFADS
CEBRA
DPAD
GPFA
LDNS
AutoLFADS
MARBLE
mVAE
PCA
SMC-RNN
TNDM
DFINE
T12
T15
T16
T17
N/A
BLEND-LFADS Linear: T12 44 36 56; T15 44 61 56; T16 36 61 68; T17 56 56 68
BLEND-LFADS Procrustes: T12 11 <0 36; T15 11 46 43; T16 <0 46 61; T17 36 43 61
CEBRA Linear: T12 52 35 52; T15 52 57 37; T16 35 57 50; T17 52 37 50
CEBRA Procrustes: T12 21 1 22; T15 21 24 <0; T16 1 24 34; T17 22 <0 34
DPAD Linear: T12 72 46 45; T15 72 45 49; T16 46 45 48; T17 45 49 48
DPAD Procrustes: T12 67 23 17; T15 67 16 30; T16 23 16 29; T17 17 30 29
GPFA Linear: T12 60 60 61; T15 60 48 49; T16 60 48 59; T17 61 49 59
GPFA Procrustes: T12 48 39 48; T15 48 16 29; T16 39 16 31; T17 48 29 31
LDNS Linear: T12 33 53 53; T15 33 54 69; T16 53 54 61; T17 53 69 61
LDNS Procrustes: T12 <0 24 29; T15 <0 25 62; T16 24 25 42; T17 29 62 42
AutoLFADS Linear: T12 34 50 46; T15 18 56 64; T16 41 56 62; T17 33 64 62
AutoLFADS Procrustes: T12 4 7 6; T15 <0 48 53; T16 <0 48 47; T17 <0 53 47
MARBLE Linear: T12 67 61 61; T15 67 49 65; T16 61 49 56; T17 61 65 56
MARBLE Procrustes: T12 58 53 36; T15 58 25 46; T16 53 25 42; T17 36 46 42
mVAE Linear: T12 47 39 43; T15 46 36 30; T16 35 36 50; T17 47 27 52
mVAE Procrustes: T12 33 21 <0; T15 6 4 <0; T16 <0 4 <0; T17 1 0 1
PCA Linear: T12 51 53 47; T15 51 46 47; T16 53 46 49; T17 47 47 49
PCA Procrustes: T12 22 26 17; T15 22 29 27; T16 26 29 28; T17 17 27 28
SMC-RNN Linear: T12 50 45 38; T15 50 58 35; T16 45 58 35; T17 38 35 35
SMC-RNN Procrustes: T12 28 23 7; T15 28 49 <0; T16 23 49 <0; T17 7 <0 <0
TNDM Linear: N/A
TNDM Procrustes: N/A
DFINE Linear: T12 70 58 50; T15 70 57 38; T16 58 57 71; T17 50 38 71
DFINE Procrustes: T12 65 48 30; T15 65 42 <0; T16 48 42 67; T17 30 <0 67
Attempted word
day
choice
feel
were
though
ban
kite
0
20
40
60
80
100
Pairwise R² (%)

**Supplementary Fig. 15 | Attempted-speech latent-consistency visualizations.**

Rows denote methods. The four visualization columns show two-dimensional views of whitened three-dimensional representations from participants T12, T15, T16 and T17 after Procrustes alignment to T12 for display. Colored points are centroids for the seven attempted-word classes; the "Do Nothing" class is omitted from the visualization but retained in the pairwise scores. Lines connect the displayed centroids in the same fixed order in every participant to aid visual comparison. The two rightmost columns show pairwise $R^2$ after intercept-free linear alignment and Procrustes alignment, respectively. Matrix rows identify the source participant and columns identify the target participant; values are percentages. Values below zero are displayed as <0, and N/A denotes an unavailable result.

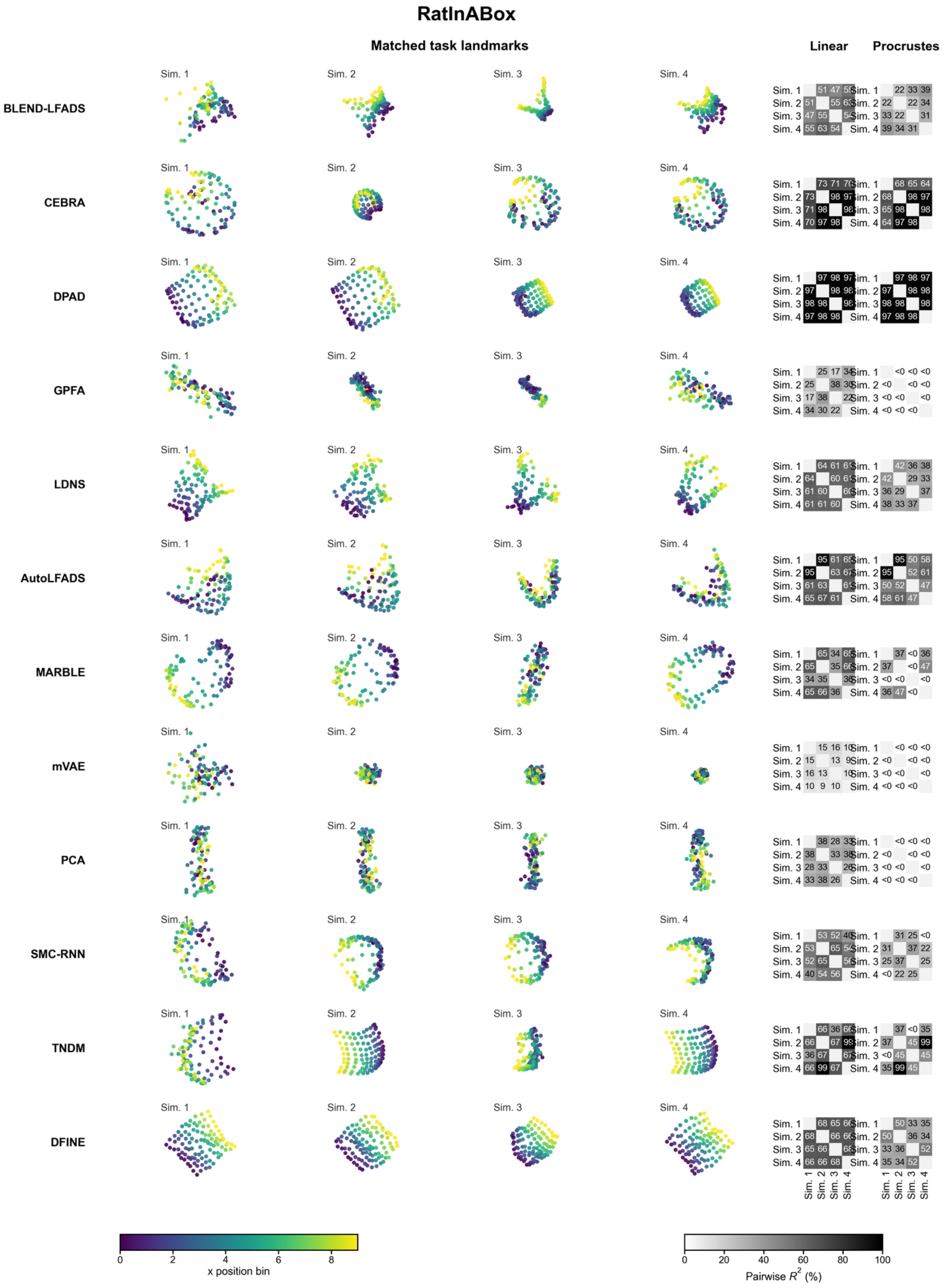
RatInABox
Matched task landmarks
Linear
Procrustes
BLEND-LFADS
CEBRA
DPAD
GPFA
LDNS
AutoLFADS
MARBLE
mVAE
PCA
SMC-RNN
TNDM
DFINE
Sim. 1
Sim. 2
Sim. 3
Sim. 4
x position bin
Pairwise $R^2$ (%)

**Supplementary Fig. 16 | RatInABox latent-consistency visualizations.**

Rows denote methods. The four visualization columns show two-dimensional views of whitened three-dimensional representations from the four synthetic sessions after Procrustes alignment to Sim. 1 for display. Points are matched spatial-landmark centroids and colors denote the landmark's $x$-position bin. The two rightmost columns show pairwise $R^2$ after intercept-free linear alignment and Procrustes alignment, respectively. Matrix rows identify the source session and columns identify the target session; values are percentages. Values below zero are displayed as <0, and N/A denotes an unavailable result.

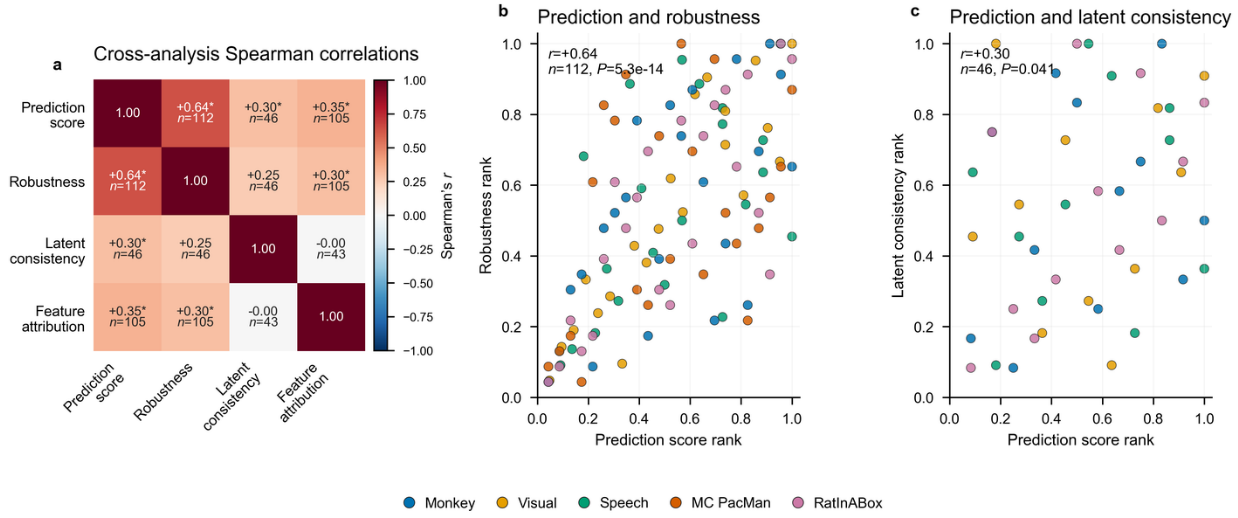


**Supplementary Fig. 17 | Cross-analysis metric correlations.**

**a,** Pairwise Spearman correlations among prediction, robustness, latent consistency and feature-attribution validation. For each measurement pair, decoder–dataset entries available for both measurements were retained, both measurements were ranked within dataset over that shared set and entries were then pooled across datasets. Off-diagonal matrix cells report Spearman's $r$ and the number of shared entries, $n$; color denotes correlation strength and direction. Asterisks indicate two-sided $P < 0.05$. **b,** Prediction percentile versus robustness percentile. **c,** Prediction percentile versus latent-consistency percentile. Each point denotes one shared decoder–dataset entry and color denotes dataset. Annotations report Spearman's $r$, the two-sided $P$ value and $n$. Sample sizes differ because each comparison includes only entries available for both measurements.

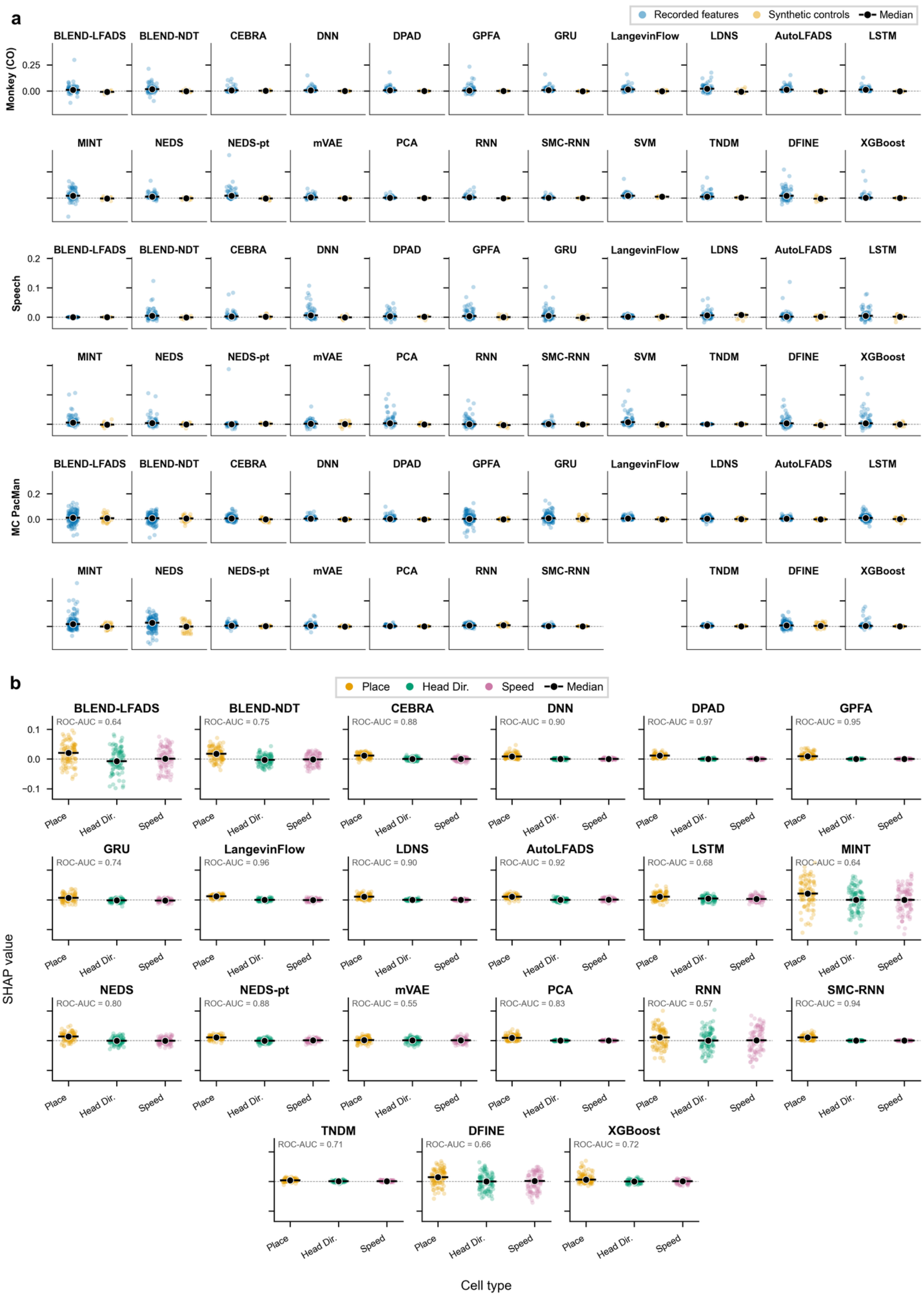
a
Recorded features
Synthetic controls
Median
Monkey (CO)
0.25
0.00
Speech
0.2
0.1
0.0
MC PacMan
0.2
0.0
BLEND-LFADS
BLEND-NDT
CEBRA
DNN
DPAD
GPFA
GRU
LangevinFlow
LDNS
AutoLFADS
LSTM
MINT
NEDS
NEDS-pt
mVAE
PCA
RNN
SMC-RNN
SVM
TNDM
DFINE
XGBoost
b
Place
Head Dir.
Speed
Median
SHAP value
0.1
0.0
−0.1
BLEND-LFADS
ROC-AUC = 0.64
BLEND-NDT
ROC-AUC = 0.75
CEBRA
ROC-AUC = 0.88
DNN
ROC-AUC = 0.90
DPAD
ROC-AUC = 0.97
GPFA
ROC-AUC = 0.95
GRU
ROC-AUC = 0.74
LangevinFlow
ROC-AUC = 0.96
LDNS
ROC-AUC = 0.90
AutoLFADS
ROC-AUC = 0.92
LSTM
ROC-AUC = 0.68
MINT
ROC-AUC = 0.64
NEDS
ROC-AUC = 0.80
NEDS-pt
ROC-AUC = 0.88
mVAE
ROC-AUC = 0.55
PCA
ROC-AUC = 0.83
RNN
ROC-AUC = 0.57
SMC-RNN
ROC-AUC = 0.94
TNDM
ROC-AUC = 0.71
DFINE
ROC-AUC = 0.66
XGBoost
ROC-AUC = 0.72
Cell type

**Supplementary Fig. 18 | Feature-attribution validation with synthetic controls and simulated cell classes.**

**a,** Per-decoder distributions of signed global Kernel SHAP values for recorded neural features and appended synthetic controls in macaque reaching, attempted speech and MC PacMan. Blue and gold points denote individual recorded features and global-rate Poisson controls, respectively; black points and horizontal bars denote medians. **b,** Per-decoder signed global Kernel SHAP values for RatInABox inputs grouped as place, head-direction and speed cells. Colored points denote individual cells and black points and horizontal bars denote medians. Annotations report ROC-AUC for ranking place cells above head-direction and speed cells; chance performance is 0.5. Positive and negative values are retained and quantify attributed contributions to held-out task utility under the benchmark masking convention.

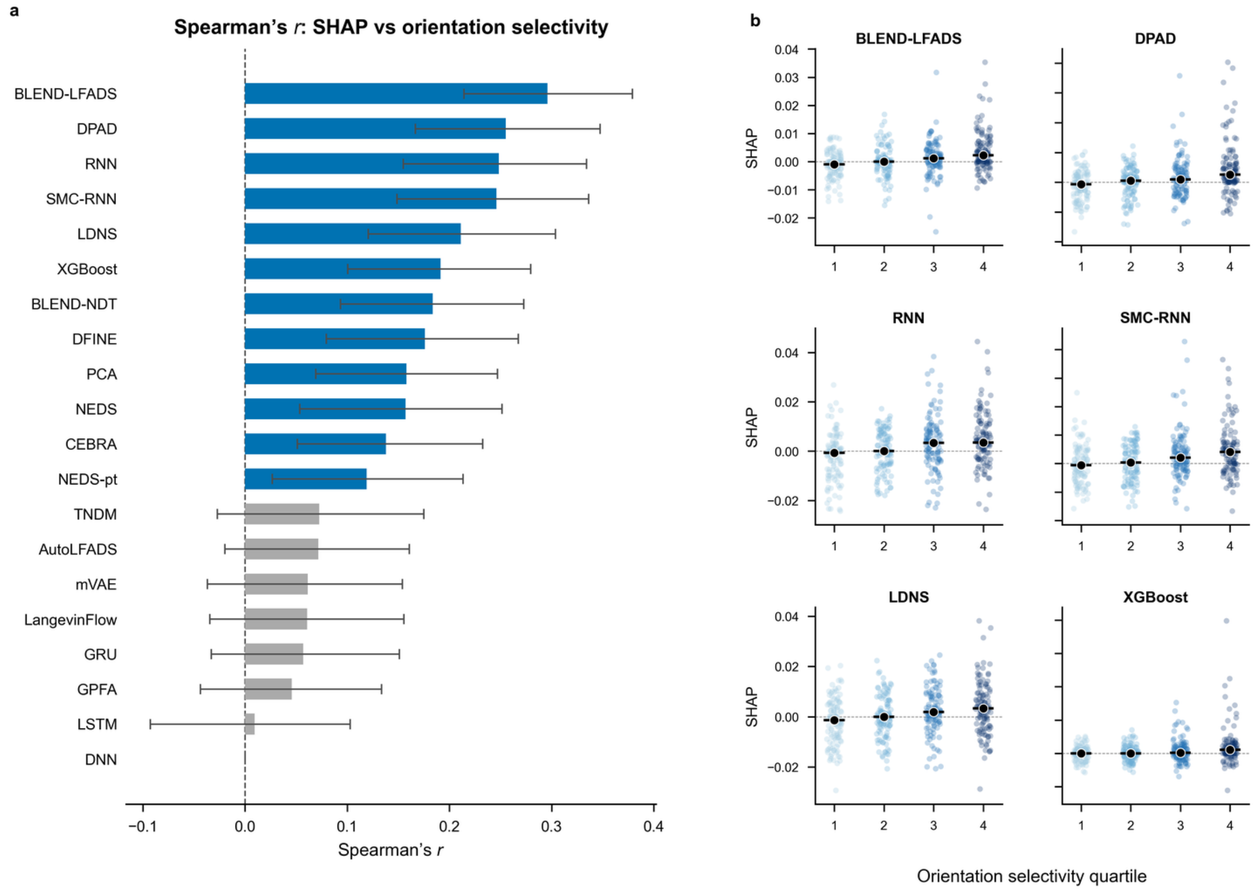


**Supplementary Fig. 19 | Allen orientation-selectivity validation.**

**a,** Spearman correlation between each unit's signed global Kernel SHAP value and drifting-grating global orientation selectivity for each decoder. Bars are ordered by Spearman's *r*; error bars show pointwise 95% bootstrap percentile intervals from 1,000 resamples of the 444 units. Blue bars denote intervals excluding zero and gray bars denote intervals including zero. DNN's constant attribution vector was scored as zero agreement (**Methods**). **b,** Signed SHAP values for the six decoders with the largest correlations in a, grouped by orientation-selectivity quartile from lowest to highest. Points denote individual units and black points and horizontal bars denote medians.

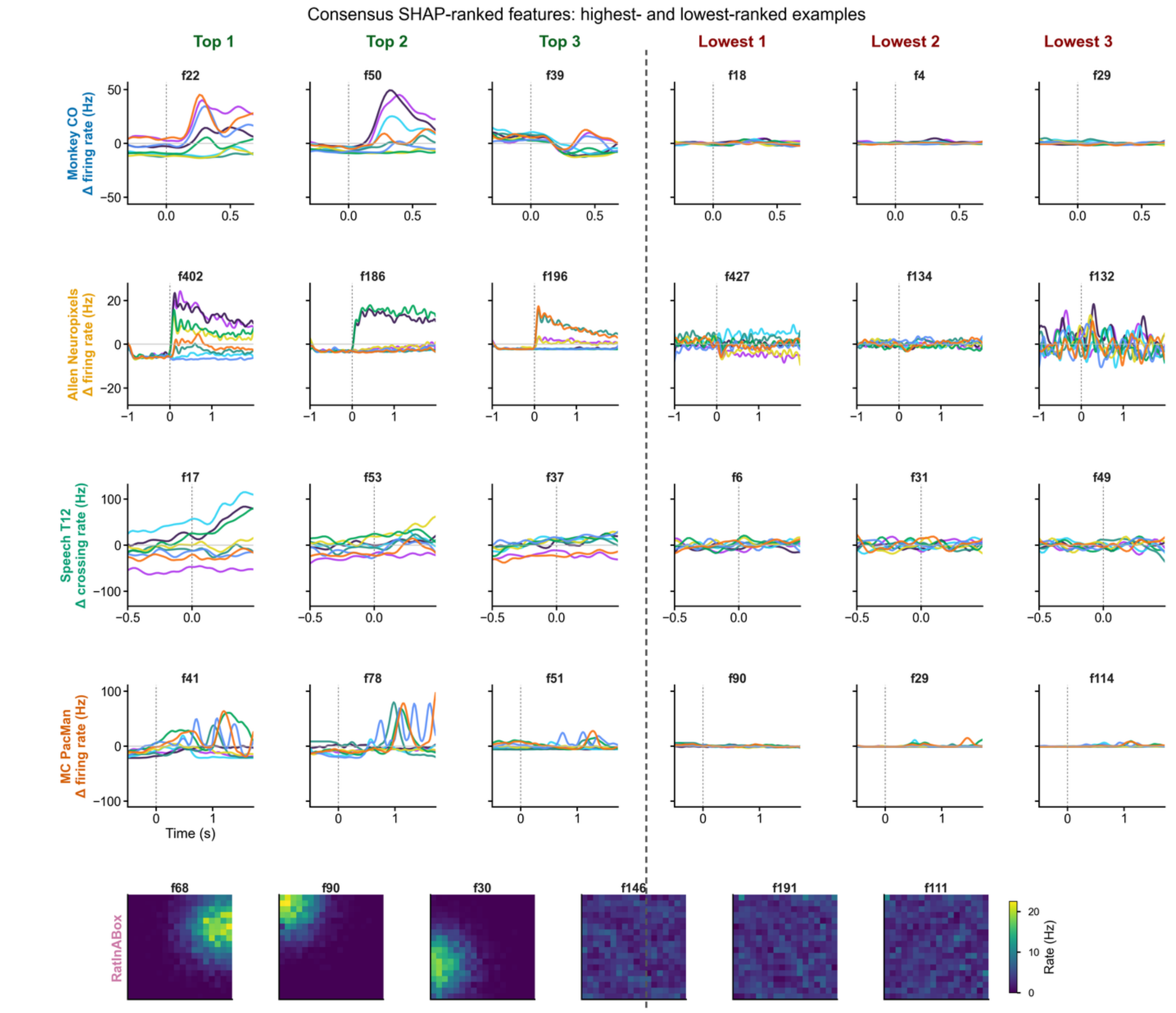


Colored lines show condition-averaged neural-event-rate profiles for reach direction, visual orientation, attempted-speech class or MC PacMan force-profile condition; RatInABox panels show spatial rate maps.

**Supplementary Fig. 20 | Highest- and lowest-ranked feature examples.**

Rows denote datasets; the first three columns show the highest-ranked features and the final three columns show the lowest-ranked features under each dataset's consensus signed global Kernel SHAP ranking. Consensus ranks average each feature's within-decoder rank across qualifying attribution solutions. Qualification required ROC-AUC ≥ 0.55 for the synthetic-control and RatInABox comparisons or Spearman's $r \geq 0$ for Allen; constant attribution vectors were excluded. Macaque reaching, Allen Neuropixels and MC PacMan show condition-averaged firing rates after subtracting each feature's grand mean; attempted speech shows similarly centered threshold-crossing event rates. Colors denote reach direction, visual orientation, attempted-speech class and force-profile condition, respectively. Temporal profiles were smoothed for display using a Gaussian kernel with a 35-ms standard deviation. Dotted vertical lines mark the start of each dataset's evaluated window. RatInABox shows spatial firing-rate maps. Panel labels give feature indices. Across the displayed examples, the highest-ranked features generally show more pronounced condition-dependent temporal modulation or spatial localization than the lowest-ranked features. SHAP values determine the rankings; the panels show the corresponding neural firing profiles.

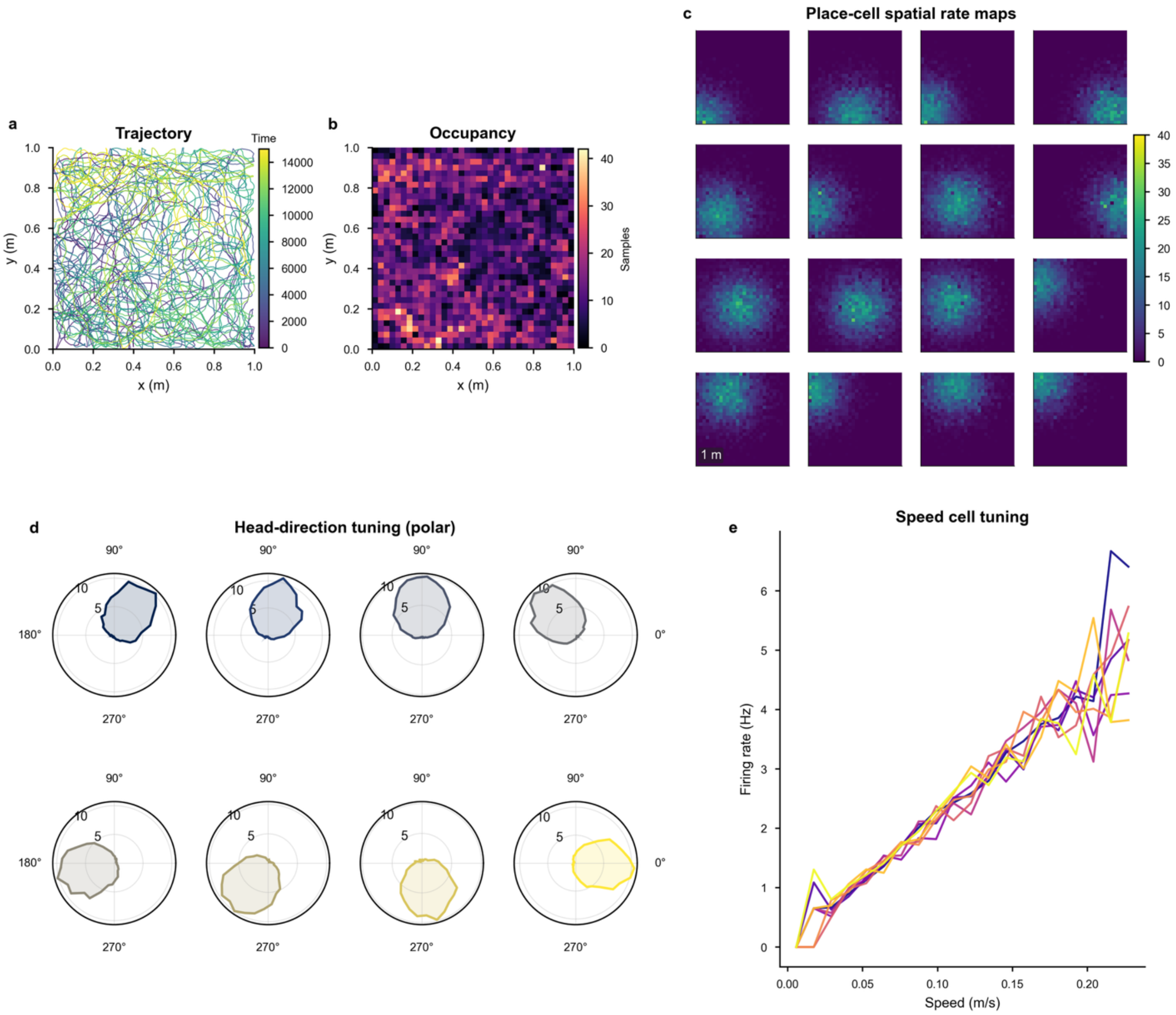


**Supplementary Fig. 21 | RatInABox cell-class atlas.**

**a,** Agent trajectory through the one-square-meter arena, colored by time. **b,** Spatial occupancy measured as the number of samples in each position bin. **c,** Spatial firing-rate maps for 16 place cells, shown with a common rate scale. **d,** Polar head-direction tuning curves for eight head-direction cells. **e,** Firing rate as a function of running speed for eight speed cells; colors denote individual cells. Cell examples in c–e were randomly selected with seed 42 from the canonical RatInABox benchmark session.

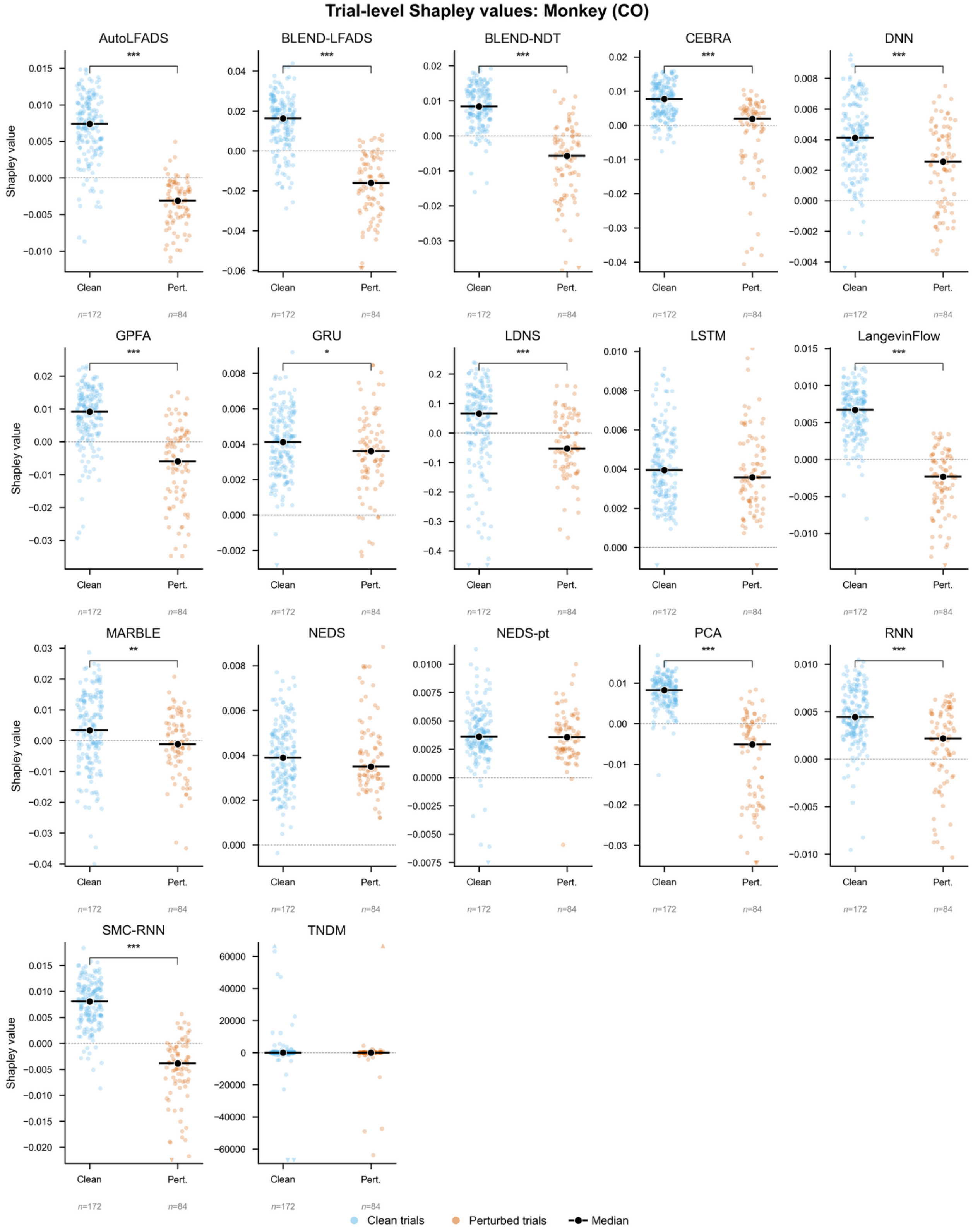


Stars: one-sided Mann–Whitney $U$ test, clean > perturbed (* $P < 0.05$, ** $P < 0.01$, *** $P < 0.001$). Y-limits are panel-specific; triangles mark clipped tails.

**Supplementary Fig. 22 | Macaque reaching trial-valuation summaries.**

Each decoder subplot shows signed trial values from the 75° full-subspace rotation analysis. Points denote unperturbed training trials (blue; $n = 172$ per decoder) and trials corrupted by rotating neural population activity while leaving targets unchanged (orange; $n = 84$ per decoder). Black points and horizontal bars indicate medians. Seventeen evaluated decoders are shown. Vertical limits are set separately for each decoder; triangles at the upper or lower boundary denote observations beyond the displayed range. Asterisks indicate $P$ values from one-sided Mann–Whitney $U$ tests of higher trial values in unperturbed than corrupted trials (*$P < 0.05$; **$P < 0.01$; ***$P < 0.001$).

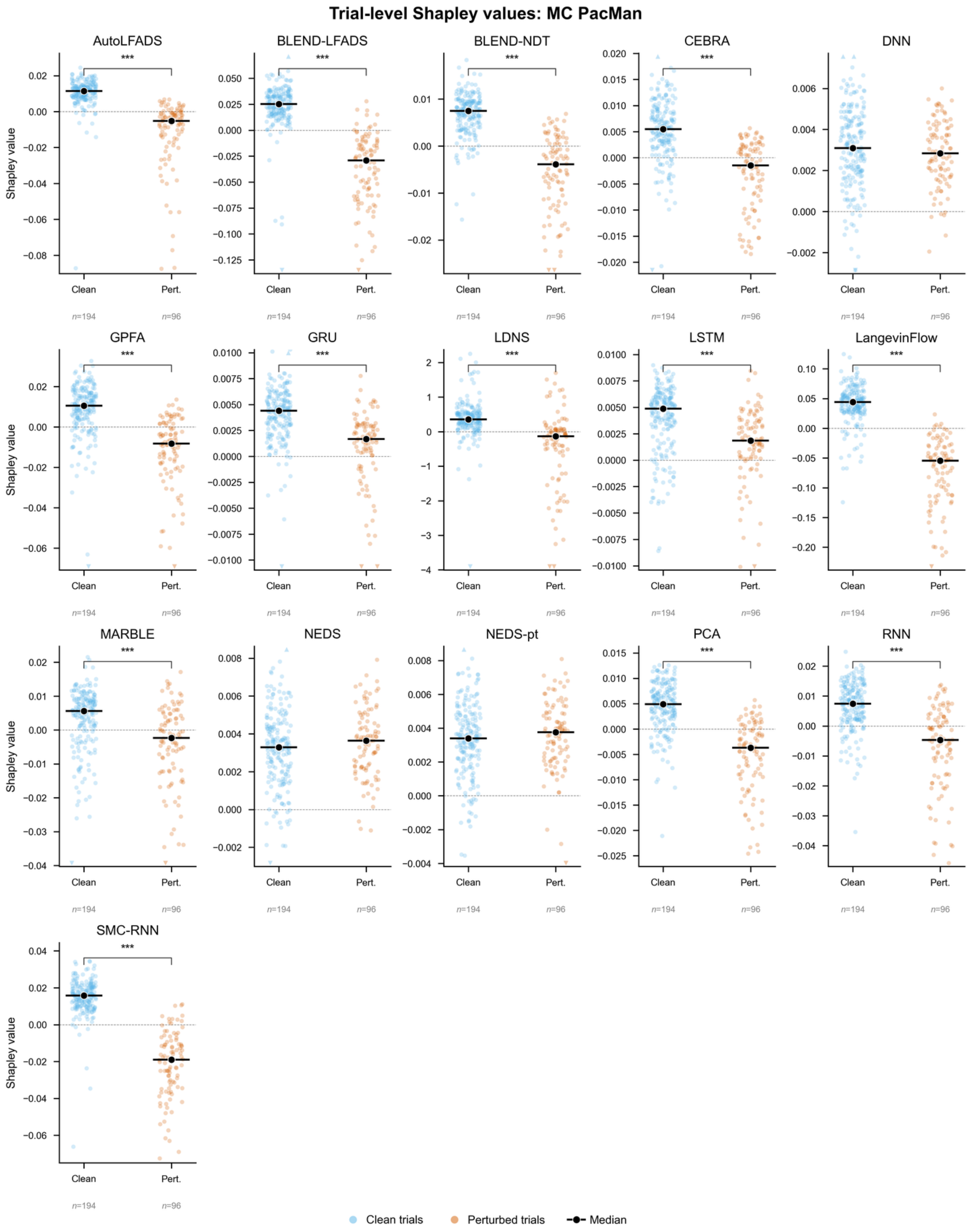




Stars: one-sided Mann–Whitney $U$ test, clean > perturbed (* $P < 0.05$, ** $P < 0.01$, *** $P < 0.001$). Y-limits are panel-specific; triangles mark clipped tails.

**Supplementary Fig. 23 | MC PacMan trial-valuation summaries.**

Each decoder subplot shows signed trial values from the 75° full-subspace rotation analysis. Points denote unperturbed training trials (blue; $n$ = 194 per decoder) and trials corrupted by rotating neural population activity while leaving targets unchanged (orange; $n$ = 96 per decoder). Black points and horizontal bars indicate medians. Sixteen evaluated decoders are shown; the unavailable entry and its cause are reported in **Supplementary Table 2**. Vertical limits are set separately for each decoder; triangles at the upper or lower boundary denote observations beyond the displayed range. Asterisks indicate $P$ values from one-sided Mann–Whitney $U$ tests of higher trial values in unperturbed than corrupted trials (*$P < 0.05$; **$P < 0.01$; ***$P < 0.001$).

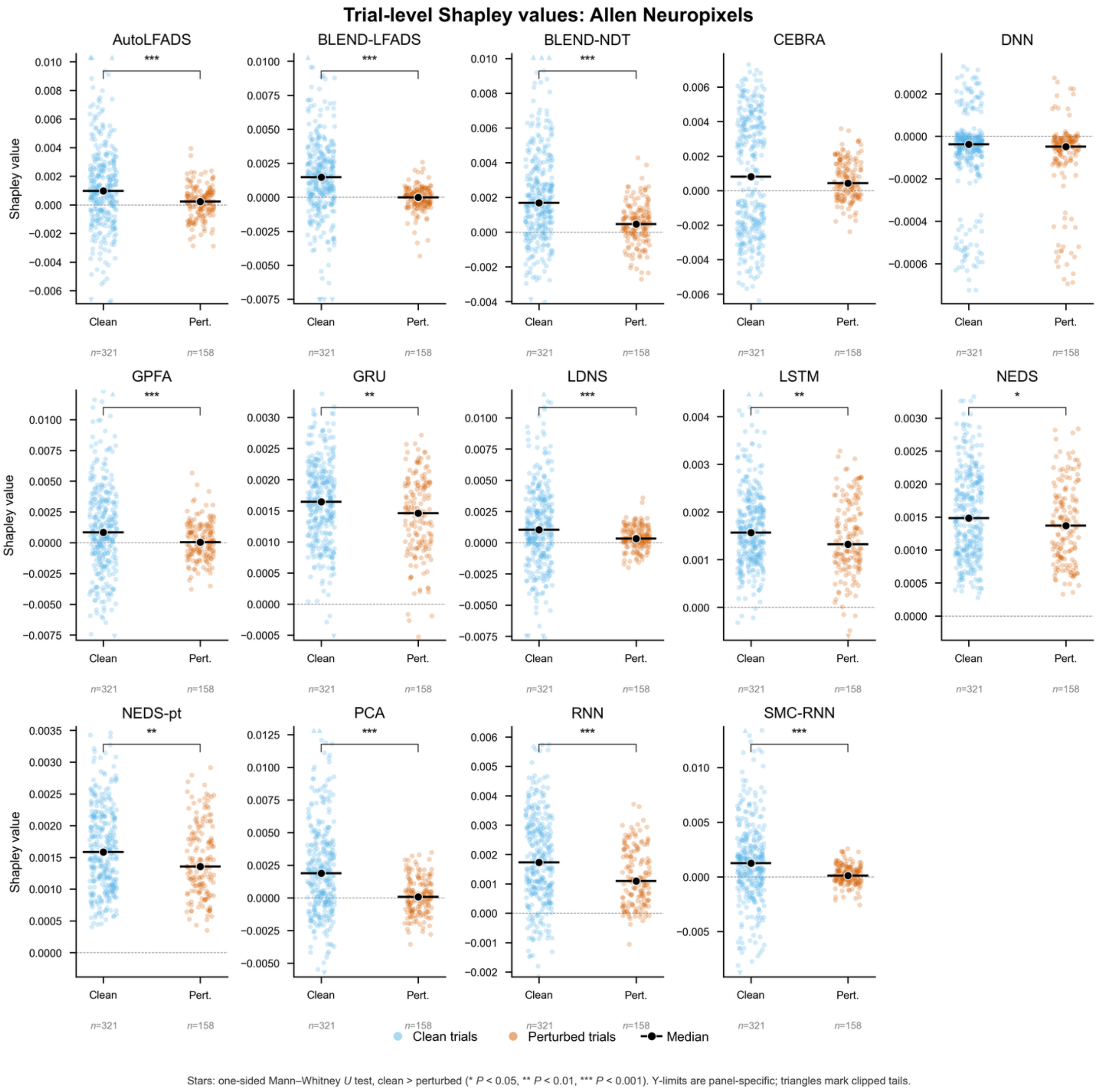

**Supplementary Fig. 24 | Allen Neuropixels trial-valuation summaries.**

Each decoder subplot shows signed trial values from the 75° full-subspace rotation analysis. Points denote unperturbed training trials (blue; $n = 321$ per decoder) and trials corrupted by rotating neural population activity while leaving class labels unchanged (orange; $n = 158$ per decoder). Black points and horizontal bars indicate medians. Fourteen evaluated decoders are shown; the unavailable entries and their causes are reported in **Supplementary Table 2**. Vertical limits are set separately for each decoder; triangles at the upper or lower boundary denote observations beyond the displayed range. Asterisks indicate $P$ values from one-sided Mann–Whitney $U$ tests of higher trial values in unperturbed than corrupted trials (*$P < 0.05$; **$P < 0.01$; ***$P < 0.001$).

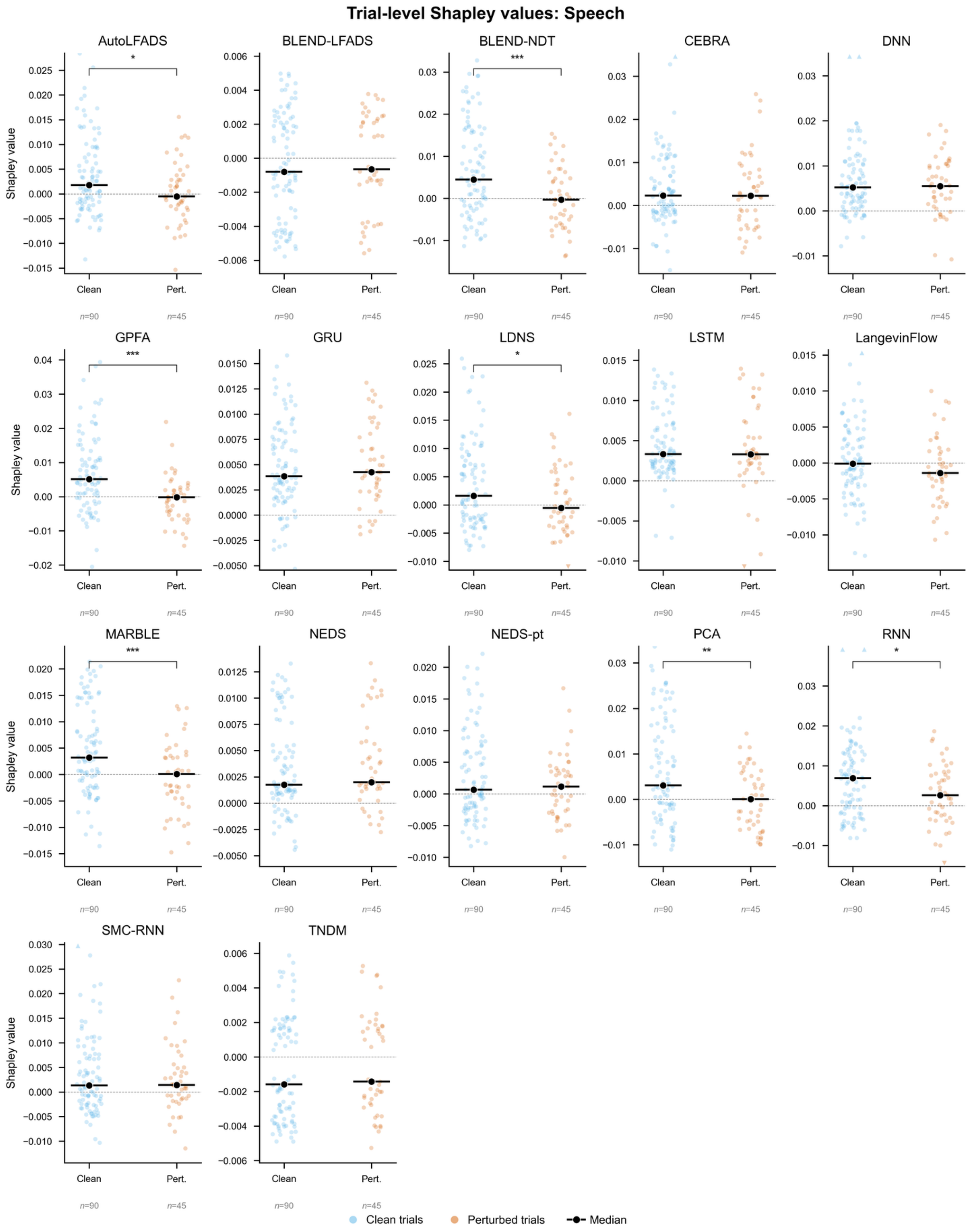


Stars: one-sided Mann–Whitney $U$ test, clean > perturbed (* $P < 0.05$, ** $P < 0.01$, *** $P < 0.001$). Y-limits are panel-specific; triangles mark clipped tails.

623

**Supplementary Fig. 25 | Attempted-speech trial-valuation summaries.**

Each decoder subplot shows signed trial values from the 75° full-subspace rotation analysis. Points denote unperturbed training trials (blue; $n = 90$ per decoder) and trials corrupted by rotating neural population activity while leaving class labels unchanged (orange; $n = 45$ per decoder). Black points and horizontal bars indicate medians. Seventeen evaluated decoders are shown. Vertical limits are set separately for each decoder; triangles at the upper or lower boundary denote observations beyond the displayed range. Asterisks indicate $P$ values from one-sided Mann–Whitney $U$ tests of higher trial values in unperturbed than corrupted trials (*$P < 0.05$; **$P < 0.01$; ***$P < 0.001$).

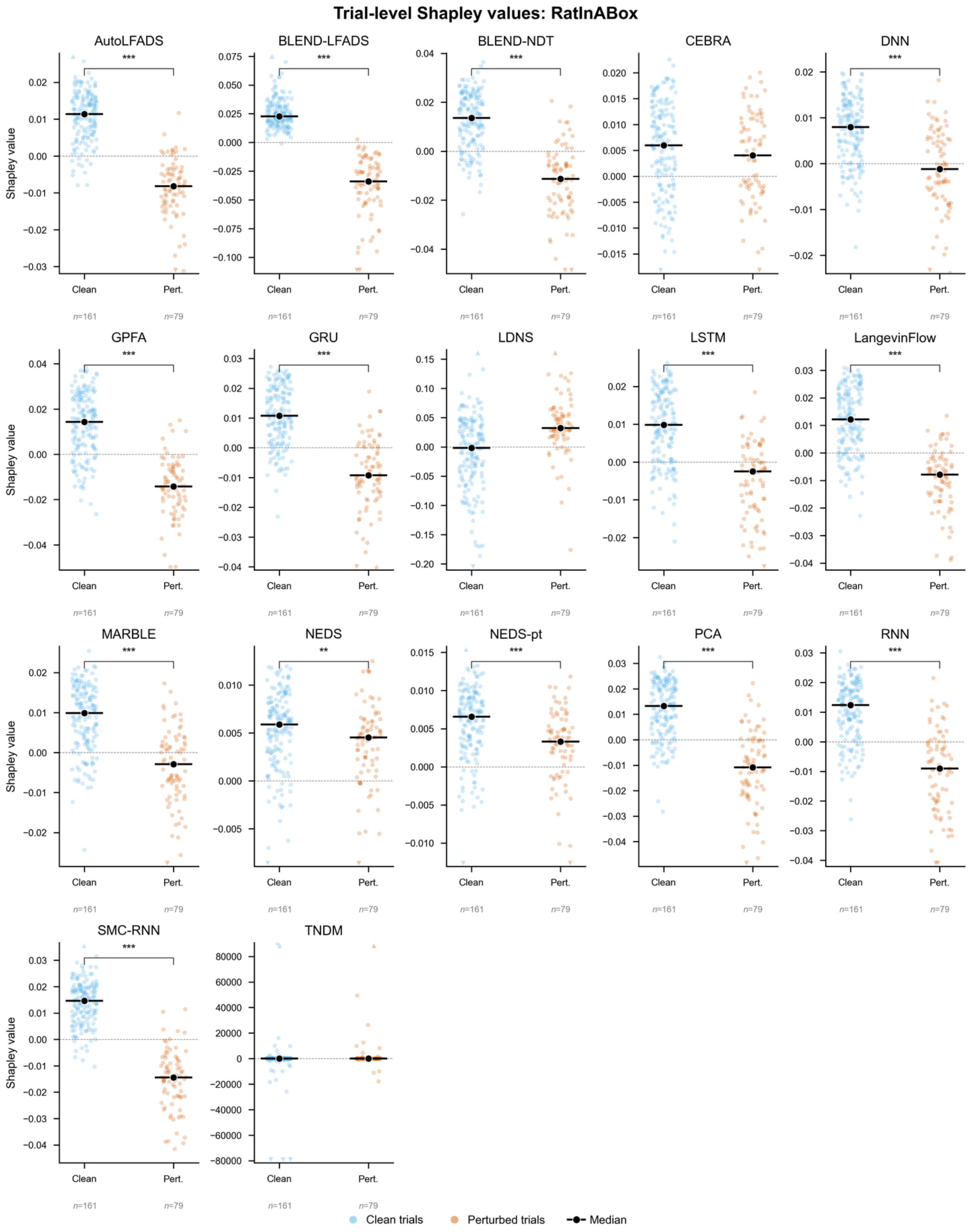


Stars: one-sided Mann–Whitney $U$ test, clean > perturbed (* $P < 0.05$, ** $P < 0.01$, *** $P < 0.001$). Y-limits are panel-specific; triangles mark clipped tails.

**Supplementary Fig. 26 | RatInABox trial-valuation summaries.**

Each decoder subplot shows signed trial values from the 75° full-subspace rotation analysis. Points denote unperturbed training trials (blue; $n$ = 161 per decoder) and trials corrupted by rotating neural population activity while leaving targets unchanged (orange; $n$ = 79 per decoder). Black points and horizontal bars indicate medians. Seventeen evaluated decoders are shown. Vertical limits are set separately for each decoder; triangles at the upper or lower boundary denote observations beyond the displayed range. Asterisks indicate $P$ values from one-sided Mann–Whitney $U$ tests of higher trial values in unperturbed than corrupted trials (*$P < 0.05$; **$P < 0.01$; ***$P < 0.001$).

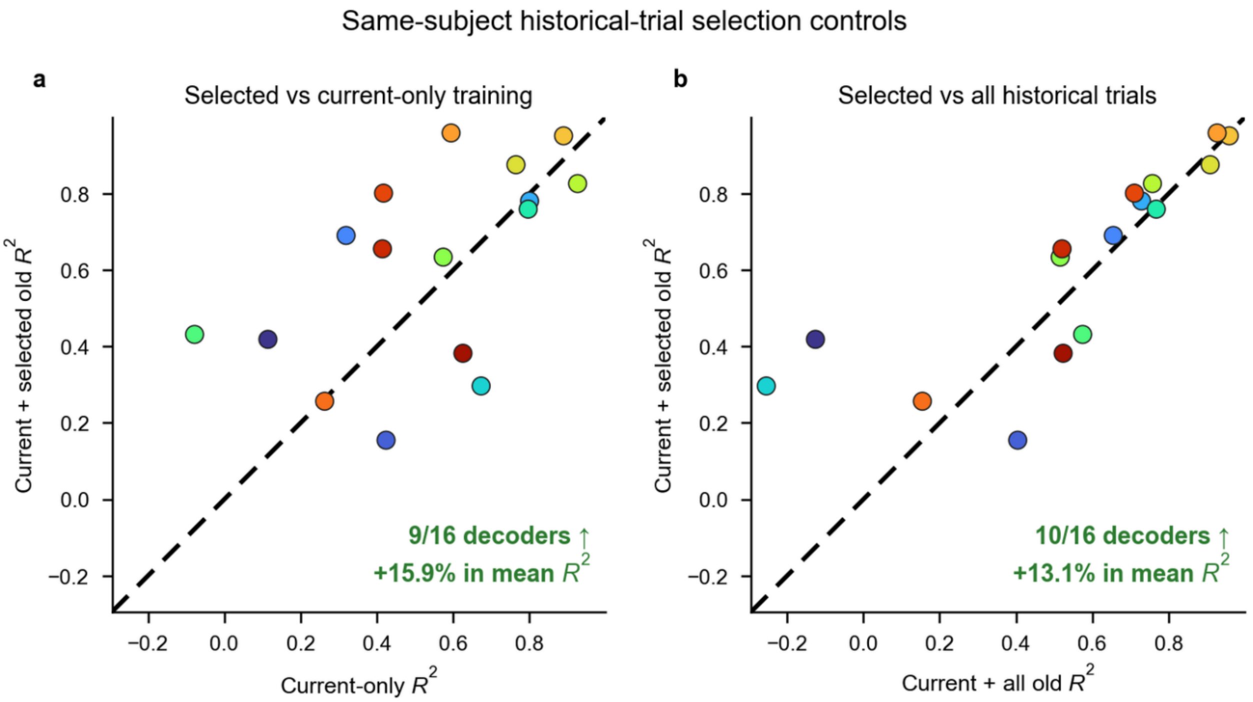


**Supplementary Fig. 27 | Same-subject historical-trial selection controls.**

**a,** Comparison of trial-value-selected historical trials with current-session-only training in the macaque reaching case study. Each point represents one of 16 evaluated decoders using the shared M1 channel feature space; values above the dashed diagonal indicate higher held-out target-session $R^2$ after adding selected historical trials to the current-session training set. **b,** Comparison of trial-value-selected historical trials with all-session pooling. Values above the dashed diagonal indicate higher held-out target-session $R^2$ after selecting historical trials than after adding all available historical trials. In both panels, current-session training trials were retained, trial values were assigned only to candidate trials from earlier sessions and historical trials with non-negative values were selected. Selection improved $R^2$ for 9 of 16 decoders relative to current-session-only training and for 10 of 16 decoders relative to all-session pooling. Mean $R^2$ was 15.9% higher than current-session-only training and 13.1% higher than all-session pooling.
